# Single-cell dissection of prognostic architecture and immunotherapy response in *Helicobacter pylori* infection associated gastric cancer

**DOI:** 10.1101/2024.05.31.596846

**Authors:** Xin Zhang, Guangyu Zhang, Shuli Sang, Yang Fei, Xiaopeng Cao, Wenge Song, Feide Liu, Jinze Che, Haoxia Tao, Hongwei Wang, Lihua Zhang, Yiyan Guan, Shipeng Rong, Lijuan Pei, Sheng Yao, Yanchun Wang, Min Zhang, Chunjie Liu

## Abstract

Most of the gastric cancer (GC) worldwide are ascribed to *Helicobacter pylori* (*H. pylori*) infections, which have a detrimental effect on the immunotherapy’s efficacy. Comprehensively dissecting the key cell players and molecular pathways associated with cancer immunotherapies is critical for developing novel therapeutic strategies against *H. pylori* infection associated GC. We performed a comprehensive single-cell transcriptome analysis of nine GC with current *H. pylori* infection (HpGC), three GC with previous *H. pylori* infection (ex-HpGC), six GC without *H. pylori* infection (non-HpGC), and six healthy controls (HC). We also investigated key cell players and molecular pathways associated with GC immunotherapy outcomes. We revealed the molecular heterogeneity of different cell components in GC including epithelium, immune cells, and cancer-associated fibroblasts (CAFs) at the single-cell level. The malignant epithelium of HpGC exhibited high expression level of inflammatory and epithelial-mesenchymal transition (EMT) signature, HpGC and ex-HpGC were enriched with VEGFA+ angiogenic tumor-associated macrophages (Angio-TAM) and IL11+ inflammatory CAF (iCAF), characterized by high expression levels of NECTIN2 and VEGFA/B. Additionally, we found significant correlations between the abundance of iCAF with Angio-TAM and TIGIT+ suppressive T cells, and iCAF interacted with Angio-TAM through the VEGF and ANGPTL angiogenic pathways. We also developed an immune signature and angiogenic signature and demonstrated that the iCAF abundance and angiogenic signature could predict poor immunotherapy outcomes in GC. We revealed the transcriptome characteristics and heterogeneity of various cellular constituents of HpGC and demonstrated that a synergistic combination of immunotherapy and anti-angiogenic targeted therapy may be an effective therapeutic modality for HpGC.

## Introduction

Gastric cancer (GC) is the fifth most diagnosed tumor and the third most common cause of cancer-causing death. GC is characterized by heterogeneous cellular characteristics and a tumor microenvironment (TME) comprising complex components, posing challenges to personalized therapy (*1*). The development of GC can be attributed to the long-term combined effects of lifestyle, environmental factors, genetic factors, and pathogenic infections. *Helicobacter pylori* (*H. pylori*) infection (*2, 3*) is an important carcinogenic factor, and responsible for approximately 90% of noncardia GC worldwide and approximately 5% of the total burden from all cancers globally (*4*). It also plays a critical role in TME regulation and is linked to response to immunotherapies (*5, 6*). According to the histological examination, serology test, rapid urease test, and *H. pylori* DNA validation, *H. pylori* infection associated GC cases can be classified to current *H. pylori* infection and past *H. pylori* infection, and the *H. pylori* infection status correlated with different molecular characteristics (*7, 8*).

Immune checkpoint inhibitor (ICI)-targeted therapy is an effective anti-cancer treatment for a wide range of human malignancies (*9–12*). However, their efficacy in GC remains controversial (*13*). Previous clinical trial studies such as JAVELIN Gastric 300 (*14*), KEYNOTE-061 (*15*), and KEYNOTE-062 (*16*) have shown that PD-1/PD-L1 inhibitors do not have significant advantages over conventional chemotherapy in the treatment of GC. In contrast, the CheckMate-649 study has revealed that the continued use of nivolumab plus chemotherapy as a standard first-line treatment achieved great success in advanced gastroesophageal adenocarcinoma (*17*). A subgroup analysis has revealed that the Asian population benefits more from immunotherapy than the global population, and among the Asian population, the Chinese population benefits the most from immunotherapy, which could be ascribed to the molecular features leading to GC in Chinese patients, such as *H. pylori* infection (*18, 19*). However, recent studies have demonstrated that *H. pylori* infection hindered the efficacy of immune checkpoint-targeting PD-1/PD-L1 inhibitors or vaccine-based immunotherapies through immune evasion, T cell suppression, and fluctuation in the intestinal flora, compromising the benefits of cancer immunotherapies (*5, 13*). The cellular and molecular heterogeneity of *H. pylori* infection associated GC prevent effective immunotherapies; thus, revealing the discrepancy in the molecular characteristics of various cellular components among GC patients with different *H. pylori* infection status is required for accurate diagnosis and treatment.

The continued availability of single-cell RNA sequencing (scRNA-seq) has allowed systematic profiling of dynamic transcriptional characteristics of thousands of cells simultaneously at the single-cell level with high throughput and low cost, enabling unbiased deciphering of the molecular heterogeneity of various components in biological tissues (*20, 21*). A previous study has applied scRNA-seq technology in various types of gastric mucosa biopsies with different lesions to reveal dynamic transcriptional changes in the cascade of GC and to construct a transcriptional regulatory network in the gastric epithelium (*22*). We have also previously constructed a systematic transcriptomic landscape of gastric adenocarcinoma and profiled the developmental trajectory of GC (*23*). Additionally, scRNA-seq has been used to dissect TME heterogeneity (*24*) and molecular features of various components in the GC TME, including immune cell diversity, which facilitates the understanding of tumor biology and sheds light on novel targets for immunotherapy (*25–27*). However, little attention has been paid to the relationship between *H. pylori* infection, diversity of GC TME, and immunotherapy response at the single-cell level.

This study aimed to reveal the diversity of the *H. pylori* infection associated GC TME and to distinguish key cellular players, molecular pathways, and effector programs associated with response to immunotherapies in GC. We performed unbiased scRNA-seq on gastric tissues of healthy control (HC, n=6), GC without *H. pylori* infection (non-HpGC, n=6), GC with current *H. pylori* infection (HpGC, n=9), and GC with previous *H. pylori* infection (ex-HpGC, n=6) to present the first cellular atlas including a total of 83,637 high-quality cells for deciphering *H. pylori* infection associated transcriptomic architecture. We also profiled the intercellular crosstalk among suppressive T cells, angiogenic tumor-associated macrophages (Angio-TAMs), and inflammatory cancer-associated fibroblasts (iCAFs), which correlated with prognosis and response to immunotherapy. In particular, iCAFs that were linked to Angio-TAM and suppressive T cells exhibited upregulation of NECTIN2 and VEGFA/B (vascular endothelial growth factor A/B), which is indicative of poor immunotherapy efficacy. Moreover, intercellular crosstalk analysis revealed that iCAFs may promote tumor angiogenesis and immune suppression in *H. pylori* infection associated GC, by intercellular interaction with Angio-TAM through the VEGFA/B-VEGFR1 pathway, and TIGIT^+^ suppressive T cells through the NECTIN2-TIGIT pathway, respectively. Our study indicates that the combination of immunotherapy and vascular-targeted therapies could be a potentially efficient approach for the treatment of *H. pylori* infection associated GC patients.

## Results

### Single-cell transcriptomic architecture and molecular features of *H. pylori* infection associated GC

In this study, we first classified the collected GC tissues into non-HpGC, HpGC, and ex-HpGC to accurately evaluate the *H. pylori* infection status of GC patients, which was defined by a combination of serology examinations along with the *H. pylori* DNA assay and histological examination (HE) of each gastric tissue (methods, table 1). The gastric tissues selected were collected from six healthy donors (HC) and six non-HpGC, nine HpGC, and six ex-HpGC patients, who were recently diagnosed and did not receive chemotherapy and radiotherapy before surgery (Fig. 1A). Viable cells with high quality were collected from each gastric tissues for further scRNA-seq using fluorescence-activated cell sorting (FACS). A total of 83,637 cells were retained for subsequent analysis after rigorous quality control, which yielded an average of 1,358 genes and 3,465 transcripts in each cell (fig. S1, A to D). We first employed the inferCNV to evaluate copy number variation (CNV) of autosomal genes among all cells to distinguish malignant and non-malignant cells (fig. S1E), We then identified and annotated nine main cell types according to the expression of canonical gene markers and CNV distribution, which were composed of the non-malignant epithelium (marked with *MUC5AC*, *GKN1*, and *TFF1/2*), malignant epithelium (*KRT17*, *S100A9* and *REG4*), endothelium (marked with *PECAM1*,*VWF* and *PLVAP*), fibroblasts (marked with *DCN*, *LUM* and *COL1A1*), plasma cells (marked with *JCHAIN* and *CD49A*), monocyte/macrophage (marked with *C1QA/B/C* and *IL1B*), mast cells (marked with *CPA3* and *TPSAB1*), B cells (marked with *CD79A* and *MS4A1*), and T cells (marked with *CD2*, *CD3D*, and *CD3E*) (Fig. 1, B to E, fig. S1F, and table S1). To investigate the influence of *H. pylori* infection on the heterogeneity of gastric TME, we analyzed the dynamic cellular component changes of gastric TME in non-HpGC, ex-HpGC and HpGC compared with that of HC. The results revealed a significant increase in the proportion of T cells in GC including non-HpGC, ex-HpGC and HpGC in contrast with HC (Fig. 1, F and G and fig. S2A), which aligned with consensus that tumorigenesis is accompanied by inflammation. In addition, the percentage of non-malignant epithelium significantly decreased in GC compared to normal HC. Whereas, there was a significant increase in the proportion of malignant epithelium in GC compared with that of HC, which is closely related to the process of carcinogenesis. However, there is no significant difference in the proportion of non-malignant epithelium and malignant epithelium among the non-HpGC, ex-HpGC and HpGC (Fig. 1G). To further elucidate the correlation between *H. pylori* infection and the transformation of non-malignant epithelium into malignant epithelium during gastric carcinogenesis, we performed comparative analysis of upregulated genes characteristics between non-malignant epithelial cells and malignant epithelial cells. The result revealed that upregulated genes in the non-malignant epithelium included *MUC5AC*, *GKN1*, and *TFF2*, which participate in secretion of stomach mucus and digestive enzymes, whereas upregulated genes in the malignant epithelium included *TM4SF1*, *IFITM3*, *IGFBP7* and *KRT17* (Fig. 1H and fig. S1, F and G). Gene set enrichment analysis (GSEA) showed that the differentially upregulated genes in the malignant epithelium compared to non-malignant epithelium were enriched in pathways associated with tumor development and progression such as hypoxia, tumor invasiveness, MYC targets, and epithelial-mesenchymal transition (EMT) (Fig. 1I). In brief, our results provided a comprehensive landscape of the cellular and transcriptomic diversity within the cell components in gastric TME associated *H. pylori* infection.

**Fig. 1.**
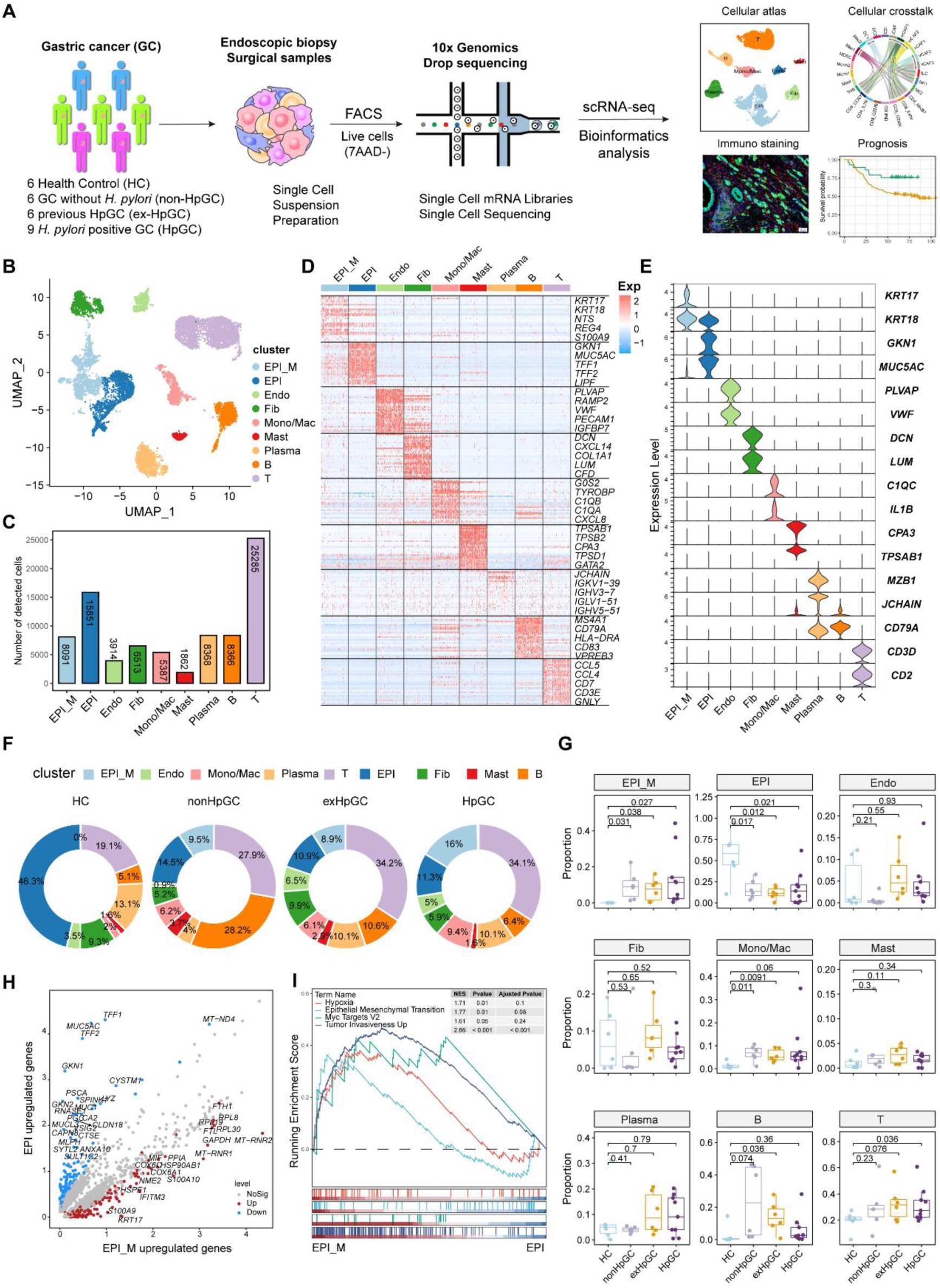
Global analysis of the tumor microenvironment and malignant features of *H. pylori* infection associated GC. (A) A general workflow of GC sample preparation and processing of single-cell suspensions for scRNA-seq analysis. In total, 27 gastric samples, including gastric tissues of healthy control (HC, n=6), gastric cancer without *H. pylori* infection (non-HpGC, n=6), gastric cancer with previous *H. pylori* infection (ex-HpGC, n=6) and gastric cancer with current *H. pylori* infection (HpGC, n=9), were collected to perform scRNA-seq. (B) Uniform Manifold Approximation and Projection (UMAP) plot for unbiased clustering and cell type annotation of 86,637 high-quality cells. EPI_M: malignant epithelium; EPI: non-malignant epithelium; Endo: endothelium; Fib: fibroblast; Mono/Mac: monocyte/macrophage; Plasma: plasma cells; Mast: mast cells; B: B cells; T: T cells. (C) the absolute quantities of nine different cell types. (D) Heatmap showing the top eight differentially expressed genes (DEGs) of nine main cell types. (E) Violin plots showing the expression of signature genes of nine cell types. (F) Pie plot revealing the proportions of nine cell types in HC, non-HpGC, ex-HpGC and HpGC. (G) Box plot showing statistical analysis of proportion of nine cell types in HC, non-HpGC, ex-HpGC and HpGC. The *P* value of Student’s t test is shown. (H) Volcano plot displaying the differential upregulated genes in EPI and EPI_M. (I) GSEA showing the pathway activity (scored per cell by Gene Set Enrichment Analysis) in malignant epithelium (EPI_M) and non-malignant epithelium (EPI).

**Table 1:** Patient characteristics of each sample in GC scRNA-seq.

| Sample | Stage | Age | Sex | Histopathological diagnosis | Site of origin | Lauren's classification | *H. pylori* serum antibody | *H. pylori* DNA | *H. pylori* cagA | *H. pylori* on H&E slide |
| --- | --- | --- | --- | --- | --- | --- | --- | --- | --- | --- |
| exHpGC1 | III | 62 | F | Moderately differentiated adenocarcinoma | Gastric antrum and corpus | Intestinal | + | - | - | - |
| exHpGC2 | III | 61 | M | Moderately differentiated adenocarcinoma | Gastric antrum | Mixed | + | - | - | - |
| exHpGC3 | I | 75 | M | Poorly differentiated adenocarcinoma | Gastric antrum | Diffuse | + | - | - | - |
| exHpGC4 | II | 67 | F | Signet ring cell carcinoma | Gastric body | Diffuse | + | - | - | - |
| exHpGC5 | III | 64 | M | Moderately poorly differentiated adenocarcinoma | Cardia | Intestinal | + | - | - | - |
| exHpGC6 | I | 65 | M | Moderately differentiated adenocarcinoma | Gastric antrum | Intestinal | + | - | - | - |
| HpGC1 | II | 73 | M | Moderately poorly differentiated adenocarcinoma | Stomach angle | Intestinal | + | + | - | + |
| HpGC2 | IV | 55 | M | Poorly differentiated adenocarcinoma | Gastric antrum and corpus | Diffuse | + | + | - | + |
| HpGC3 | I | 49 | F | Moderately-well differentiated adenocarcinoma | Gastric antrum and gastric angle | Intestinal | + | + | - | + |
| HpGC4 | I | 47 | M | Poorly differentiated adenocarcinoma, mostly signet ring cell carcinoma | Gastric antrum | Diffuse | + | + | - | + |
| HpGC5 | II | 67 | F | Signet ring cell carcinoma | Gastric antrum | Diffuse | + | + | - | + |
| HpGC6 | I | 58 | M | Moderately poorly differentiated adenocarcinoma | Greater curvature | Mixed | + | + | + | + |
| HpGC7 | III | 67 | F | Moderately poorly differentiated adenocarcinoma | Angular incisure | Intestinal | + | + | N/A | + |
| HpGC8 | III | 74 | M | Moderately poorly differentiated adenocarcinoma | Greater curvature | Intestinal | + | + | N/A | + |
| HpGC9 | II | 62 | M | Poorly differentiated adenocarcinoma, partial signet ring cell carcinoma | Angular incisure | Mixed | + | + | N/A | + |
| nonHpGC1 | I | 54 | F | Poorly differentiated adenocarcinoma | Cardia | Diffuse | - | - | N/A | - |
| nonHpGC2 | III | 65 | M | Moderately differentiated adenocarcinoma | Cardia | Intestinal | - | - | N/A | - |
| nonHpGC3 | III | 56 | M | Moderately differentiated adenocarcinoma | Cardia | Intestinal | - | - | N/A | - |
| nonHpGC4 | III | 72 | M | Moderately poorly differentiated adenocarcinoma | Angular incisure | Intestinal | - | - | N/A | - |
| nonHpGC5 | I | 56 | M | Poorly differentiated adenocarcinoma | Antrum | Mixed | - | - | N/A | - |
| nonHpGC6 | II | 63 | M | Poorly differentiated adenocarcinoma | Angular incisure | Mixed | - | - | N/A | - |
| HC1 | N/A | 38 | M | Normal | Greater curvature | N/A | - | - | N/A | - |
| HC2 | N/A | 43 | F | Normal | Greater curvature | N/A | - | - | N/A | - |
| HC3 | N/A | 62 | F | Chronic gastritis | Gastric body | N/A | - | - | - | - |
| HC4 | N/A | 61 | F | Normal | Gastric antrum | N/A | - | - | - | - |
| HC5 | N/A | 25 | M | Normal | Gastric antrum | N/A | - | - | - | - |
| HC6 | N/A | 32 | F | Normal | Gastric antrum | N/A | - | - | - | - |

### Cellular characterization of the malignant epithelium of *H. pylori* infection associated GC

To further explore the molecular features of *H. pylori* infection associated GC and the heterogeneity of the malignant epithelium, we performed a sub-clustering analysis of malignant cells and generated six distinct malignant epithelium subpopulations (M1−M6), in which clusters M1 and M2 were mainly derived from non-HpGC, whereas clusters M5 and M6 were mainly originated from *H. pylori* infection associated GC samples including HpGC and ex-HpGC (Fig. 2, A and B and fig. S2, A and B). The expression distribution of these clusters was validated in HpGC by deconvolution analysis using bulk RNA sequencing datasets (Wilcoxon test; fig. S2C) (*28*). We found that malignant epithelium of M1/M2/M3/M4 exhibited high expression levels of epithelium differentiation-related genes, such as *PHGR1* and *KRT20* (Fig. 2C), whereas M5/M6 or *H. pylori* infection associated GC samples exhibited high expression levels of inflammation-related genes, such as *CXCL1/2/3/8*, *S100A9*, *S100P*, and EMT signature, such as *SPARC* and *VIM* (Fig. 2, C and D). In addition, among the *H. pylori* infection associated GC samples, the expression of inflammation-related genes and IgG family members’ genes were much higher in HpGC (Fig. 2D, fig. S2D and table S2). Interestingly, GSEA of the upregulated genes in the HpGC malignant epithelium were mainly associated with EMT, epithelial cell proliferation, and inflammatory response pathways (Fig. 2E). We also found that the malignant epithelium exhibited a high degree of differentiation inter- and intra-heterogeneity (Fig. 2, F and G), the differentiation score of ex-HpGC2 and ex-HpGC5 presented two distinct peaks, one of which is comparable to the peak observed in HpGC, while the another of which resembled the peak observed in non-HpGC sample (Fig. 2F). The finding suggested that a history of *H. pylori* infection in GC can still contribute to the differentiation heterogeneity of malignant epithelium. Furthermore, malignant epithelium from ex-HpGC displayed low differentiation scores, which were even lower in malignant epithelium of HpGC (*P* < 0.05, Wilcoxon test; Fig. 2H). This further substantiate the correlation between the inflammatory and differentiation ability with the severity of *H. pylori* infection. Furthermore, the abundance of malignant epithelium cluster could predict different prognosis outcomes (fig. S2E), and the high differentiation score is highly associated with a better prognosis of GC and vice versa (Fig. 2I).

**Fig. 2.**
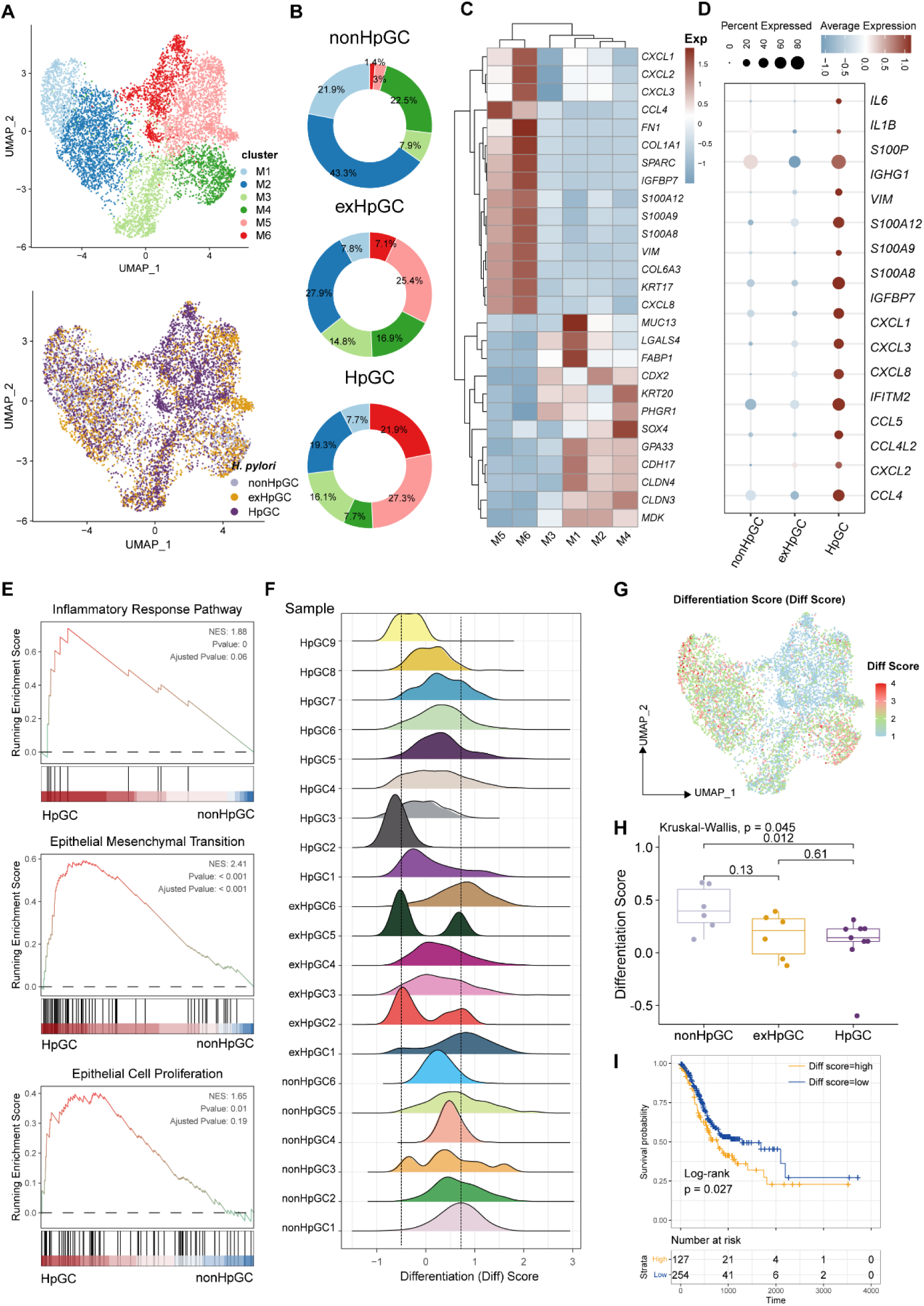
Characterization of the malignant epithelium within GC with different *H. pylori* infection status. (A) UMAP plot showing six subpopulations of malignant epithelium, colored by different cell types (upper) and different *H. pylori* infection status (lower). (B) Pie plot showing the proportion of six subset of malignant epithelium in non-HpGC, ex-HpGC, and HpGC. (C) Heatmap displaying the differentially expressed genes (DEGs) among the six subsets of malignant epithelium. (D) Bubble plot showing the difference of representative molecular among the non-HpGC, ex-HpGC, and HpGC. (E) GSEA showing the pathway activity (scored per cell by Gene Set Enrichment Analysis) in malignant epithelium of HpGC compared to that of nonHpGC. NES, normalized enrichment score. (F) The ridge plot showing the differentiation score (Diff Score) of malignant epithelium within each sample. (G) Heatmap showing the Diff Score of six subpopulations of malignant epithelium. (H) Box plot showing differentiation score among non-HpGC, ex-HpGC, and HpGC, *P*-values were assessed by Wilcoxon test, Two-way ANOVA test is used for comparison of multiple groups. (I) Kaplan-Meier survival analysis of TCGA STAD patients stratified by tumor sample differentiation scores, which was used to group samples into high and low groups based on 33^th^ and 67^th^ percentile. The *P* value of two-sided log-rank test is shown.

Collectively, the findings highlight the transcriptomic heterogeneity of malignant epithelium with different *H. pylori* infection status and the potential utility of the molecular features within the malignant epithelium to distinguish HpGC from non-HpGC, indicative of a high degree of tumor heterogeneity and the influence of the *H. pylori* infection on gastric carcinogenesis and prognosis.

### Cellular characterization of the non-malignant epithelium of *H. pylori* infection associated GC

To determine the cellular phenotype and composition of the gastric mucosa during the progression of *H. pylori* infection, we performed a transcriptomic analysis of the non-malignant epithelium, and these cells were partitioned into nine cell types (Fig. 3A), including chief cells (marked by *PGA3*, and *PGA4*), neck cells (marked by *MUC6*), spasmolytic polypeptide-expressing metaplasia (SPEM, marked by *MUC6* and *TFF2*), intestinal metaplasia (IM, marked by *MUC2*), enterocytes (marked by *APOC3* and *FABP1*), endocrine cells (marked by *CHGA*), pit mucous cell (PMC, marked by *GKN1/2* and *MUC5AC*), pre-PMC (marked by medium expression level of *MUC5AC* and *GKN2*), and parietal cells (marked by *ATP4A*) (Fig. 3, B and C, and table S3). We compared the composition of cells in normal mucosae with that of GC mucosae and observed that the neck cell (*P* = 0.034, Student’s t-test), chief cell (*P* = 0.014, Student’s t-test) and parietal cells (*P* =0.0087, Student’s t-test) decreased dramatically in HpGC compared to HC, whereas the HpGC exhibited increase partition of metaplasia lineage SPEM (*P* = 0.017, Student’s t-test) and intestinal-specific cell types, such as IM (*P* = 0.036, Student’s t-test) and enterocytes (*P* = 0.036, Student’s t-test). Furthermore, HpGC samples had even a higher proportion of metaplasia lineage SPEM and intestinal-specific cell types and a lower proportion of neck cells and parietal cells than that of ex-HpGC and non-HpGC (Fig. 3C). To conform the validity of our findings, we performed deconvolution analysis with independent microarray data (GSE2669) including 124 normal mucosa control (NC), chronic gastritis (CG), intestinal metaplasia (IM), GC associated with *H. pylori* infection samples to decipher the changes in the proportion of cell components with different stages of gastric tissue. Consistent with our scRNA-seq findings, the proportion of SPEM, IM and enterocytes in GC increased gradually during the gastric carcinogenesis process, with even higher proportion in GC compared with that of CG and IM. In contrast, the proportion of chief cells, neck cells and parietal cells decreased significantly in GC, with the lowest proportion in GC compared with that of CG and IM (*P* < 0.05, Student’s t-test; Fig. 3, D and E). These findings further support the validity of our results and indicates that chronic atrophic gastritis induced by *H. pylori* infection is the main cause of premalignant lesions.

**Fig. 3.**
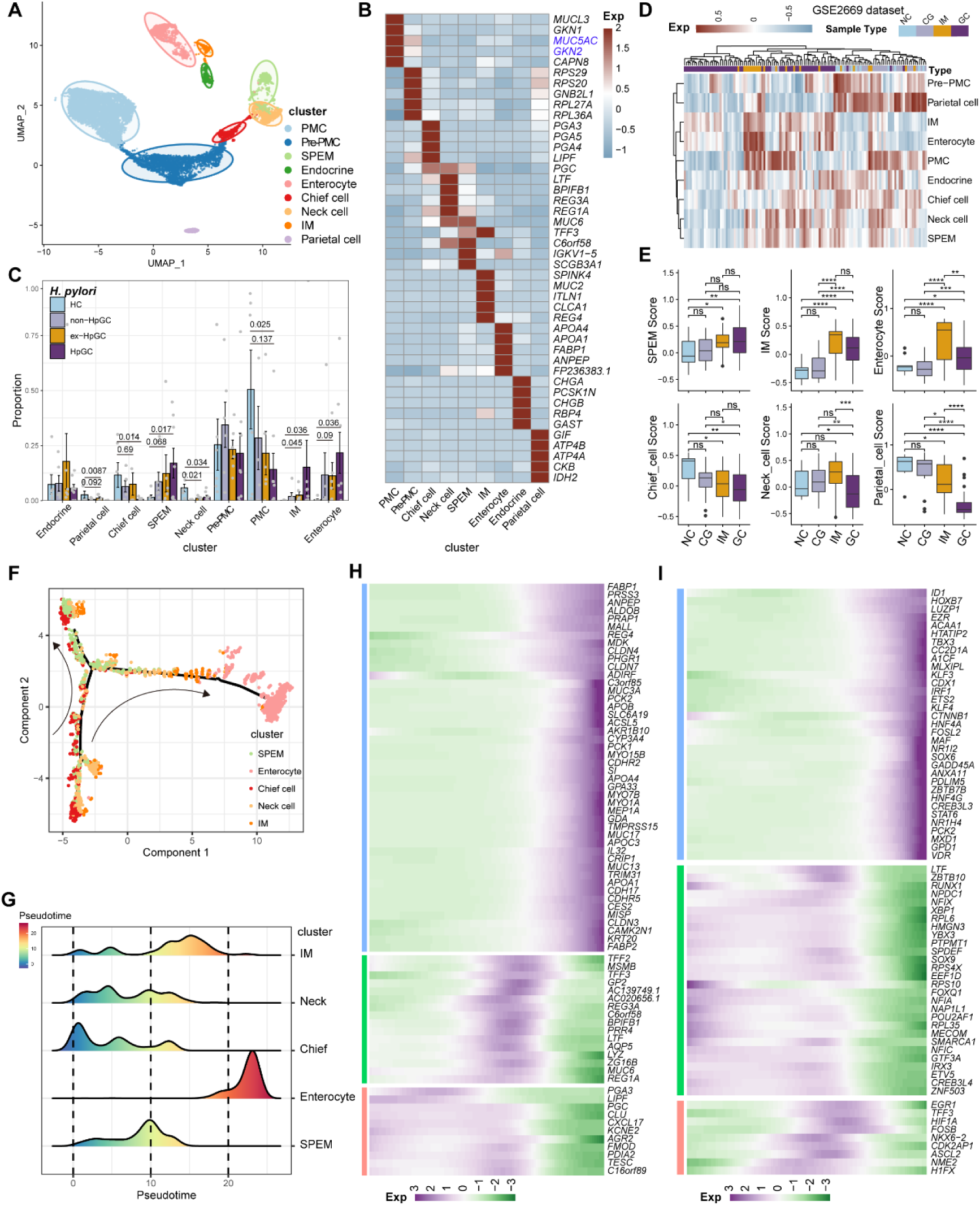
Characterization of the non-malignant epithelium within GC with different *H. pylori* infection status. (A) Unbiased clustering of non-malignant epithelium generated nine subtypes. (B) Heatmap showing the molecular feature of non-malignant epithelium according to the top five DEGs. (C) Box plot showing the dynamic proportion of different cell types in non-malignant epithelium with different *H. pylori* infection status. (D) Heatmap showing the relative abundance (estimated by GSVA) of non-malignant epithelium subtypes in normal control (NC), chronic gastritis (CG), intestinal metaplasia (IM), GC samples (GSE2669). (E) Boxplot showing the relative abundance (estimated by GSVA) of non-malignant epithelium subtypes SPEM, IM, Enterocytes, Chief cells, Neck cells and Parietal cells within NC, CG, IM, and GC samples (GSE2669). The *P* value of Student’s t test is shown. (F) The trajectory analysis showing potential differentiation and transition trajectories in non-malignant epithelium clusters. (G) The ridge plot showing the pseudotime of non-malignant epithelium revealed the gastric pre-lesion process. (H−I) Heatmap showing scaled expression of dynamic genes (G) and TFs (H) along cell pseudotime.

To explore the origin of the premalignant cellular lineage and possible transition mechanism, we performed a single-cell analysis to reveal the developmental trajectories of various cell types in the non-malignant epithelium of GC (Fig. 3F). The results showed that chief cells would differentiate into necks, and then the developmental path would divide these cells into two branches, one of which would develop into SPEM, whereas the other represents SPEM that could further develop into IM and enterocytes through transition states under a persistent inflammatory stimulus (Fig. 3G). In addition, the pattern of pseudotemporal dynamic expression of specific representative genes and transcription factors (TFs) also supports the transition of chief cells to SPEM and IM metaplasia status and finally enterocytes (Fig. 3, H and I).

In brief, our results reveal the transition of GC non-malignant epithelium from chief cells to neck cells, SPEM and IM, and finally to enterocytes, under chronic inflammation at the single-cell level, which may deepen our understanding of the role of *H. pylori* infection in GC.

### Cellular characterization of T lymphoid cells of *H. pylori* infection associated GC

Tumor infiltrating lymphocytes (TILs) are highly heterogeneous lymphocyte subsets. The activation of the different subsets of effector T cells influences the clinical outcome of *H. pylori* infection. Sub-clustering of T lymphoid cells and natural killer (NK) cells generated 11 main cell clusters in HC, non-HpGC, ex-HpGC and HpGC (Fig. 4A), including innate lymphoid cell (ILC, marked by *KIT*, *LST1*, and *ZBTB16*), NK1 (marked by *TYROBP*, *GZMA*, and *CEBPD*), NK2 (marked by *NKG7*, *CCL3*, *GZMH*, and *EOMES*), Treg (marked by *FOXP3*, *BATF*, and *TIGIT*), proliferative CD8_MKI67 (marked by *MK167*, and *CXCL13*), exhausted CD8_CXCL13 (marked by *CXCL13* and *TIGIT*), CD8_GZMK (marked by *GZMK*, *CCL4*, and *YBX3*), effective CD8_IFNG (marked by *IFNG*, *ATF3*), cytotoxic CD8_GZMB (marked by *GZMB*, *KLRC1*, and *MYBL1*), naïve CD4_IL7R (marked by *CCR7*, *SELL*, and *CD40LG*) and CD4_CCR7 (marked by *CCR7*, *SELL*, and *CD40LG*) (Fig. 4, B and C, and table S4). The organization of the T cell compartment showed great differences in GC with different *H. pylori* infection status (Fig. 4D). The exhausted CD8_CXCL13, CD8_MKI67 and Tregs were prominently enriched in ex-HpGC and HpGC samples compared to that in HC (*P* < 0.05, Student’s t-test; Fig. 4E), with a higher percentage in ex-HpGC and HpGC than in non-HpGC. The cytotoxic CD8_GZMB level was significantly decreased in ex-HpGC and HpGC (*P* < 0.05, Student’s t-test; Fig. 4E), which coincides with previous research indicating immunosuppression in the TME. The deconvolution analysis using The Cancer Genome Atlas (TCGA) GC bulk dataset (*29*) indicated that the abundance of Treg and CD8_CXCL13 were elevated in HpGC than non-HpGC (*P* = 0.018, and = 0.12, respectively, Wilcoxon test; Fig. 4F). Next, we calculated the cytotoxic and inhibitory scores across various T lymphoid cell types, revealing that two subtypes of NK cells have the highest cytotoxic score, indicative of apoptosis, and Tregs have the highest inhibitory score, implying immunosuppression (Fig. 4G). Overall survival analysis demonstrated that a high proportion of Tregs and CD8_CXCL13 represent a worse prognosis (*P* = 0.022 and = 0.22, respectively, log-rank test; Fig. 4H). TIGIT, an important immune checkpoint with ligands including PVR and NECTIN2, is highly expressed in various immune cell types, particularly in Tregs within the TME, and is emerging as an immunotherapy target (*30–32*). Interestingly, we also found that the suppressive T cells, including Treg and CD8_CXCL13, interacted closely with the malignant epithelium through TIGIT-PVR/NECTIN2 ligand-receptor pairs, with even stronger interactions between suppressive T cells and malignant epithelium M4 and M6 (Fig. 4I). Interestingly, NECTIN2 and PVR were more enriched in ex-HpGC and HpGC, compared with non-HpGC (Fig. 4J). Immunostaining of HpGC also revealed that suppressive TIGIT^+^ T cells were located close to the pan-CK^+^ NECTIN2^+^ malignant epithelium (Fig. 4K and fig. S3), which supported the transcriptional findings that suppressive T cells could interact with the malignant epithelium through TIGIT–NECTIN2/PVR pairs. Collectively, we found that HpGC and ex-HpGC exhibits high expression levels of NECTIN2, PVR, and its target TIGIT, which may participate in immune escape in GC, indicated that blockade of TIGIT-NECTIN or TIGIT-PVR axes may be a promising immunotherapy modality for *H. pylori* infection associated GC.

**Fig. 4.**
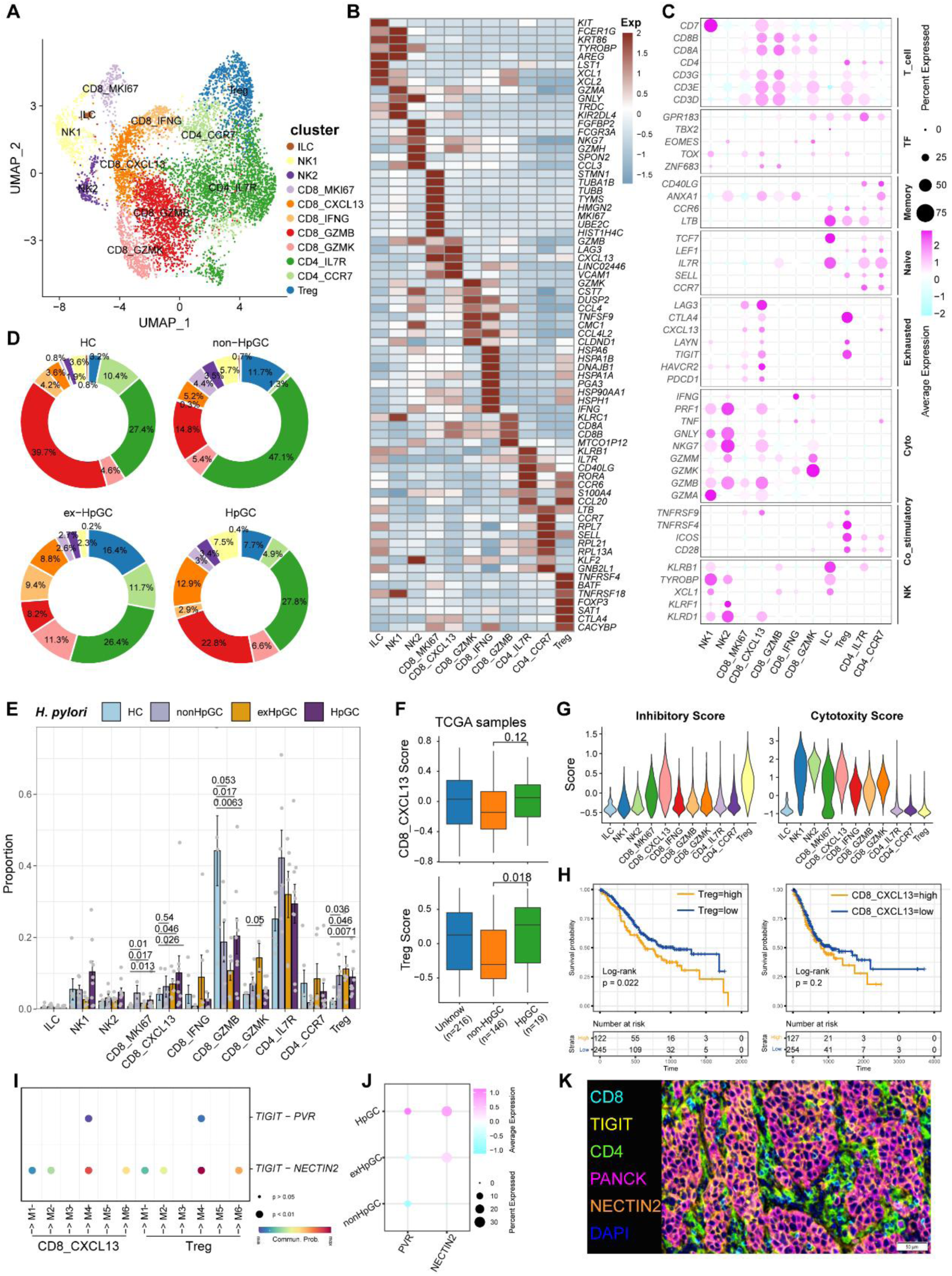
Characterization of tumor-infiltrating T and natural killer (NK) cells in *H. pylori* infection associated GC. (A) Unbiased clustering of T and NK cells generated 11 clusters. (B−C) Molecular features annotations according to the top five DEGs (B), and representative genes (C). (D) Pie plot showing the T/NK cell subtype abundance distribution in HC, non-HpGC, ex-HpGC, and HpGC. (E) The percentage contribution of T/NK cell subtype in HC, non-HpGC, ex-HpGC, and HpGC samples. *P*-values were assessed by Student t test. (F) The deconvolution analysis showing the relative abundance of CD8_CXCL13 and Tregs in GC with different *H. pylori* infection status with TCGA STAD dataset, *P*-values were assessed by Wilcoxon test. (G) The cytotoxic and inhibitory expression scores in T and NK clusters. (H) Kaplan-Meier survival analysis of TCGA STAD patients stratified by CD8_CXCL13 and Tregs relative abundance, which was used to group samples into high and low groups based on 33^th^ and 67^th^ percentile. The *P* value of two-sided log-rank test is shown. (I) Bubble plot showing intercellular interactions between suppressive T cells and malignant cells. (J) Dotplot showing the expression of NECTIN2 and PVR in malignant non-HpGC, ex-HpGC, and HpGC cells. (K) Immunostaining showing the ligand TIGIT expressed in suppressive T cells and the receptor NECTIN2 expressed on the malignant epithelium, respectively, in one HpGC sample.

### Cellular characterization of myeloid cells of *H. pylori* infection associated GC

Myeloid cells are also an important component of tumor-infiltrating immune cells, which play an important role in the regulation of tumor inflammatory responses and angiogenesis. Sub-clustering analysis of various myeloid cells in GC and normal gastric tissues (Fig. 5A) revealed mast cells (marked by *CPA3*), neutrophils (marked by *CXCR2*, *IL1R2*), monocytes (marked by *S100A9*, *EREG*, and *VCAN*), FOLR2_TAM (marked by *LYVE1* and *FOLR2*), TREM2_TAM (marked by *TREM2*), C1QC_TAM (marked by *C1QC*), VEGFA^+^SPP1^+^ Angio-TAM (marked by *VEGFA* and *SPP1*), and three subsets of dendritic cells (DCs, marked by *CD1C*, *LAMP3*, *IDO1*, and *IRF8*) (Fig. 5, B and C, and fig. S4A). The proportion of TREM2_TAMs was significantly increased in HpGC and ex-HpGC, and Angio-TAMs was higher in HpGC tissues than in normal gastric tissues (*P* < 0.05, Student’s t-test; Fig. 5D). TREM2_TAMs exhibited high expression levels of complement genes such as C1QA/C1QB/C1QC, which are predictive factors of prognosis, whereas Angio-TAMs expressed chemokines such as CXCL3, CXCL5, CXCL8, and IL1RN, which are involved in the immune escape (*33*) (Fig. 5E and table S5). Further signature enrichment analysis revealed that Angio-TAMs exhibited high expression levels of angiogenesis signature, whereas TREM2_TAMs exhibited a high level of M2 signature (Fig. 5F). Immunostaining revealed that Angio-TAMs were mainly located in the tumor vasculature, whereas TREM2_TAMs were mainly located in the tumor stroma (Fig. 5G and fig. S5). Furthermore, gene set variation analysis (GSVA) of upregulated genes revealed that TREM2_TAMs were enriched for PD1 signaling, innate immune system, and GC chemosensitivity, whereas Angio-TAMs were enriched for tumor angiogenesis, chemokines, HIF, VEGF pathway, and ECM organization involved in promoting the development of vascular and hypoxic tumors (Fig. 5H). Furthermore, we found that TREM2_TAMs and Angio-TAM abundance were both highly correlated with Treg abundance (Fig. 5I and fig. S4B), which supports the notion that TAMs play crucial roles in the modulation of Treg maturation and recruitment. Interestingly, the intercellular crosstalk analysis revealed different ligand-receptor axes between TREM2_TAM-Tregs and Angio-TAM-Tregs. The SPP1-CD44 axis and the “do not eat me” axis THBS1-CD47, which is an immune checkpoint controlling CD8^+^ T cell activation and immune escape, were mainly enriched in Angio-TAM and Tregs. Additionally, we found Angio-TAM derived from HpGC or exHpGC showed higher expression of *THBS1*, *VEGFA*, and *SPP1* than Angio-TAM of non-HpGC, and HC (fig. S4D). The immune checkpoint axes NECTIN2-TIGIT, CD86-CD28, CD86-CTLA4, CXCL16-CXCR6, and LGALS9-HAVCR2 (TIM3) were exclusively enriched in TREM2_TAM and Tregs (Fig. 5J and fig. S4C). Furthermore, the deconvolution analysis in TCGA bulk GC transcriptome dataset revealed that the enrichment of Angio-TAMs was more significant in HpGC compared with non-HpGC (*P* = 0.047, Wilcoxon test; Fig. 5K) and was associated with poor prognosis of GC (*P* = 0.0035, log-rank test; Fig. 5L). Overall, our results revealed the heterogeneity and function differences of myeloid cells of GC with different *H. pylori* infection status, and decipher distinct roles of TREM2_TAM and Angio-TAM involved in gastric carcinogenesis and prognosis.

**Fig. 5.**
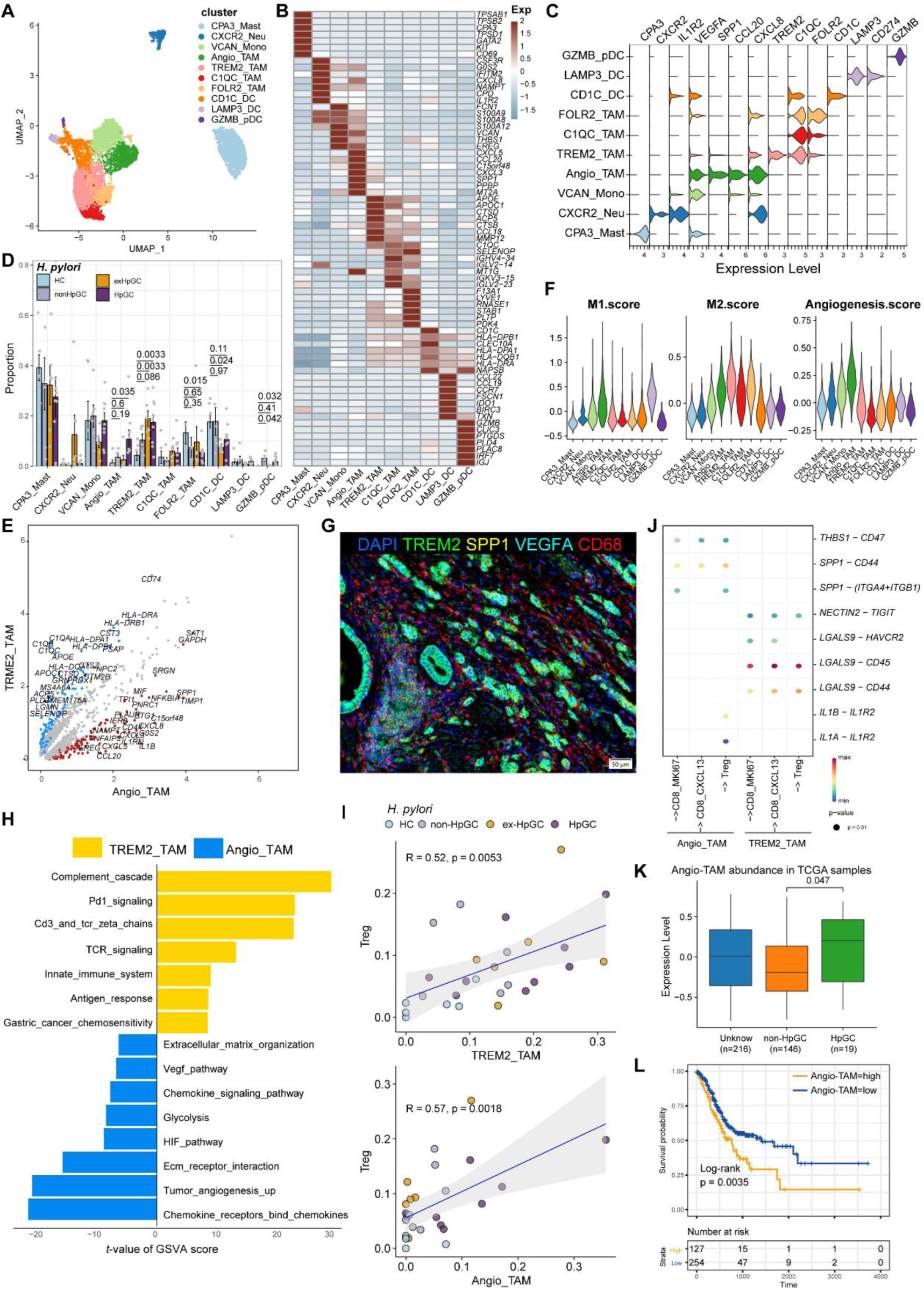
Characterization of tumor-infiltrating myeloid cells by scRNA-seq in *H. pylori* infection associated GC. (A–C) Unbiased clustering of myeloid cells generated nine clusters (A), and molecular features were annotated according to the top seven DEGs (B) and representative genes (C). (D) The percentage contribution of each myeloid cells cluster in HC, non-HpGC, ex-HpGC, and HpGC. *P*-values were assessed by Student t test. (E) Volcano plot showing the DEGs between Angio-TAM and TREM2^+^ TAM. (F) Violin plot showing the expression of functional gene sets in myeloid clusters. (G) Immunostaining showing the distribution of Angio-TAM and TREM2^+^ TAM in one HpGC sample. (H) Barplot showing the enriched signaling pathway between Angio-TAM and TREM2^+^ TAM. (I) The correlation of cell type (percentage) between Tregs and Angio-TAM and TREM2^+^ TAM. (J) Dotplot showing intercellular interactions among suppressive T cell and Angio-TAM and TREM2^+^ TAM. (K) The relative abundance of Angio-TAM in HpGC and non-HpGC in TCGA STAD dataset, *P*-values were assessed by Wilcoxon test. (L) Kaplan-Meier survival analysis of TCGA STAD patients stratified by Angio-TAM relative abundance, which was used to group samples into high and low groups based on 33^th^ and 67^th^ percentile. The *P* value of two-sided log-rank test is shown.

### Transcriptomic diversity of cancer-associated fibroblasts in *H. pylori* infection associated GC

Cancer-associated fibroblasts (CAFs), another major component of the TME, not only produce ECM but also secrete various molecules, including cytokines, chemokines, and growth factors, which are correlated with migration, infiltration, and EMT, thereby playing a role in GC development and resistance to immunotherapy (*34*). Therefore, we focused on the molecular characteristics of CAFs. CAFs can be divided into six subpopulations, including three subsets of vascular-related CAF (vCAF) featuring the expression of microvasculature genes MCAM, RGS5, and MYH11; and two subsets of matrix CAF (mCAF) characterized by high expression of extracellular matrix (ECM) genes including *PDGFRA*, *LUM*, and *DCN* and inflammatory CAF (iCAF), which was identified by high expression of *TWIST1*, *VEGFA* and inflammatory chemokine genes *CXCL5/8*, *IL11* (*35*), and *IL24* (Fig. 6, A to C, fig. S6A, and table S6). Interestingly, iCAF shared highly consistent transcriptome phenotype of iCAFs derived from human intrahepatic cholangiocarcinoma (fig. S6B) (*36*). The organization of the CAF compartment showed great differences in GC with different *H. pylori* infection status (Fig. 6D) and the ratio of iCAF was significantly higher in HpGC tissues than in normal gastric tissues (*P* = 0.023, Student’s t-test; Fig. 6E) and the ex-HpGC also showed an increase trend (*P* = 0.083, Student’s t-test; Fig. 6E). Further enrichment analysis revealed that iCAF enriched chemokines, TGFβ, ECM, IL6 pathway, PDGFRA pathway, inflammatory response, hypoxia, tumor angiogenesis, and VEGF signaling pathway (Fig. 6F), which are crucial for tumor progression, metastasis, and immune escape (*35*). Additionally, the abundance of iCAF significantly correlated with Angio-TAM and suppressive Treg cells (Fig. 6G), the above results indicated an important role of iCAF in tumor TME regulation and immune escape.

**Fig. 6.**
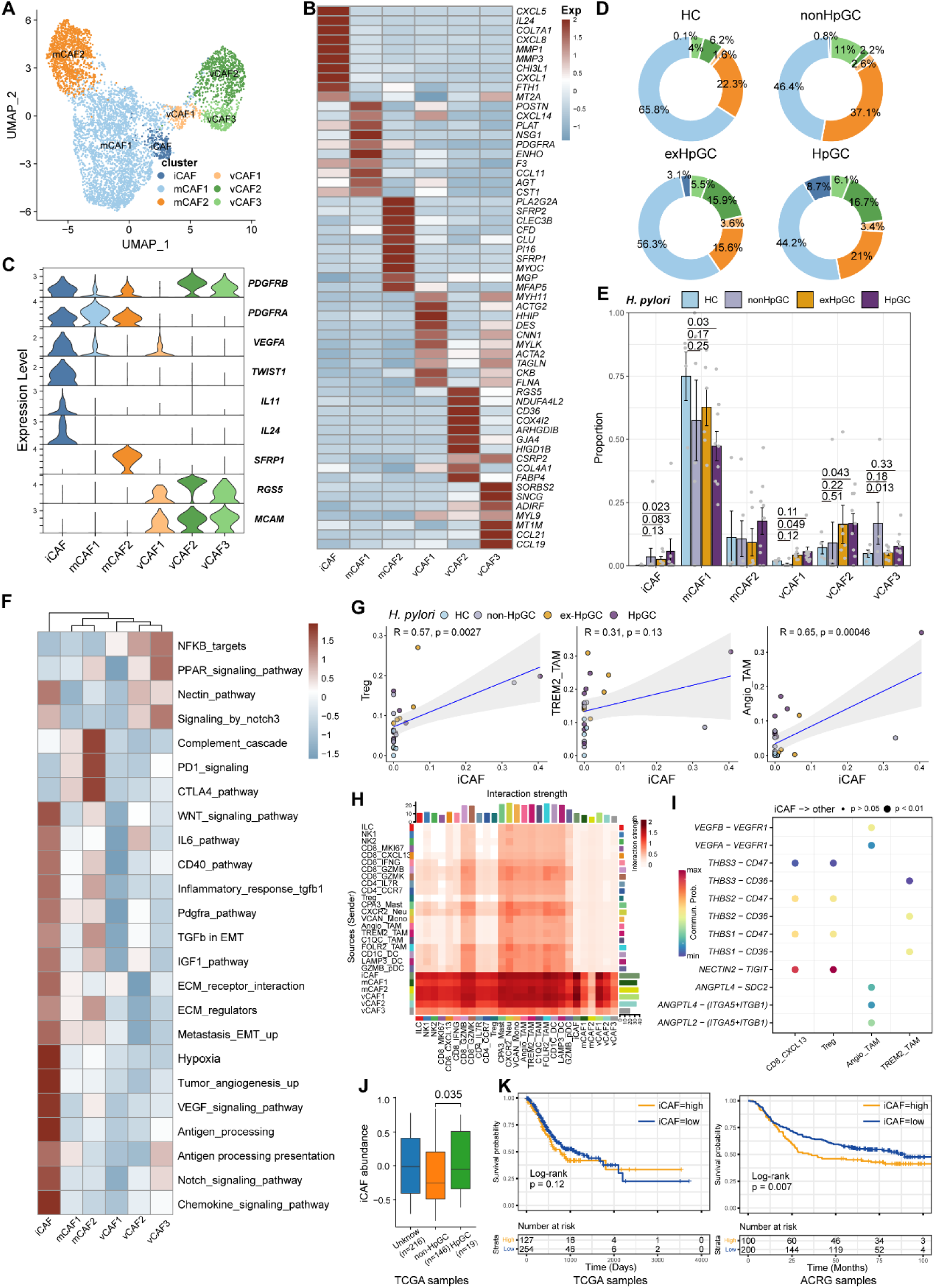
Characterization of cancer-associated fibroblasts (CAFs) by scRNA-seq in *H. pylor* infection associated GC. (A) Unbiased clustering of CAF generated 6 clusters. (B–C) Molecular features annotations according to the top ten DEGs (B) and representative genes (C). (D) The pie plot showing the abundance distribution of six CAF subset in HC, non-HpGC, ex-HpGC, and HpGC. (E) The percentage contribution of each CAF cluster in HC, non-HpGC, ex-HpGC, and HpGC. *P*-values were assessed by Student t test. (F) Heatmap showing the enriched signaling pathway among six CAF clusters. (G) The cell type percentage correlation between iCAF and Treg, Angio-TAM, and TREM2^+^ TAM. (H) Heatmap showing intercellular interaction strength among immune cells and different subsets of CAF. (I) Enriched signaling pathway among suppressive T cell, Angio-TAM, and TREM2^+^ TAM with iCAF. (J) The relative abundance of iCAF in HpGC and non-HpGC in TCGA STAD dataset. *P*-values were assessed by Wilcoxon test. (K) Kaplan-Meier plot shows that the abundance of iCAF predicts poor prognosis of GC using two public bulk RNA sequencing dataset (left, TCGA; right, ACRG). The abundance of iCAF was used to group samples into high and low groups based on 33^th^ and 67^th^ percentile. The *P* value of two-sided log-rank test is shown. ACRG: Asian Cancer Research Group.

Further intercellular crosstalk analysis showed that iCAF had the strongest interaction with suppressive T cells and TAM, indicative of complex roles in the TME (Fig. 6H). Specifically, the THBS, NECTIN, TIGIT, ANGPTL, and VEGF signaling pathways were enriched in the cellular interplay between iCAF and suppressive T cells and TAM (Fig. 6I). Detailed ligand-receptor crosstalk analysis showed that iCAF interacts with CD8_CXCL13, Tregs mainly by NECTIN2-TIGIT ligand-receptor pair (Fig. 6I), which strongly implied that *H. pylori* infection upregulates the expression of NECTIN2 in stromal cells that competitively bind to the TIGIT receptor on T cells, leading to inhibition of T cell responses and immune escape of GC. In addition, angiogenesis-associated ligand-receptor pairs such as VEGFA/B-VEGFR1 and ANGPTL4-SDC2 were mainly enriched between iCAF and Angio-TAMs (Fig. 6I). Interestingly, we found iCAF derived from HpGC showed higher expression of *NECTIN2*, *VEGFA*, *PVR* and inflammatory chemokine genes *IL11* and *IL24* compared to ex-HpGC, non-HpGC, and HC. Additionally, we found that the elevated expression level of *IL11*, *VEGFA*, *IL24*, and *TWIST1* was correlated with poor prognosis in GC (*P =* 0.14, *P <* 0.05, *P <* 0.05 and *P <* 0.05, respectively, log-rank test; fig. S6, C and D). Furthermore, the validation results by virtue of two public GC bulk RNA datasets (*28, 29*) revealed that the abundance of iCAF was elevated in HpGC samples than non-HpGC (*P* = 0.035, Wilcoxon test; Fig. 6J) and was highly associated with poor prognosis in two public bulk transcriptomic dataset (*P =* 0.12 and = 0.007, respectively, log-rank test; Fig. 6K). The above results indicated that iCAFs promoted tumor angiogenesis and immune suppression in *H. pylori* infection associated GC, by upregulation of VEGFA/B-VEGFR1 pathway, and NECTIN2-TIGIT pathway.

### The association of immune and stromal composition abundance with GC immunotherapy

*H. pylori* infection has multiple immunomodulatory effects on the host, which can not only activate the immune response but also negatively regulate it, causing immune escape. However, the key cell players and molecular features for predicting the GC immunotherapy response remains largely unknown. To reveal the detailed molecular features of the *H. pylori* infection associated GC TME for predicting the outcomes of immunotherapy, we performed the devolution analysis to evaluate the cell type abundance, immune checkpoint, and angiogenic signature expression in GC immunotherapy-treated bulk RNA sequencing dataset (*37*). GSEA and devolution analysis demonstrated a high abundance of iCAF and Angio-TAM in anti-PD-1 immunotherapy non-responsive (NR) patients, while CD8_CXCL13, CD8_GZMB, CD8_IFNG and CD8_MKI67 were more abundant in anti-PD-1 responsive cases (Fig. 7A and *P* < 0.05, Student’s t-test; fig. S7A). The relative abundance of several cell clusters such as CD8_CXCL13, CD8_MKI67 had a remarkable correlation with improved anti-PD-1 overall survival (OS) and progression-free survival (PFS), while the abundance of iCAF and Angio-TAM was significantly associated with poor survival (Fig. 7, B and C). Furthermore, we defined an anti-PD-1 immune signature and an angiogenic signature (Fig. 7D and fig. S7B) derived from the cell type signatures of scRNA-seq result and found that the immune signature was highly correlated with CD8_GZMK and CD8_GZMB, while the angiogenic signature highly correlated with iCAF and Angio-TAM (fig. S7C). Interestingly, we found that the expression of immune signature was higher in immunotherapy responsive GC cases than non-responsive cases, while the angiogenic signature showed the opposite trend (*P* < 0.05, Wilcoxon test; Fig. 7E). To better understand the cellular and molecular characteristics that react to immunotherapy, we constructed a comprehensive model comprising cellular composition and signature to evaluate the immunotherapy response of different parameters (fig. S7, D to F). The results showed that immune signatures such as *CXCL13*, *LAG3*, *TIGIT*, and *PDCD1*, and cell types such as CD8_CXCL13 and TREM2_TAM had high diagnostic power to distinguish between anti-PD-1 responsive cases and non-responsive cases (Area under the curve (AUC) > 0.75). Additionally, the immune signature we defined also effectively distinguished anti-PD-1 responsive cases from non-responsive cases to achieve better anti-PD-1 immunotherapy prognosis (AUC = 0.755, Fig. 7F), whereas the angiogenic signature had poor diagnostic ability for identifying anti-PD-1 responses as well as poor anti-PD-1 immunotherapy prognosis (AUC = 0.367, Fig. 7F). Further Kaplan-Meier survival analysis revealed that the defined immune signature was correlated with better immunotherapy efficacy in terms of OS and PFS in GC while the angiogenic signature linked with poor immunotherapy response (*P* < 0.05, log-rank test; Fig. 7G and fig. S7G). In brief, by integrating our single-cell transcriptional results of GC with public bulk RNA-seq data of immunotherapy-treated GC, we identified several cell types and molecules that could serve as indicators to predict the immunotherapy response of GC, thus flagging up individualized GC therapy.

**Fig. 7.**
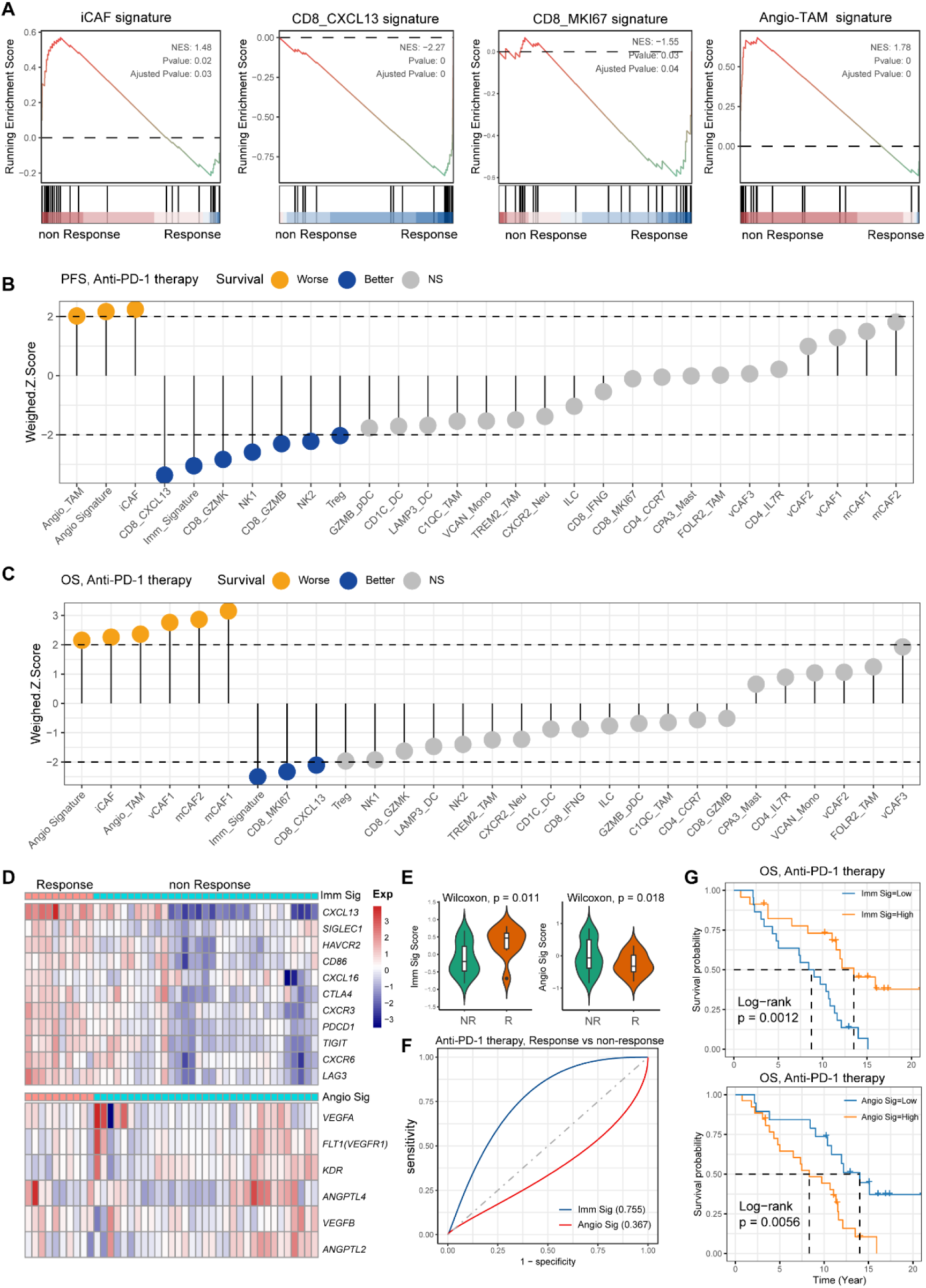
Single cell TME composition associated with GC immunotherapy outcome. (A) GSEA plot showing the enrichment of iCAF, CD8_MKI67, Angio-TAM, and CD8_CXCL13 in anti-PD-1 responsive or non-responsive GC. NES, normalized enrichment score. (B–C) Bar chart showing the cell subtypes relative abundance, the immune signature, angiogenesis signature derived from scRNA-seq predicted GC immunotherapy outcome, progression-free survival (B), and overall survival (C). (D) Heatmap showing the expression of immune and angiogenesis signature derived from scRNA-seq in immunotherapy responsive and non-responsive GC. (E) Violin plot showing the expression of immune signature and angiogenesis signature in immunotherapy responsive and non-responsive GC. The *P* value of Wilcoxon test is shown. (F) A model for evaluating the GC immunotherapy sensitivity and specificity using the immune signature and angiogenesis signature derived from scRNA-seq. (G) The Kaplan-Meier plot showing the immune signature and angiogenesis signature could efficiently predict prognosis of GC anti-PD-1 therapy. The *P* value of two-sided log-rank test is shown.

## DISCUSSION

GC is the third most common tumor in Asia and is mainly caused by *H. pylori* infection. Numerous studies had shown that *H. pylori* infection not only led to GC but also affected the TME of GC and response to immunotherapy (*5, 13, 38, 39*). Owing to advances in scRNA-seq, the heterogeneity of cell components in the TME of human malignancies can be elucidated. In this study, we profiled the single-cell transcriptional characteristics of the TME of GC tissues to reveal the role of *H. pylori* infection in GC development, TME regulation, and immunotherapy response. Specifically, we revealed the TME transcriptomic phenotype and composition, functional features, and potential intercellular interactions of GC with different *H. pylori* infection status, which provided evidence and biomarkers for predicting the prognosis of immunotherapies for *H. pylori* infection associated GC.

First, we used inferCNV algorithms to distinguish non-malignant epithelium from the malignant epithelium and found that malignant epithelium derived from *H. pylori* associated GC exhibited inflammation and EMT signatures, which provided potential biomarkers for the diagnosis of *H. pylori* infection associated GC (*40*). We also revealed that *H. pylori* infection leads to dramatic loss or death of parietal cells and delineated the development trajectory of gastric chief cells along the cascade GC at the single-cell level, which broadened our previous findings to reveal the chief cell transitions (*23*). Based on the expression pattern of signature genes and TFs, we found that chief cells could give rise to neck cells; subsequently, SPEM then showed distinct branches, and only part of SPEM developed into intestinal-specific cell types, including IM and enterocytes, under conditions of chronic inflammation, which further validates the evolutionary path of the non-malignant epithelium of GC under *H. pylori* infection.

Next, we evaluated the heterogeneity and molecular features of immune and stromal components among the TME of normal and all GC tissue and found that HpGC tissues were enriched with iCAF, Angio-TAM, TREM2^+^ TAM and suppressive T cells, and that iCAF was highly linked to Angio-TAM and suppressive T cells, also the abundance of iCAF is associated with poor immunotherapy efficacy. Intercellular crosstalk analysis revealed that iCAF and TREM2^+^TAM interacted with suppressive T cells: Treg and CXCL13^+^CD8 via the NECTIN2-TIGIT pathway. Previous studies had shown that *H. pylori* infection upregulates PD-L1 expression in a variety of cells, including TAM, eosinophils, and dendritic cells (*13, 19, 41*). PD-L1 upregulation in these cells may compete with PD-1 binding to T cells, thereby inhibiting T cell responses, and affecting the therapeutic efficacy of PD-1/PD-L1 inhibitors (*42, 43*). The above results indicated that *H. pylori* infection could upregulate NECTIN2 expression in immune and stromal components, which mediated immune escape and impairs immunotherapy efficacy. Hence, inhibition of the TIGIT-NECTIN2 axis may be a promising treatment strategy in HpGC.

*H. pylori* infection impairs the efficacy of anti-PD-1/CTLA4 immunotherapies or anti-cancer vaccines, and a lack of efficient targets impedes the treatment of *H. pylori* infection associated GC (*5*). However, the key cellular players and signaling pathways involved in the association of *H. pylori* infection with immunotherapy efficacy remain poorly investigated. In this study, iCAFs exhibited upregulation of angiogenic molecules VEGFA/B and ANGPTL, with the highest expression levels identified in cells derived from HpGC. Additionally, iCAFs were notably enriched in tumor angiogenesis, ECM, TGFβ, and VEGF pathway, our results also showed that iCAFs could interact not only with immunosuppressive T cells through the NECTIN2-TIGIT and THBS1/2-CD47 axes, but also with Angio-TAMs through the VEGFA/B-VEGFR1 and ANGPTL-SDC2 axes. These results indicated an important role of iCAFs in stromal TME regulation, immune escape, and tumor angiogenesis. Recent research displayed that block of placental growth factor (PIGF), a member of the vascular endothelial growth factor, could inactive CAFs and reduce the deposition of fibrosis-associated collagen and desmoplasia in pancreatic cancer TME, which could improve survival rate of murine pancreatic ductal adenocarcinoma (PDAC) model. Furthermore, PD-L1 directed PIGF/VEGF block by fusing of single chain of anti-PD-L1 antibody (atezolizumab) to VEGF-Grab could target PD-L1-expressing CAF thus promote anti-fibrotic and anti-tumor effect (*44*). As we know, CAFs can shape the ECM, and form a barrier for drug or therapeutic immune cell penetration, which prevents the deep penetration of drugs and immune cells into tumor tissues, thereby reducing the effectiveness of treatments. Therefore, inhibition of tumors by modulating CAFs or overcoming their barrier effects is a new therapeutic strategy. Taken together, our findings were consistent with those of previous ones that had hypothesized that anti-angiogenic targeted therapy could augment antitumoral immunity and enhance immunotherapy efficacy by targeting tumor angiogenesis, exhausted T cells proliferation and myeloid cell inflammation(*45–48*).

This study has some limitations and the main findings should be validated in a large patient cohort. Secondly, our findings were mainly based on scRNA-seq and bioinformatic analysis, only a small number of key molecules and cell types have been verified by immunostaining, additionally, we conducted deconvolution analysis using external datasets to further validate our results. Even these validations provide support at genomic and protein level, it is important to note that functional validation is still lacking. Further clinical trial research should be performed to validate the efficacy of combination immunotherapy alone or in combination with anti-VEGF therapy in *H. pylori* infection associated GC. Nevertheless, we provided vital insights into the molecular and cellular heterogeneity of *H. pylori* associated GC at a single-cell level. Moreover, the enrichment of the TIGIT-NECTIN immune checkpoint and angiogenesis signaling pathways in different cell types in the TME of *H. pylori* infection associated GC implies that the potential combination of immunotherapies and anti-angiogenic targeted therapeutic modalities could be an effective therapeutic approach for *H. pylori* infection associated GC.

## Materials and methods

### Ethics approval

The study was conducted according to the guidelines of the Declaration of Helsinki and approved by the Ethics Committee of Fourth Medical Center of PLA General Hospital. (2021KY011-HS001, 2021KY041-HS001) All study participants provided written informed consent.

### Acquisition of fresh tissue materials and preparation of single cell suspension

Normal gastric tissue biopsy specimens were obtained from 2–3 gastric antrum sites through conventional upper gastrointestinal endoscopy using biopsy forceps. GC tissues were obtained immediately after surgical resection with a scalpel. The fresh tissue samples were then washed with phosphate-buffered saline (PBS) and divided into two parts. One part was processed into a single cell suspension, and the second part was used for other experiments, including the detection of *H. pylori* DNA, H&E and immunofluorescence staining. The patients’ serum was used to detect the IgG antibody of *H. pylori*. Patients with positive serum antibody and genomic DNA of *H. pylori*, and *H*.

*pylori* observed on H&E slides, were defined as GC patients with current *H. pylori* infection (HpGC). Patients with positive serum antibody but negative genomic DNA had a history of *H. pylori* eradication or antibiotic use before surgery, and *H. pylori* cannot be observed on H&E slides. They were defined as GC patients with previous *H. pylori* infection (ex-HpGC). The negative detection of serum antibody and genomic DNA, as well as H&E, indicated that the volunteers never had *H. pylori* infection, which was defined as GC patients or healthy volunteers without *H. pylori* infection (non-HpGC and HC).

The biopsy and surgical samples used to prepare the single cell suspension were immediately put into a tissue preservation solution (Miltenyi Biotec, Germany) and transported to the field laboratory in an ice bath for immediate preparation of the single cell suspension. Fresh tissue samples were washed with 4 °C precooled Dulbecco’s Phosphate-Buffered Salines (DPBS, Solarbio, Beijing) for 2–3 times, cut into small pieces with surgical scissors, and then transferred to a 1.5 ml centrifuge tube. The tissue samples were incubated in a shaker at 37 °C for 30–50 min with an in-house prepared enzymolysis solution (1 mg/ml type IV collagenase (Solarbio, Beijing) + 10 U/μL DNase I DNase I (Roches)) or MACS Human Tumor Dissociation kit (DS_130-095-929, Miltenyi Biotec, Germany). The incubation was terminated when the digestive fluid turned turbid and the tissue block disappeared. The cell suspension was filtered with a 40-µm cell sieve and centrifuged at 4 °C at 300 r/min for 5 min. After the supernatant was discarded, the cells were resuspended in 1 ml of DPBS, and 3 ml of precooled red blood cell lysate (Solarbio, Beijing) were added. The cells were evenly aspirated, incubated at 4 °C for 5–10 min, and centrifuged again. The cells were stained with 7-Aminoactinomycin D (7-AAD, eBiocience, Cat# 00-6993-50) staining solution (100 μL 1% BSA/PBS + 5 μL 7-AAD) at 25 °C for 5 mins. Consequently, the individual cells with high quality were sorted by FACS performed with a BD Aira II instrument and checked by staining them with trypan blue under the microscope for single cell transcriptomic library construction.

### Detection of *H. pylori* nucleic acid in the gastric mucosa

*H. pylori* in human gastric mucosa was detected using a *H. pylori* nucleic acid detection kit (PCR-fluorescent probe method, Daan Gene, Guangzhou, China), according to the manufacturer’s protocols. Briefly, the following steps were performed: (1) DNA extraction: DNA extraction solution was added to the negative/positive quality controls and the samples to be tested. The solutions were treated at a constant temperature of 100 °C for 10 min, and centrifuged at 12,000 r/min for 5 min for later use; (2) Fluorescence PCR: 2 μL samples were briefly centrifuged at 8,000 r/min and then placed in a PCR machine, 50°C for 8 min, 93°C for 2 min, 93°C for 45 s→55°C for 1 min (10 cycles), 93°C for 30 s→55°C for 45 s (30 cycles), and 50°C for 45 s to collect fluorescence values. The system automatically analyzed the results to calculate the threshold cycle value (CT value); (3) Quality control and results calculation: the fluorescence signal of the negative control material did not increase, the growth curve of the positive control material was S-shaped, and the Ct value was ≤27.04. The sample results were compared to those of the quality control material. *H. pylori* cagA was detected with the Nucleic acid test kit of *H. pylori* type I (fluorescent PCR method, Beijing Xinji Yongkang, China), and the method was carried out according to the manufacturer’s protocols.

### Detection of serum antibody against *H. pylori*

*H. pylori* IgG ELISA kit (IBL, Germany) was used to detect the presence of anti-*H. pylori* antibodies in the serum of the subjects following the manufacturer’s instructions. The cut-off index (COI) was determined by dividing the optical density (OD) value of the sample by the OD value of cut-off standard. The cut-off index (COI) was determined by dividing the optical density (OD) at 450nm value of the sample by the OD_450_ value of cut-off standard. The serum was considered positive for *H. pylori* if the COI was > 1.2, while the serum was considered negative if the COI < 0.8.

### Single-cell sequencing and pre-processing data

We prepared single-cell RNA-seq libraries on the Chromium platform (10× Genomics, Pleasanton, CA, USA) using the Chromium Next GEM Single-Cell 3’ Kit v2 following the manufacturer’s protocol to generate a complementary deoxyribonucleic acid (cDNA) library in Biomarker Technologies and Capitalbio Technology Corporation (China). Briefly, viable cells (7-AAD negative) with high quality were pooled together and washed thrice with RPMI-1640, concentrated to 700−1000 cells per μL, and then immediately loaded onto a 10× microfluidic chip (10× Genomics, v4) to generate single-cell mRNA libraries and then sequenced across six lanes on an Illumina X ten or NovaSeq 6000 system (Illumina, Inc., San Diego, California, US). Raw sequencing data were aligned to the GRH38 reference genome using the cellranger (10×Genomics, v4) count function. The count matrixes of gene expression from each sample were imported into the Seurat V4.1 (*49*). We selected high-quality cells for further analysis following three measurements: (1) cells had either over 2001 unique molecular identifiers (UMIs), fewer than 6,000 or more than 301 expressed genes or fewer than 20% UMIs derived from the mitochondrial genome; (2) genes expressed in more than ten cells in a sample; (3) cell doublets were removed using the DoubletFinder R package (V2.0.3) (*50*). The cell-by-gene expression matrixes of the remaining high-quality cells were integrated with the RunFastMNN function provided by the SeuratWrappers R package (V0.4.0) and then normalized to the total cellular UMI count. The union of the top 2,000 genes with the highest dispersion for each dataset was used to generate an integrated matrix. We then performed data normalization, dimension reduction, and cluster detection as previously reported (*23*). Briefly, the gene expression matrixes were scaled by regressing the total cellular UMI counts and percentage of mitochondrial genes. Principal component analysis (PCA) was conducted using highly variable genes (HVGs), and the top 30 significant principal components (PCs) were selected to perform uniform manifold approximation, projection (UMAP) dimension reduction, and visualization of gene expression. We annotated cell sub-clusters with similar gene expression patterns as the same cell type, and cell types in the resulting two-dimensional representation were annotated to known biological cell types using canonical marker genes.

### Detection of single-cell CNVs

We employed the inferCNV (*51*) R package (https://github.com/broadinstitute/inferCNV/wiki, v1.22.0) to distinguish malignant and non-malignant epithelia of GC, and the initial CNV signal of each region was estimated based on the expression level from the scRNA-seq results with default parameters.

### Pathway enrichment and cell type abundance deconvolution analysis

To illustrate the enriched signaling pathways of fibroblast and myeloid subtypes, we used the GSVA (v2.0.6) (*52*) package to assess pathway differences using the C2 curated gene set provided by the Molecular Signatures Database (MSigDB), which were calculated with a linear model offered by the limma package. We used the GSVA(*52*) package to calculate the abundance of malignant epithelium subclusters in *H. pylori* positive GC using the TCGA (*29*) and Asian Cancer Research Group (ACRG) cohorts (*28*). We used GSEA package to calculate the distribution of gene sets and cell-type-specific signatures (top 30 DEGs) in lists of genes ordered by population expression differences. Normalized enrichment score (NES).

### Malignant epithelium differentiation score

To assess the malignant epithelium differentiation heterogeneity at single cell level, we included a tumor differentiation-associated signature to evaluate the GC differentiation heterogeneity based on our previous study (*23*), including PHGR1, MUC13, MDK, KRT20, LGALS4, GPA33, CLDN3, CLDN4, and CDH17, and then we employed the “AddModuleScore” function of Seurat R package to calculate the differentiation degree of each tumor cell based on the expression level of differentiation-associated signature.

### TCGA data analysis

For the analysis of the correlation of tumor differentiation score and cell subtypes abundance with patients’ clinical outcome, we employed the GSVA to calculate the differentiation score and cell subtypes abundance in each TCGA stomach adenocarcinoma (STAD) sample based on the tumor differentiation signature and the top 30 DEGs of each cell subtype, and then grouped samples into high and low groups based on 67^th^ and 33^th^ percentile respectively (*53*). Kaplan-Meier survival curves were plotted using the R package “survminer” (v0.4.9).

### Immunotherapy response and prognosis analysis

GC immunotherapy bulk RNA-seq data along with curated clinical data from patients were obtained from a previous study (*37*). To evaluate the association between the signatures of immune and stromal subtypes identified in our study and immunotherapy response to GC patient survival, the GSVA (*52*) was used to calculate the combined expression value of the cell-type-specific signatures (top 30 DEGs). We classified the patients into high and low groups based on the 50^th^ and 50^th^ of cell subtypes abundance. Kaplan-Meier survival curves were plotted using the R package “survminer”. Subsequently, a Cox proportional hazards model was conducted that included age and tumor stage. The results of the Cox regression model among the cell subtypes were visualized using weighted Z-scores.

### Trajectory analysis

To explore the potential differentiation routines between the non-malignant epithelium of GC, we performed trajectory analysis using the monocle R package as previously reported (v3.0) (*54*). First, we constructed the monocle object using the “newCellDataSet” function, and the differentially expressed genes calculated via the “differentialGeneTest” function were selected for trajectory analysis. Then “DDRTree” function was used for dimensionality reduction and the “plot_cell_trajectory” function for visualization.

### Intercellular crosstalk

We used the Cellchat (*55*) package (v0.0.2) to infer the intercellular communications and significant ligand-receptor pair of the *H. pylori* infection associated GC TME, following a standard pipeline implemented in R (https://github.com/sqjin/CellChat). We first set the ligand-receptor interaction list in humans and projected the gene expression data onto the protein-protein interaction (PPI) network by identifying the overexpressed ligand-receptor interactions. To obtain biologically significant cell-cell communication, the probability values for each interaction were calculated by performing permutation tests. The inferred intercellular communication network of each ligand-receptor pair and each signaling pathway was summarized and visualized using circle plots and heatmaps.

### Multi-labelled immunofluorescence staining and multispectral imaging

A PANO 7-plex IHC kit (cat 10004100100; Panovue, Beijing, China) was used to perform multiplexed immunofluorescence staining. Slides were placed in a 65 °C oven overnight to for deparaffinization, and tissues were sequentially treated with xylene, ethanol, and distilled water. Slides were then microwaved (with antigen retrieval solution [citric acid solution, pH6.0/pH9.0]) and sequentially incubated with different primary antibodies and horseradish peroxidase-conjugated secondary antibodies (100 μL/tissue, table S7). This incubation was followed by tyramide signal amplification (TSA Fluorescence Kits, Panovue, Beijing, China). The slides were washed with 1× PBST after each incubation and microwaved after each round of TSA. After all the antigens were labelled, the nuclei were stained with 4’-6’-diamidino-2-phenylindole (DAPI, Sigma-Aldrich, MO, USA). Stained slides were scanned using a Mantra system (PerkinElmer, Waltham, Massachusetts, USA) to obtain multispectral images.

## Abbreviations used in this paper

GC: gastric cancer
*H. pylori*: *Helicobacter pylori*
HC: healthy control
non-HpGC: gastric cancer without *H. pylori* infection
HpGC: gastric cancer with current *H. pylori* infection
ex-HpGC: gastric cancer with previous *H. pylori* infection
CAF: cancer-associated fibroblasts
iCAF: inflammatory CAF
Vcaf: vascular-related CAF
mCAF: matrix CAF
EMT: epithelial-mesenchymal transition
Angio-TAM: angiogenic tumor-associated macrophages
TME: tumor microenvironment
ICIs: Immune checkpoint inhibitors
scRNA-seq: single-cell RNA sequencing
VEGFR: VEGF receptor
TIGIT: T cell immunoreceptor with Ig and ITIM domains
FACS: fluorescence-activated cell sorting
CNV: copy number variation
GO: Gene ontology
SPEM: spasmolytic polypeptide-expressing metaplasia
IM: intestinal metaplasia
TFs: transcription factors
TILs: Tumor infiltrating lymphocytes
NK: natural killer
ECM: extracellular matrix.

## Acknowledgments

We thank all patients enrolled in this study. All authors declare that they have no competing interests.

## Author contributions

CL, MZ, YW and SY were responsible for the study concept, design, interpretation and revising the manuscript. XZ, GZ, SS, YF and XC designed the experiments, collected the biopsies, analyzed the data, and wrote the manuscript. FL, JC, SR, and LP collected the biopsies. WS, HT and QG collected the biopsies, analyzed the data. HW and LZ evaluated gastric tumor pathology.

## Data and materials availability

The processed scRNA-seq data required to reproduce the analysis and figures have been deposited on Zenodo, which can be accessed with link https://zenodo.org/record/8082331. The TCGA GC bulk RNA-seq expression data was downloaded from the UCSC Xena website (https://xenabrowser.net/) and the GC immunotherapy expression matrix data can be accessed in European Nucleotide Archive with accession PRJEB25780. Other GC bulk RNA and microarray data used in this study are available in the NCBI database under accession code GEO accession numbers GSE62254, and GSE2669.

All data needed to evaluate the conclusions in the paper are present in the paper and/or the Supplementary Materials

## Reference

1. Y. Oya, Y. Hayakawa, K. Koike, Tumor microenvironment in gastric cancers. Cancer Sci 111, 2696–2707 (2020).

2. N. Uemura, S. Okamoto, S. Yamamoto, N. Matsumura, S. Yamaguchi, M. Yamakido, K. Taniyama, N. Sasaki, R. J. Schlemper, Helicobacter pylori infection and the development of gastric cancer. The New England journal of medicine 345, 784–789 (2001).

3. P. I. Hsu, K. H. Lai, P. N. Hsu, G. H. Lo, H. C. Yu, W. C. Chen, F. W. Tsay, H. C. Lin, H. H. Tseng, L. P. Ger, H. C. Chen, Helicobacter pylori infection and the risk of gastric malignancy. The American journal of gastroenterology 102, 725–730 (2007).

4. M. Plummer, S. Franceschi, J. Vignat, D. Forman, C. de Martel, Global burden of gastric cancer attributable to Helicobacter pylori. International journal of cancer 136, 487–490 (2015).

5. P. Oster, L. Vaillant, E. Riva, B. McMillan, C. Begka, C. Truntzer, C. Richard, M. M. Leblond, M. Messaoudene, E. Machremi, E. Limagne, F. Ghiringhelli, B. Routy, G. Verdeil, D. Velin, Helicobacter pylori infection has a detrimental impact on the efficacy of cancer immunotherapies. Gut 71, 457–466 (2022).

6. M. Hatakeyama, The role of Helicobacter pylori CagA in gastric carcinogenesis. Int J Hematol 84, 301–308 (2006).

7. H. W. Kwak, I. J. Choi, S. J. Cho, J. Y. Lee, C. G. Kim, M. C. Kook, K. W. Ryu, Y. W. Kim, Characteristics of gastric cancer according to Helicobacter pylori infection status. J Gastroenterol Hepatol 29, 1671–1677 (2014).

8. S. Y. Son, C. H. Lee, S. Y. Lee, Different Metabolites of the Gastric Mucosa between Patients with Current Helicobacter pylori Infection, Past Infection, and No Infection History. Biomedicines 10, (2022).

9. J. A. Thompson, New NCCN Guidelines: Recognition and Management of Immunotherapy-Related Toxicity. J Natl Compr Canc Netw 16, 594–596 (2018).

10. M. S. Carlino, J. Larkin, G. V. Long, Immune checkpoint inhibitors in melanoma. Lancet 398, 1002–1014 (2021).

11. S. Bagchi, R. Yuan, E. G. Engleman, Immune Checkpoint Inhibitors for the Treatment of Cancer: Clinical Impact and Mechanisms of Response and Resistance. Annual review of pathology 16, 223–249 (2021).

12. D. B. Doroshow, S. Bhalla, M. B. Beasley, L. M. Sholl, K. M. Kerr, S. Gnjatic, Wistuba, II, D. L. Rimm, M. S. Tsao, F. R. Hirsch, PD-L1 as a biomarker of response to immune-checkpoint inhibitors. Nature reviews. Clinical oncology 18, 345–362 (2021).

13. Y. Shi, H. Zheng, M. Wang, S. Ding, Influence of Helicobacter pylori infection on PD-1/PD-L1 blockade therapy needs more attention. Helicobacter 27, e12878 (2022).

14. Y. J. Bang, E. Y. Ruiz, E. Van Cutsem, K. W. Lee, L. Wyrwicz, M. Schenker, M. Alsina, M. H. Ryu, H. C. Chung, L. Evesque, S. E. Al-Batran, S. H. Park, M. Lichinitser, N. Boku, M. H. Moehler, J. Hong, H. Xiong, R. Hallwachs, I. Conti, J. Taieb, Phase III, randomised trial of avelumab versus physician’s choice of chemotherapy as third-line treatment of patients with advanced gastric or gastro-oesophageal junction cancer: primary analysis of JAVELIN Gastric 300. Annals of oncology : official journal of the European Society for Medical Oncology / ESMO 29, 2052–2060 (2018).

15. K. Shitara, M. Özgüroğ lu, Y. J. Bang, M. Di Bartolomeo, M. Mandalà, M. H. Ryu, L. Fornaro, T. Olesiński, C. Caglevic, H. C. Chung, K. Muro, E. Goekkurt, W. Mansoor, R. S. McDermott, E. Shacham-Shmueli, X. Chen, C. Mayo, S. P. Kang, A. Ohtsu, C. S. Fuchs, Pembrolizumab versus paclitaxel for previously treated, advanced gastric or gastro-oesophageal junction cancer (KEYNOTE-061): a randomised, open-label, controlled, phase 3 trial. *Lancet (London*, England) 392, 123–133 (2018).

16. K. Shitara, E. Van Cutsem, Y. J. Bang, C. Fuchs, L. Wyrwicz, K. W. Lee, I. Kudaba, M. Garrido, H. C. Chung, J. Lee, H. R. Castro, W. Mansoor, M. I. Braghiroli, N. Karaseva, C. Caglevic, L. Villanueva, E. Goekkurt, H. Satake, P. Enzinger, M. Alsina, A. Benson, J. Chao, A. H. Ko, Z. A. Wainberg, U. Kher, S. Shah, S. P. Kang, J. Tabernero, Efficacy and Safety of Pembrolizumab or Pembrolizumab Plus Chemotherapy vs Chemotherapy Alone for Patients With First-line, Advanced Gastric Cancer: The KEYNOTE-062 Phase 3 Randomized Clinical Trial. JAMA oncology 6, 1571–1580 (2020).

17. K. Shitara, J. A. Ajani, M. Moehler, M. Garrido, C. Gallardo, L. Shen, K. Yamaguchi, L. Wyrwicz, T. Skoczylas, A. C. Bragagnoli, T. Liu, M. Tehfe, E. Elimova, R. Bruges, T. Zander, S. de Azevedo, R. Kowalyszyn, R. Pazo-Cid, M. Schenker, J. M. Cleary, P. Yanez, K. Feeney, M. V. Karamouzis, V. Poulart, M. Lei, H. Xiao, K. Kondo, M. Li, Y. Y. Janjigian, Nivolumab plus chemotherapy or ipilimumab in gastro-oesophageal cancer. Nature 603, 942–948 (2022).

18. Z. Zhang, Z. Liu, Z. Chen, Comparison of Treatment Efficacy and Survival Outcomes Between Asian and Western Patients With Unresectable Gastric or Gastro-Esophageal Adenocarcinoma: A Systematic Review and Meta-Analysis. Front Oncol 12, 831207 (2022).

19. L. Peng, B. D. Qin, K. Xiao, S. Xu, J. S. Yang, Y. S. Zang, J. Stebbing, L. P. Xie, A meta-analysis comparing responses of Asian versus non-Asian cancer patients to PD-1 and PD-L1 inhibitor-based therapy. Oncoimmunology 9, 1781333 (2020).

20. S. H. Gohil, J. B. Iorgulescu, D. A. Braun, D. B. Keskin, K. J. Livak, Applying high-dimensional single-cell technologies to the analysis of cancer immunotherapy. Nature reviews. Clinical oncology 18, 244–256 (2021).

21. S. Z. Wu, A. Swarbrick, Single-cell advances in stromal-leukocyte interactions in cancer. Immunological reviews, (2021).

22. P. Zhang, M. Yang, Y. Zhang, S. Xiao, X. Lai, A. Tan, S. Du, S. Li, Dissecting the Single-Cell Transcriptome Network Underlying Gastric Premalignant Lesions and Early Gastric Cancer. Cell Rep 27, 1934–1947.e1935 (2019).

23. M. Zhang, S. Hu, M. Min, Y. Ni, Z. Lu, X. Sun, J. Wu, B. Liu, X. Ying, Y. Liu, Dissecting transcriptional heterogeneity in primary gastric adenocarcinoma by single cell RNA sequencing. Gut 70, 464–475 (2021).

24. R. Wang, M. Dang, K. Harada, G. Han, F. Wang, M. Pool Pizzi, M. Zhao, G. Tatlonghari, S. Zhang, D. Hao, Y. Lu, S. Zhao, B. D. Badgwell, M. Blum Murphy, N. Shanbhag, J. S. Estrella, S. Roy-Chowdhuri, A. A. F. Abdelhakeem, Y. Wang, G. Peng, S. Hanash, G. A. Calin, X. Song, Y. Chu, J. Zhang, M. Li, K. Chen, A. J. Lazar, A. Futreal, S. Song, J. A. Ajani, L. Wang, Single-cell dissection of intratumoral heterogeneity and lineage diversity in metastatic gastric adenocarcinoma. Nature medicine 27, 141–151 (2021).

25. K. Fu, B. Hui, Q. Wang, C. Lu, W. Shi, Z. Zhang, D. Rong, B. Zhang, Z. Tian, W. Tang, H. Cao, X. Wang, Z. Chen, Single-cell RNA sequencing of immune cells in gastric cancer patients. Aging 12, 2747–2763 (2020).

26. A. Sathe, S. M. Grimes, B. T. Lau, J. Chen, C. Suarez, R. J. Huang, G. Poultsides, H. P. Ji, Single-Cell Genomic Characterization Reveals the Cellular Reprogramming of the Gastric Tumor Microenvironment. Clinical cancer research : an official journal of the American Association for Cancer Research 26, 2640–2653 (2020).

27. R. Kim, M. An, H. Lee, A. Mehta, Y. J. Heo, K. M. Kim, S. Y. Lee, J. Moon, S. T. Kim, B. H. Min, T. J. Kim, S. Y. Rha, W. K. Kang, W. Y. Park, S. J. Klempner, J. Lee, Early Tumor-Immune Microenvironmental Remodeling and Response to First-Line Fluoropyrimidine and Platinum Chemotherapy in Advanced Gastric Cancer. Cancer discovery 12, 984–1001 (2022).

28. R. Cristescu, J. Lee, M. Nebozhyn, K. M. Kim, J. C. Ting, S. S. Wong, J. Liu, Y. G. Yue, J. Wang, K. Yu, X. S. Ye, I. G. Do, S. Liu, L. Gong, J. Fu, J. G. Jin, M. G. Choi, T. S. Sohn, J. H. Lee, J. M. Bae, S. T. Kim, S. H. Park, I. Sohn, S. H. Jung, P. Tan, R. Chen, J. Hardwick, W. K. Kang, M. Ayers, D. Hongyue, C. Reinhard, A. Loboda, S. Kim, A. Aggarwal, Molecular analysis of gastric cancer identifies subtypes associated with distinct clinical outcomes. Nature medicine 21, 449–456 (2015).

29. A. J. Bass, V. Thorsson, I. Shmulevich, S. M. Reynolds, M. Miller, B. Bernard, T. Hinoue, P. W. Laird, C. Curtis, H. Shen, D. J. Weisenberger, N. Schultz, R. Shen, N. Weinhold, D. P. Kelsen, R. Bowlby, A. Chu, K. Kasaian, A. J. Mungall, A. G. Robertson, P. Sipahimalani, A. Cherniack, G. Getz, Y. Liu, M. S. Noble, C. Pedamallu, C. Sougnez, A. Taylor-Weiner, R. Akbani, J.-S. Lee, W. Liu, G. B. Mills, D. Yang, W. Zhang, A. Pantazi, M. Parfenov, M. Gulley, M. B. Piazuelo, B. G. Schneider, J. Kim, A. Boussioutas, M. Sheth, J. A. Demchok, C. S. Rabkin, J. E. Willis, S. Ng, K. Garman, D. G. Beer, A. Pennathur, B. J. Raphael, H.-T. Wu, R. Odze, H. K. Kim, J. Bowen, K. M. Leraas, T. M. Lichtenberg, S. Weaver, M. McLellan, M. Wiznerowicz, R. Sakai, G. Getz, C. Sougnez, M. S. Lawrence, K. Cibulskis, L. Lichtenstein, S. Fisher, S. B. Gabriel, E. S. Lander, L. Ding, B. Niu, A. Ally, M. Balasundaram, I. Birol, R. Bowlby, D. Brooks, Y. S. N. Butterfield, R. Carlsen, A. Chu, J. Chu, E. Chuah, H.-J. E. Chun, A. Clarke, N. Dhalla, R. Guin, R. A. Holt, S. J. M. Jones, K. Kasaian, D. Lee, H. A. Li, E. Lim, Y. Ma, M. A. Marra, M. Mayo, R. A. Moore, A. J. Mungall, K. L. Mungall, K. Ming Nip, A. G. Robertson, J. E. Schein, P. Sipahimalani, A. Tam, N. Thiessen, R. Beroukhim, S. L. Carter, A. D. Cherniack, J. Cho, K. Cibulskis, D. DiCara, S. Frazer, S. Fisher, S. B. Gabriel, N. Gehlenborg, D. I. Heiman, J. Jung, J. Kim, E. S. Lander, M. S. Lawrence, L. Lichtenstein, P. Lin, M. Meyerson, A. I. Ojesina, C. Sekhar Pedamallu, G. Saksena, S. E. Schumacher, C. Sougnez, P. Stojanov, B. Tabak, A. Taylor-Weiner, D. Voet, M. Rosenberg, T. I. Zack, H. Zhang, L. Zou, A. Protopopov, N. Santoso, M. Parfenov, S. Lee, J. Zhang, H. S. Mahadeshwar, J. Tang, X. Ren, S. Seth, L. Yang, A. W. Xu, X. Song, A. Pantazi, R. Xi, C. A. Bristow, A. Hadjipanayis, J. Seidman, L. Chin, P. J. Park, R. Kucherlapati, R. Akbani, S. Ling, W. Liu, A. Rao, J. N. Weinstein, S.-B. Kim, J.-S. Lee, Y. Lu, G. Mills, P. W. Laird, T. Hinoue, D. J. Weisenberger, M. S. Bootwalla, P. H. Lai, H. Shen, T. Triche Jr, D. J. Van Den Berg, S. B. Baylin, J. G. Herman, G. Getz, L. Chin, Y. Liu, B. A. Murray, M. S. Noble, r. B. Arman Askoy, G. Ciriello, G. Dresdner, J. Gao, B. Gross, A. Jacobsen, W. Lee, R. Ramirez, C. Sander, N. Schultz, Y. Senbabaoglu, R. Sinha, S. Onur Sumer, Y. Sun, N. Weinhold, V. Thorsson, B. Bernard, L. Iype, R. W. Kramer, R. Kreisberg, M. Miller, S. M. Reynolds, H. Rovira, N. Tasman, I. Shmulevich, S. Ng, D. Haussler, J. M. Stuart, R. Akbani, S. Ling, W. Liu, A. Rao, J. N. Weinstein, R. G. W. Verhaak, G. B. Mills, M. D. M. Leiserson, B. J. Raphael, H.-T. Wu, B. S. Taylor, A. D. Black, J. Bowen, J. Ann Carney, J. M. Gastier-Foster, C. Helsel, K. M. Leraas, T. M. Lichtenberg, C. McAllister, N. C. Ramirez, T. R. Tabler, L. Wise, E. Zmuda, R. Penny, D. Crain, J. Gardner, K. Lau, E. Curely, D. Mallery, S. Morris, J. Paulauskis, T. Shelton, C. Shelton, M. Sherman, C. Benz, J.-H. Lee, K. Fedosenko, G. Manikhas, O. Potapova, O. Voronina, D. Belyaev, O. Dolzhansky, W. Kimryn Rathmell, J. Brzezinski, M. Ibbs, K. Korski, N. The Cancer Genome Atlas Research, I. Analysis Working Group: Dana-Farber Cancer, B. Institute for Systems, C. University of Southern, C. Memorial Sloan Kettering Cancer, B. C. C. Agency, E. The, L. B. I. Edythe, M. D. A. C. Center, S. Harvard Medical, C. University of North, U. Vanderbilt, C. Asan Medical, M. University of, I. National Cancer, U. Case Western Reserve, C. University of California at Santa, U. Duke, M. University of, P. University of, U. Brown, Brigham, H. Women’s, C. National Cancer, H. Nationwide Children’s, U. Washington, C. Greater Poland Cancer, K. U. Leuven, E. Genome Sequencing Center: The, L. B. I. Edythe, L. Washington University in St, B. C. C. A. Genome Characterization Centers, S. B. Harvard Medical, M. D. A. C. C. Women’s Hospital, C. University of Southern California Epigenome, U. The Sidney Kimmel Comprehensive Cancer Center at Johns Hopkins, E. Genome Data Analysis Centers: The, L. B. I. Edythe, C. Memorial Sloan-Kettering Cancer, S. C. University of California, F. University of California San, H. Biospecimen Core Resource: The Research Institute at Nationwide Children’s, C. International Genomics, A. Tissue Source Sites: Buck Institute for Research on, S. Chonnam National University Medical, D. City Clinical Oncology, Cureline, U. N. C. L. C. C. Center, Comprehensive molecular characterization of gastric adenocarcinoma. Nature 513, 202–209 (2014).

30. J. M. Chauvin, H. M. Zarour, TIGIT in cancer immunotherapy. J Immunother Cancer 8, (2020).

31. H. Harjunpaa, C. Guillerey, TIGIT as an emerging immune checkpoint. Clin Exp Immunol 200, 108–119 (2020).

32. Z. Ge, M. P. Peppelenbosch, D. Sprengers, J. Kwekkeboom, TIGIT, the Next Step Towards Successful Combination Immune Checkpoint Therapy in Cancer. Front Immunol 12, 699895 (2021).

33. C. Lin, H. He, H. Liu, R. Li, Y. Chen, Y. Qi, Q. Jiang, L. Chen, P. Zhang, H. Zhang, H. Li, W. Zhang, Y. Sun, J. Xu, Tumour-associated macrophages-derived CXCL8 determines immune evasion through autonomous PD-L1 expression in gastric cancer. Gut 68, 1764–1773 (2019).

34. X. Chen, E. Song, Turning foes to friends: targeting cancer-associated fibroblasts. Nat Rev Drug Discov 18, 99–115 (2019).

35. T. Nishina, Y. Deguchi, D. Ohshima, W. Takeda, M. Ohtsuka, S. Shichino, S. Ueha, S. Yamazaki, M. Kawauchi, E. Nakamura, C. Nishiyama, Y. Kojima, S. Adachi-Akahane, M. Hasegawa, M. Nakayama, M. Oshima, H. Yagita, K. Shibuya, T. Mikami, N. Inohara, K. Matsushima, N. Tada, H. Nakano, Interleukin-11-expressing fibroblasts have a unique gene signature correlated with poor prognosis of colorectal cancer. Nature communications 12, 2281 (2021).

36. M. Zhang, H. Yang, L. Wan, Z. Wang, H. Wang, C. Ge, Y. Liu, Y. Hao, D. Zhang, G. Shi, Y. Gong, Y. Ni, C. Wang, Y. Zhang, J. Xi, S. Wang, L. Shi, L. Zhang, W. Yue, X. Pei, B. Liu, X. Yan, Single-cell transcriptomic architecture and intercellular crosstalk of human intrahepatic cholangiocarcinoma. Journal of hepatology 73, 1118–1130 (2020).

37. S. T. Kim, R. Cristescu, A. J. Bass, K. M. Kim, J. I. Odegaard, K. Kim, X. Q. Liu, X. Sher, H. Jung, M. Lee, S. Lee, S. H. Park, J. O. Park, Y. S. Park, H. Y. Lim, H. Lee, M. Choi, A. Talasaz, P. S. Kang, J. Cheng, A. Loboda, J. Lee, W. K. Kang, Comprehensive molecular characterization of clinical responses to PD-1 inhibition in metastatic gastric cancer. Nature medicine 24, 1449–1458 (2018).

38. X. Hong, B. Cui, M. Wang, Z. Yang, L. Wang, Q. Xu, Systemic Immune-inflammation Index, Based on Platelet Counts and Neutrophil-Lymphocyte Ratio, Is Useful for Predicting Prognosis in Small Cell Lung Cancer. The Tohoku journal of experimental medicine 236, 297–304 (2015).

39. L. E. Wroblewski, R. M. Peek, Jr., K. T. Wilson, Helicobacter pylori and gastric cancer: factors that modulate disease risk. Clin Microbiol Rev 23, 713–739 (2010).

40. H. Chang, N. Kim, J. H. Park, R. H. Nam, Y. J. Choi, S. M. Park, Y. J. Choi, H. Yoon, C. M. Shin, D. H. Lee, Helicobacter pylori Might Induce TGF-β1-Mediated EMT by Means of cagE. Helicobacter 20, 438–448 (2015).

41. R. Silva, I. Gullo, F. Carneiro, The PD-1:PD-L1 immune inhibitory checkpoint in Helicobacter pylori infection and gastric cancer: a comprehensive review and future perspectives. Porto Biomed J 1, 4–11 (2016).

42. L. Holokai, J. Chakrabarti, T. Broda, J. Chang, J. A. Hawkins, N. Sundaram, L. E. Wroblewski, R. M. Peek, Jr., J. Wang, M. Helmrath, J. M. Wells, Y. Zavros, Increased Programmed Death-Ligand 1 is an Early Epithelial Cell Response to Helicobacter pylori Infection. PLoS Pathog 15, e1007468 (2019).

43. B. Shen, A. Qian, W. Lao, W. Li, X. Chen, B. Zhang, H. Wang, F. Yuan, Y. Sun, Relationship between Helicobacter pylori and expression of programmed death-1 and its ligand in gastric intraepithelial neoplasia and early-stage gastric cancer. Cancer Manag Res 11, 3909–3919 (2019).

44. D. K. Kim, J. Jeong, D. S. Lee, D. Y. Hyeon, G. W. Park, S. Jeon, K. B. Lee, J. Y. Jang, D. Hwang, H. M. Kim, K. Jung, PD-L1-directed PlGF/VEGF blockade synergizes with chemotherapy by targeting CD141(+) cancer-associated fibroblasts in pancreatic cancer. Nat Commun 13, 6292 (2022).

45. A. X. Zhu, A. R. Abbas, M. R. de Galarreta, Y. Guan, S. Lu, H. Koeppen, W. Zhang, C.-H. Hsu, A. R. He, B.-Y. Ryoo, T. Yau, A. O. Kaseb, A. M. Burgoyne, F. Dayyani, J. Spahn, W. Verret, R. S. Finn, H. C. Toh, A. Lujambio, Y. Wang, Molecular correlates of clinical response and resistance to atezolizumab in combination with bevacizumab in advanced hepatocellular carcinoma. Nature medicine, (2022).

46. D. Fukumura, J. Kloepper, Z. Amoozgar, D. G. Duda, R. K. Jain, Enhancing cancer immunotherapy using antiangiogenics: opportunities and challenges. Nature reviews. Clinical oncology 15, 325–340 (2018).

47. P. S. Hegde, J. J. Wallin, C. Mancao, Predictive markers of anti-VEGF and emerging role of angiogenesis inhibitors as immunotherapeutics. Seminars in cancer biology 52, 117–124 (2018).

48. S. Ragusa, B. Prat-Luri, A. González-Loyola, S. Nassiri, M. L. Squadrito, A. Guichard, S. Cavin, N. Gjorevski, D. Barras, G. Marra, M. P. Lutolf, J. Perentes, E. Corse, R. Bianchi, L. Wetterwald, J. Kim, G. Oliver, M. Delorenzi, M. De Palma, T. V. Petrova, Antiangiogenic immunotherapy suppresses desmoplastic and chemoresistant intestinal tumors in mice. The Journal of clinical investigation 130, 1199–1216 (2020).

49. T. Stuart, A. Butler, P. Hoffman, C. Hafemeister, E. Papalexi, W. M. Mauck, Y. Hao, M. Stoeckius, P. Smibert, R. Satija, Comprehensive Integration of Single-Cell Data. Cell 177, 1888–1902.e1821 (2019).

50. C. S. McGinnis, L. M. Murrow, Z. J. Gartner, DoubletFinder: Doublet Detection in Single-Cell RNA Sequencing Data Using Artificial Nearest Neighbors. Cell Systems 8, 329–337.e324 (2019).

51. I. Tirosh, A. S. Venteicher, C. Hebert, L. E. Escalante, A. P. Patel, K. Yizhak, J. M. Fisher, C. Rodman, C. Mount, M. G. Filbin, C. Neftel, N. Desai, J. Nyman, B. Izar, C. C. Luo, J. M. Francis, A. A. Patel, M. L. Onozato, N. Riggi, K. J. Livak, D. Gennert, R. Satija, B. V. Nahed, W. T. Curry, R. L. Martuza, R. Mylvaganam, A. J. Iafrate, M. P. Frosch, T. R. Golub, M. N. Rivera, G. Getz, O. Rozenblatt-Rosen, D. P. Cahill, M. Monje, B. E. Bernstein, D. N. Louis, A. Regev, M. L. Suvà, Single-cell RNA-seq supports a developmental hierarchy in human oligodendroglioma. Nature 539, 309–313 (2016).

52. S. Hänzelmann, R. Castelo, J. Guinney, GSVA: gene set variation analysis for microarray and RNA-Seq data. BMC bioinformatics 14, 7 (2013).

53. S. Cheng, Z. Li, R. Gao, B. Xing, Y. Gao, Y. Yang, S. Qin, L. Zhang, H. Ouyang, P. Du, L. Jiang, B. Zhang, Y. Yang, X. Wang, X. Ren, J.-X. Bei, X. Hu, Z. Bu, J. Ji, Z. Zhang, A pan-cancer single-cell transcriptional atlas of tumor infiltrating myeloid cells. Cell 184, 792–809.e723 (2021).

54. C. Trapnell, D. Cacchiarelli, J. Grimsby, P. Pokharel, S. Li, M. Morse, N. J. Lennon, K. J. Livak, T. S. Mikkelsen, J. L. Rinn, The dynamics and regulators of cell fate decisions are revealed by pseudotemporal ordering of single cells. Nature biotechnology 32, 381–386 (2014).

55. S. Jin, C. F. Guerrero-Juarez, L. Zhang, I. Chang, R. Ramos, C. H. Kuan, P. Myung, M. V. Plikus, Q. Nie, Inference and analysis of cell-cell communication using CellChat. Nature communications 12, 1088 (2021).

