## Supplemental Figures for "Single-cell dissection of prognostic architecture and immunotherapy response in *Helicobacter pylori* infection associated gastric cancer"

.

**Supplementary figures and legends**

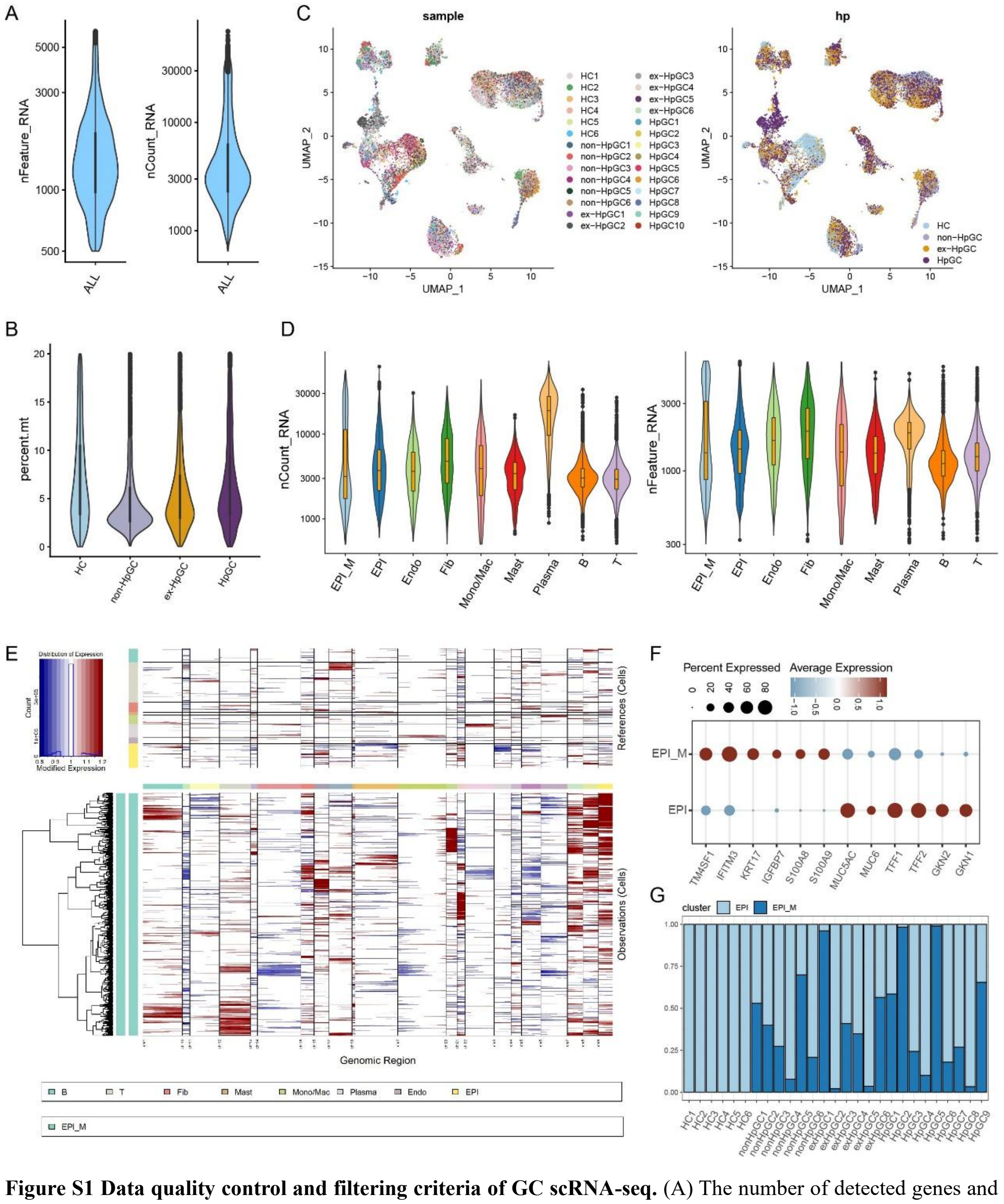


**Figure S1 Data quality control and filtering criteria of GC scRNA-seq.** (A) The number of detected genes and UMI counts in all the cells. (B) The percentage of mitochondria genome in gastric mucosal samples of HC, non-HpGC, ex-HpGC, and HpGC. (C) UMAP plot showing the sample (right) and pathological (left) distributions of 83,637 high-quality cells. (E) The copy number variation (CNV) signal across cell types, estimated using inferCNV. (F) The expression of DEGs in assumed non-malignant epithelium (EPI) and malignant epithelium (EPI_M). (G) Bar plot showing the percentage of EPI and EPI_M in each gastric mucosal sample of HC, non-HpGC, ex-HpGC, and HpGC.

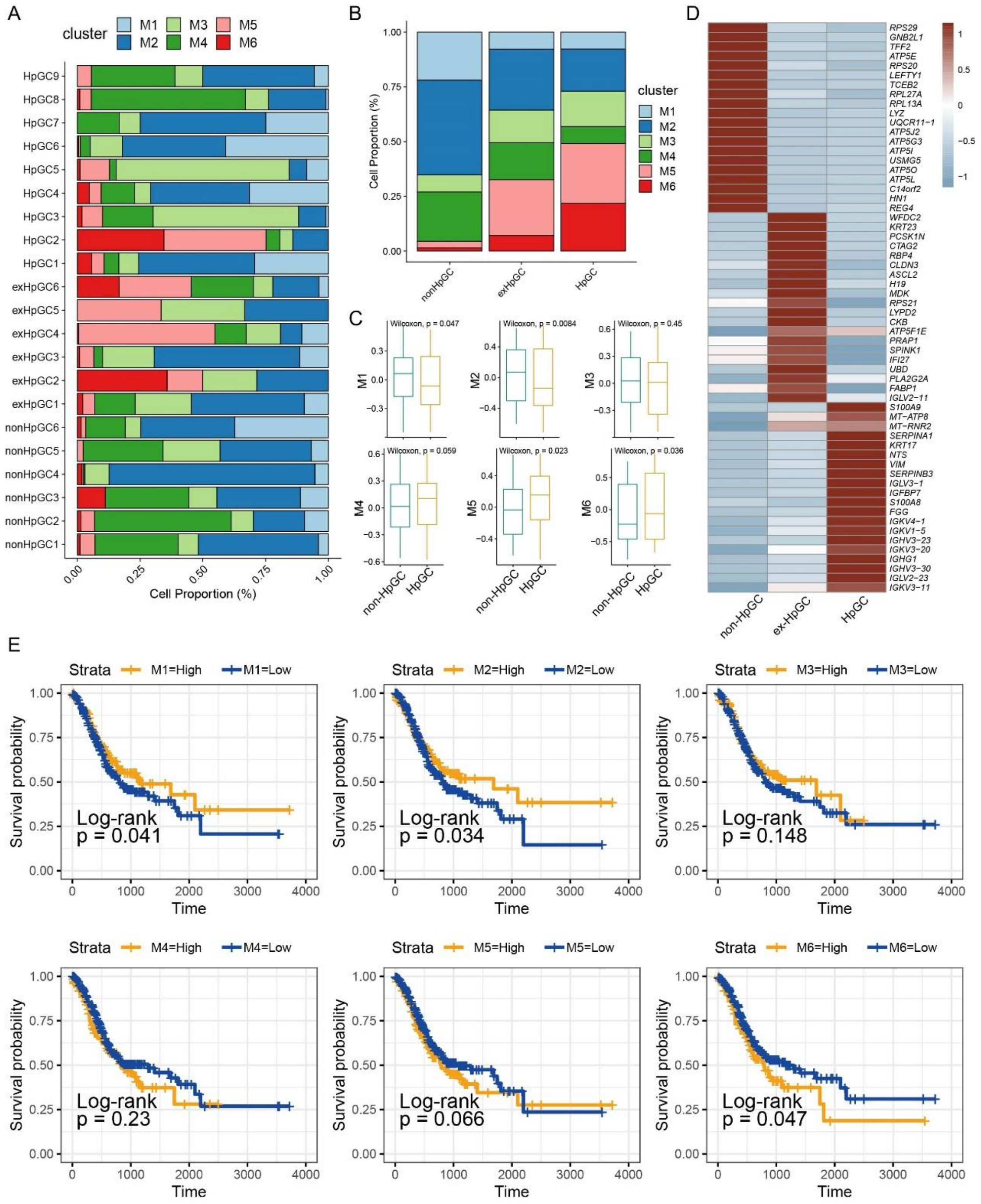


**Figure S2 Malignant epithelium characteristic in GC with different *H. pylori* infection status** (A-B) Proportion of six malignant epithelium subtypes in each GC sample (A) and non-HpGC, ex-HpGC, and HpGC(B). (C) The malignant epithelium subtypes relative abundance in *H. pylori* infection associated GC using TCGA STAD samples (estimated by GSVA). *P* values were assessed by Wilcoxon test. (D) Heatmap showing expression of top 20 DEGs in distinct malignant epithelium subtypes. (E) Kaplan-Meier survival analysis of TCGA STAD patients stratified by the relative abundance of six malignant epithelium subclusters, which was used to group samples into high and low groups based on 33$^{th}$ and 67$^{th}$ percentile. The P value of two-sided log-rank test is shown.

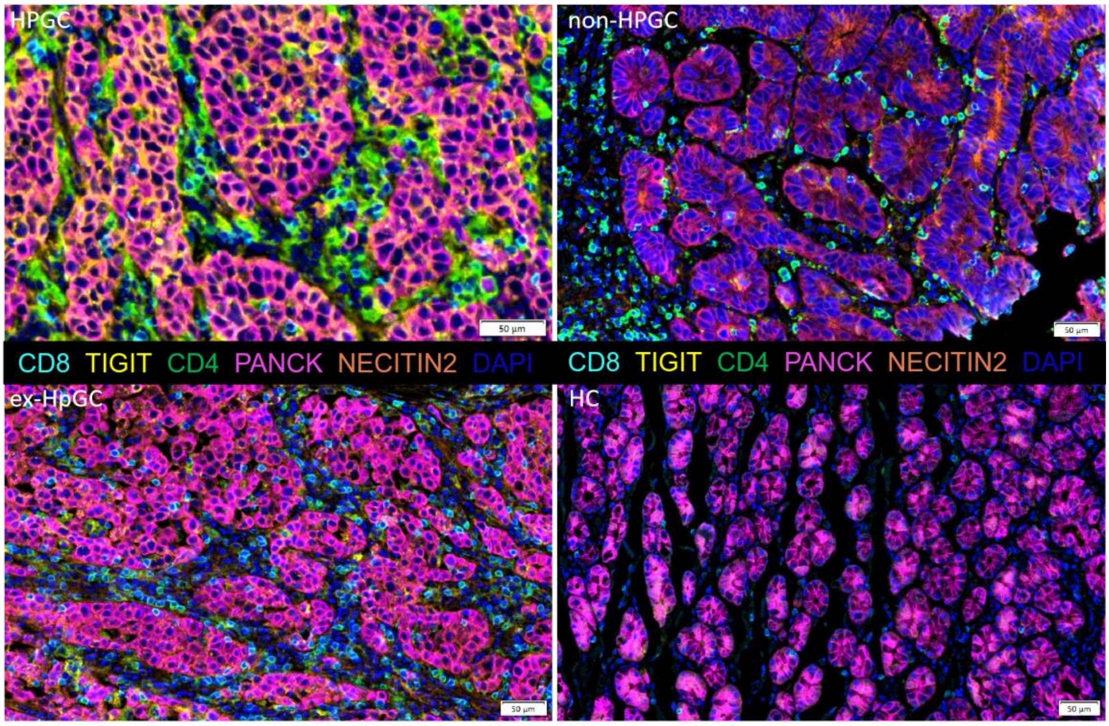


**Figure S3** Immunostaining of the ligand TIGIT and the receptor NECTIN 2 on suppressive T cells and on the malignant epithelium, respectively, in HpGC, ex-HpGC, non-HpGC and HC.

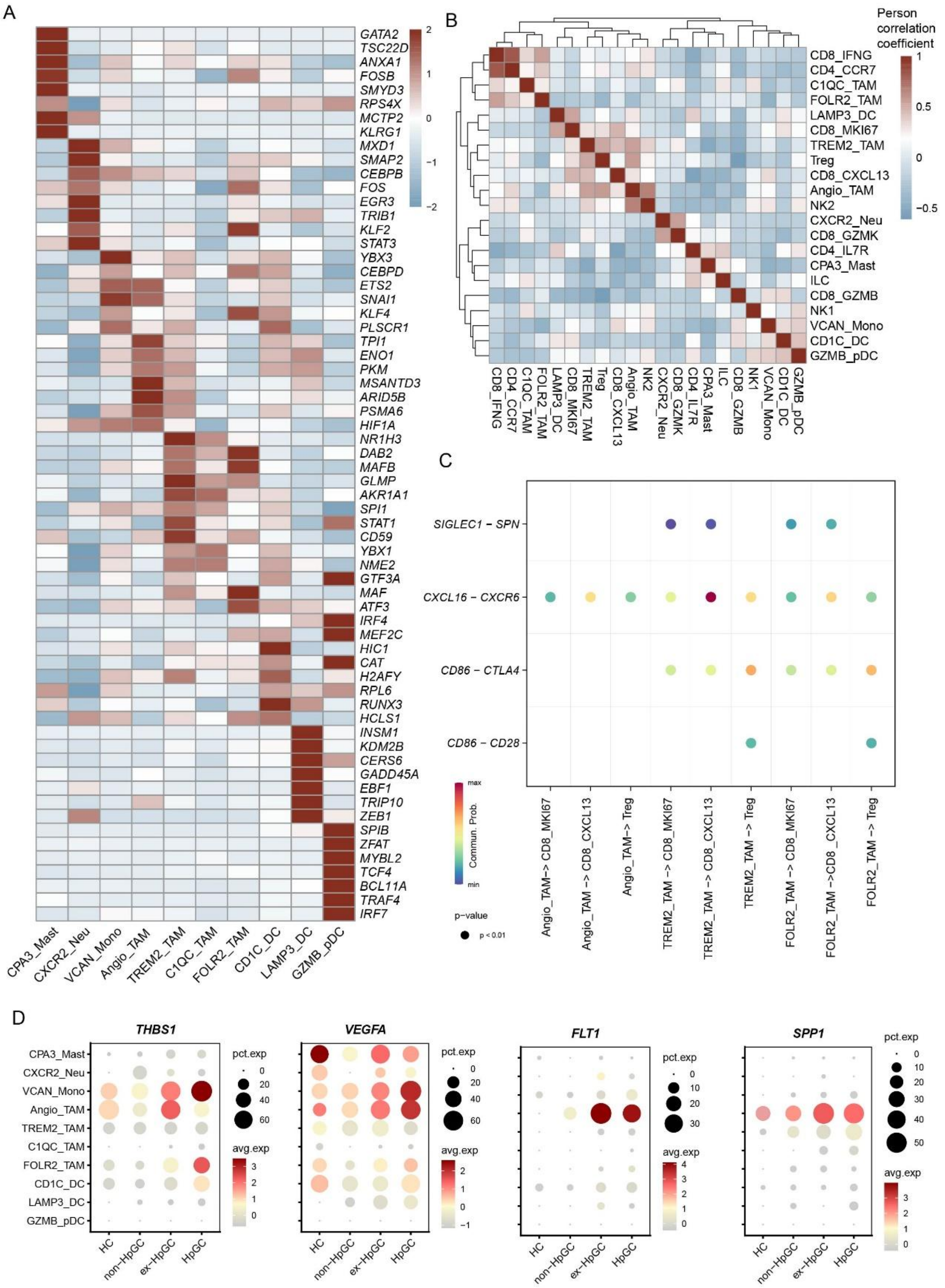


**Figure S4 Myeloid cell characteristic in GC with different *H. pylori* infection status** (A) Heatmap showing the top 7 TFs in different myeloid cell subtypes. (B) Heatmap showing the Pearson correlation coefficient of myeloid cell types and T/NK cell subtype abundance. (C) Bubble plot showing the expression of ligand-receptor gene pairs associated with immune signaling pathway networks: CD86, SN, and CXCL signaling pathways involved in the interactions between TAM and T cell subtypes. (D) Dotplot showing the expression of THBS1, VEGFA, FLT1

(VEGFR1), SPP1 in TAM derived from HC and GC with distinct *H. pylori* infection status.

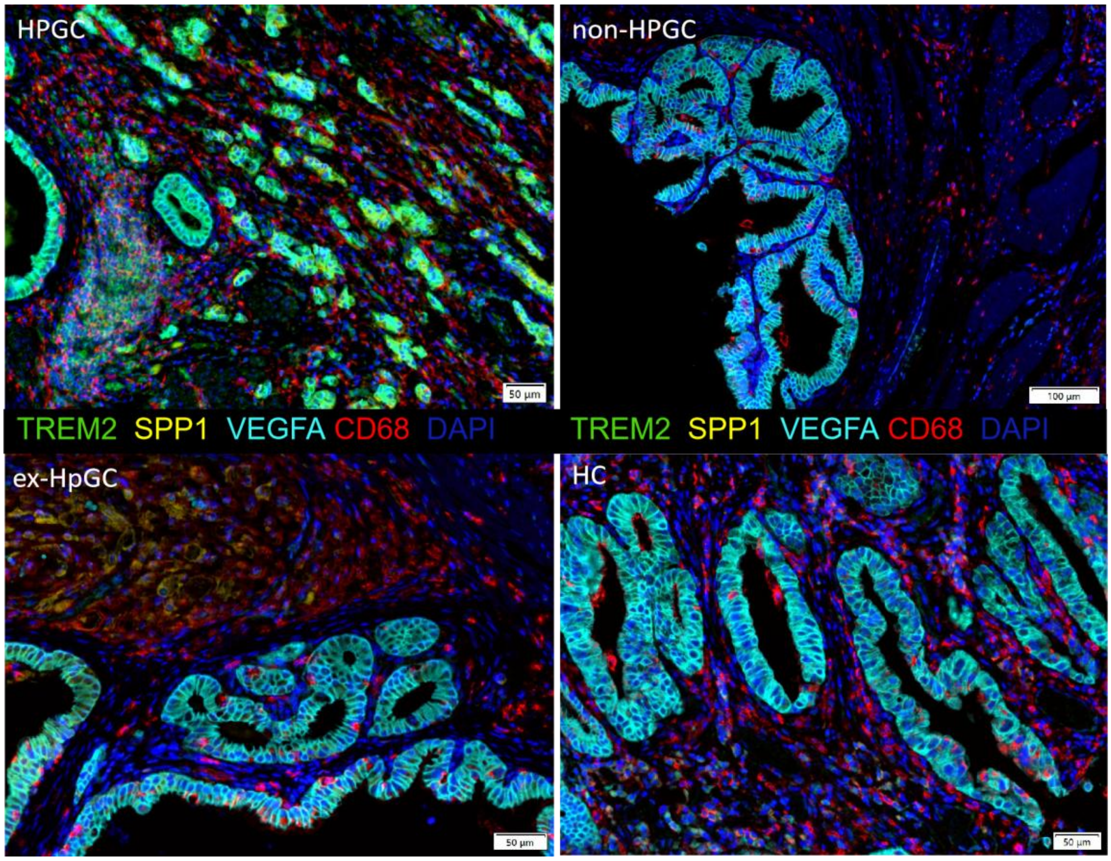


**Figure S5** Immunostaining showing the expression of Angio-TAM and TREM2+ TAM, respectively, in HpGC, ex-HpHC, non-HpGC and HC.

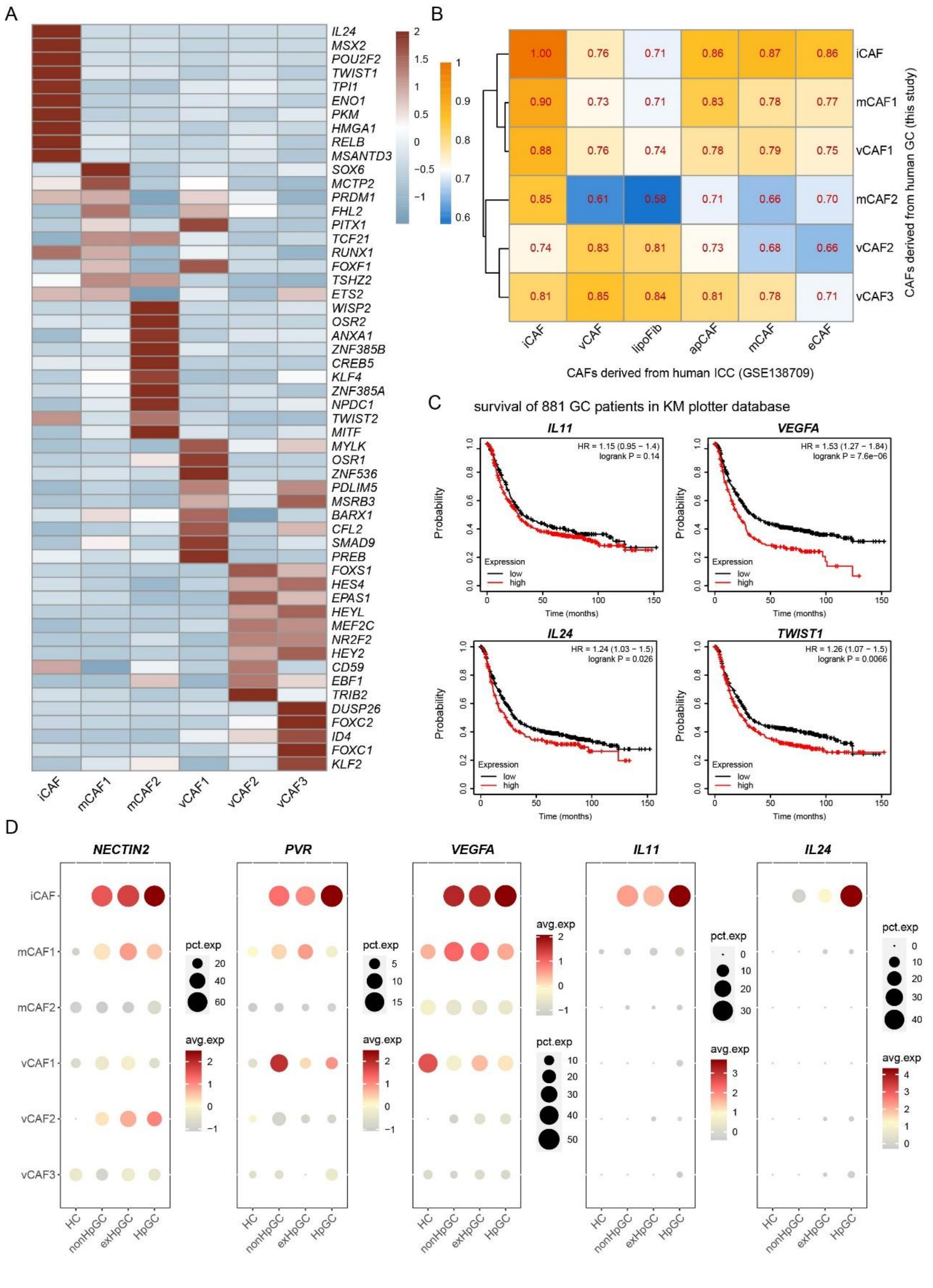


**Figure S6 CAF subtypes characteristic in GC with different *H. pylori* infection status** (A) Heatmap showing the top 5 TFs in different CAF subtypes. (B) Heatmap showing the Pearson correlation coefficient of CAF subtypes in human GC and ICC (GSE138709). (C) Kaplan-Meier survival analysis of 881 GC patients (km plotter database) stratified by high and low expression of iCAF associated genes IL11, IL23, VEGFA, TWIST1. The *P* value of two-sided log-rank test is shown. (D) Dotplot showing the expression of IL11, IL24, PVR, VEGFA, and NECTIN2 in

CAF subtypes derived from HC and GC with distinct *H. pylori* infection status.

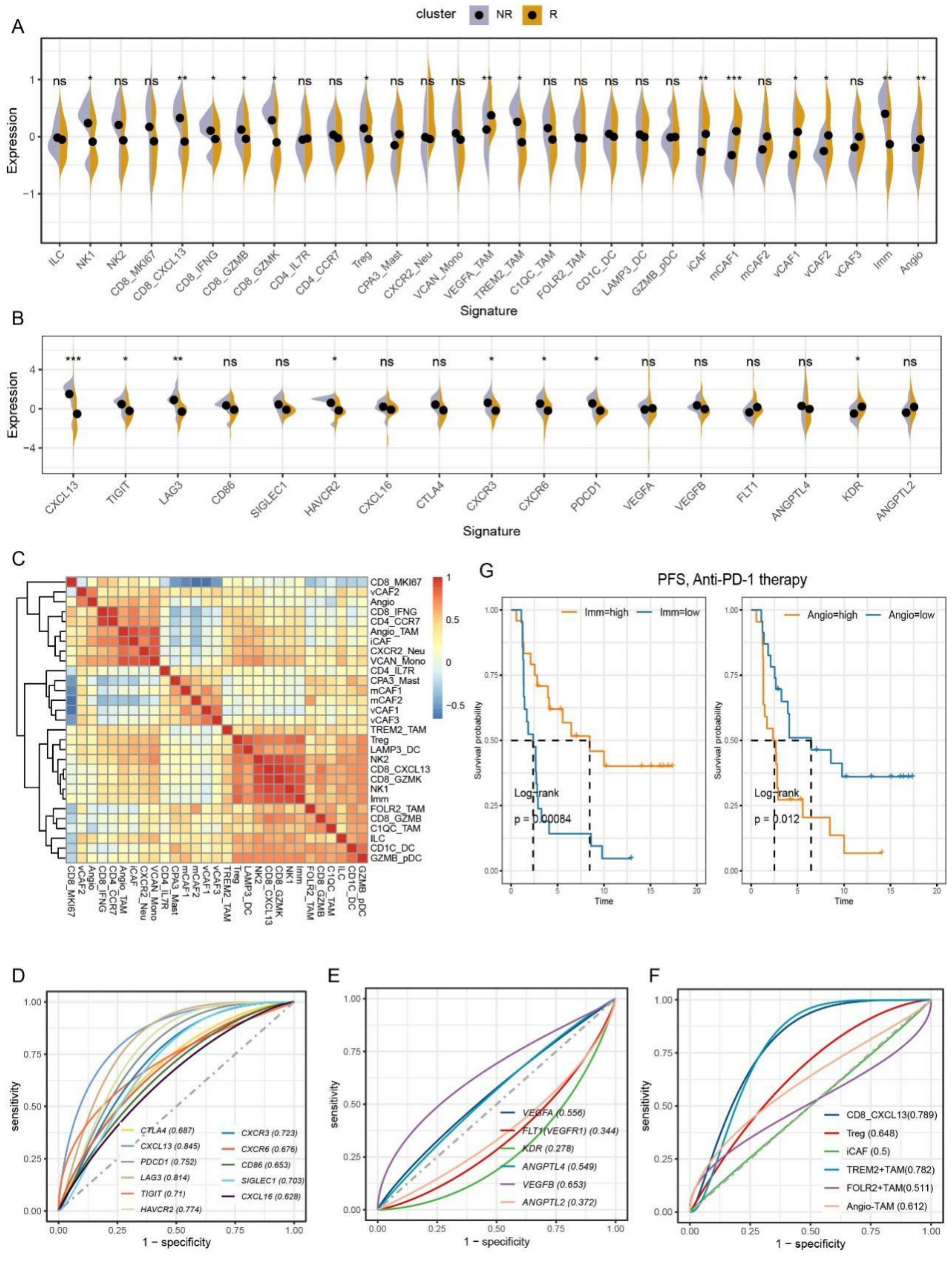


**Figure S7 GC TME characteristics related to GC immunotherapy response** (A) Relative abundances of cell types identified using scRNA-seq data predicts GC immunotherapy efficacy, R: responsive, NR: non-responsive. P values were assessed by Wilcoxon test. (B) The immune and angiogenesis signature identified using scRNA-seq data predicts GC immunotherapy efficacy, R: responsive, NR: non-responsive. P values were assessed by Wilcoxon test. (C) Heatmap showing the Pearson correlations between cell subtypes and immune and angiogenesis signature identified in the GC TME. (D–F) Evaluation of the sensitivity of GC to immunotherapy based on immune signature, angiogenesis signature and cell component. (G) Kaplan-Meier plot shows that the expression of Angio and Immune signature predicts anti-PD1 immunotherapy response (PFS) of GC.

**Supplementary tables legends:**

Supplementary Table S1: Top 30 DEGs of 9 main cell types.

Supplementary Table S2: Top 30 DEGs of 6 malignant epithelium subclusters.

Supplementary Table S3: Top 30 DEGs of 9 non-malignant epithelium subclusters

Supplementary Table S4: Top 30 DEGs of T/NK subclusters.

Supplementary Table S5: Top 30 DEGs of myeloid cells subclusters.

Supplementary Table S6: Top 30 DEGs of cancer associated fibroblasts subclusters.

Supplementary Table S7: Multilabel immunofluorescence staining antibody.
