## Supplemental Tables for "Single-cell dissection of prognostic architecture and immunotherapy response in *Helicobacter pylori* infection associated gastric cancer"

**Supplementary tables legends:**

Supplementary Table S1: Top 30 DEGs of 9 main cell types.

Supplementary Table S2: Top 30 DEGs of 6 malignant epithelium subclusters.

Supplementary Table S3: Top 30 DEGs of 9 non-malignant epithelium subclusters

Supplementary Table S4: Top 30 DEGs of T/NK subclusters.

Supplementary Table S5: Top 30 DEGs of myeloid cells subclusters.

Supplementary Table S6: Top 30 DEGs of cancer associated fibroblasts subclusters.

Supplementary Table S7: Multilabel immunofluorescence staining antibody.

Supplementary Table S1 DEGs of 9 main cell types

| | p_val | avg_log2FC | pct.1 | pct.2 | p_val_adj | cluster | gene |
|---|---|---|---|---|---|---|---|
| 1 | 0 | 4.054722338 | 0.427 | 0.016 | 0 | EPI_M | KRT17 |
| 2 | 0 | 3.413041725 | 0.333 | 0.082 | 0 | EPI_M | S100A9 |
| 3 | 0 | 2.999510382 | 0.245 | 0.032 | 0 | EPI_M | FABP1 |
| 4 | 0 | 2.893957158 | 0.246 | 0.035 | 0 | EPI_M | REG4 |
| 5 | 0 | 2.829650318 | 0.194 | 0.012 | 0 | EPI_M | S100A2 |
| 6 | 0 | 2.756827622 | 0.861 | 0.206 | 0 | EPI_M | KRT18 |
| 7 | 0 | 2.66663036 | 0.154 | 0.006 | 0 | EPI_M | NTS |
| 8 | 0 | 2.602133794 | 0.146 | 0.01 | 0 | EPI_M | KRT5 |
| 9 | 0 | 2.491240694 | 0.754 | 0.188 | 0 | EPI_M | KRT19 |
| 10 | 0 | 2.476826407 | 0.583 | 0.049 | 0 | EPI_M | CLDN4 |
| 11 | 0 | 2.448632278 | 0.339 | 0.062 | 0 | EPI_M | TFF3 |
| 12 | 0 | 2.307039015 | 0.892 | 0.237 | 0 | EPI_M | KRT8 |
| 13 | 0 | 2.271435561 | 0.504 | 0.137 | 0 | EPI_M | LGALS4 |
| 14 | 0 | 2.135031261 | 0.178 | 0.006 | 0 | EPI_M | SERPINB3 |
| 15 | 0 | 2.132341479 | 0.303 | 0.022 | 0 | EPI_M | PRAP1 |
| 16 | 0 | 2.115209813 | 0.147 | 0.005 | 0 | EPI_M | KRT6A |
| 17 | 0 | 2.070658467 | 0.131 | 0.007 | 0 | EPI_M | CALML3 |
| 18 | 0 | 2.062425708 | 0.577 | 0.081 | 0 | EPI_M | S100A14 |
| 19 | 0 | 2.041463822 | 0.637 | 0.102 | 0 | EPI_M | EPCAM |
| 20 | 0 | 2.03260156 | 0.541 | 0.087 | 0 | EPI_M | SFN |
| 21 | 0 | 1.99282924 | 0.121 | 0.003 | 0 | EPI_M | MIR205HG |
| 22 | 0 | 1.990148584 | 0.574 | 0.033 | 0 | EPI_M | CLDN7 |
| 23 | 0 | 1.965979268 | 0.393 | 0.017 | 0 | EPI_M | CLDN3 |
| 24 | 0 | 1.9493613 | 0.68 | 0.246 | 0 | EPI_M | SPINT2 |
| 25 | 0 | 1.874926443 | 0.53 | 0.176 | 0 | EPI_M | MDK |
| 26 | 1.8744E-281 | 2.065812171 | 0.28 | 0.066 | 6.1651E-277 | EPI_M | S100A8 |
| 27 | 3.3806E-274 | 2.679122632 | 0.115 | 0.008 | 1.1119E-269 | EPI_M | KRT14 |
| 28 | 1.08E-253 | 2.887899635 | 0.156 | 0.021 | 3.5522E-249 | EPI_M | SPINK4 |
| 29 | 7.7664E-194 | 1.882830101 | 0.123 | 0.017 | 2.5545E-189 | EPI_M | TCN1 |
| 30 | 1.0817E-156 | 1.869423317 | 0.19 | 0.053 | 3.5577E-152 | EPI_M | ANPEP |
| 31 | 0 | 7.253803327 | 0.673 | 0.098 | 0 | EPI | GKN1 |
| 32 | 0 | 6.72615374 | 0.875 | 0.155 | 0 | EPI | MUC5AC |
| 33 | 0 | 6.056063262 | 0.9 | 0.187 | 0 | EPI | TFF1 |
| 34 | 0 | 5.918745189 | 0.909 | 0.153 | 0 | EPI | TFF2 |
| 35 | 0 | 5.663176712 | 0.611 | 0.179 | 0 | EPI | PGC |
| 36 | 0 | 5.530326467 | 0.696 | 0.033 | 0 | EPI | GKN2 |
| 37 | 0 | 4.900684581 | 0.536 | 0.024 | 0 | EPI | MUCL3 |
| 38 | 0 | 4.493487012 | 0.816 | 0.07 | 0 | EPI | PSCA |
| 39 | 0 | 3.744852767 | 0.689 | 0.045 | 0 | EPI | CAPN8 |
| 40 | 0 | 3.632525586 | 0.834 | 0.202 | 0 | EPI | LYZ |
| 41 | 0 | 3.618300682 | 0.829 | 0.095 | 0 | EPI | SPINK1 |
| 42 | 0 | 3.558333688 | 0.907 | 0.105 | 0 | EPI | MUC1 |

| | | | | | | | |
|---|---|---|---|---|---|---|---|
| 43 | 0 | 3.52556685 | 0.878 | 0.095 | 0 | EPI | CA2 |
| 44 | 0 | 3.39578524 | 0.893 | 0.185 | 0 | EPI | AGR2 |
| 45 | 0 | 3.392050848 | 0.857 | 0.053 | 0 | EPI | CLDN18 |
| 46 | 0 | 3.284889532 | 0.921 | 0.292 | 0 | EPI | CYSTM1 |
| 47 | 0.00E+00 | 3.270290121 | 0.839 | 0.153 | 0.00E+00 | EPI | S100P |
| 48 | 0.00E+00 | 2.999656936 | 0.857 | 0.05 | 0.00E+00 | EPI | VSIG2 |
| 49 | 0.00E+00 | 2.865320575 | 0.79 | 0.047 | 0.00E+00 | EPI | CTSE |
| 50 | 0.00E+00 | 2.749997853 | 0.764 | 0.066 | 0.00E+00 | EPI | MLPH |
| 51 | 0 | 2.745397482 | 0.595 | 0.164 | 0 | EPI | PHGR1 |
| 52 | 0 | 2.737642916 | 0.846 | 0.136 | 0 | EPI | RNASE1 |
| 53 | 0 | 2.696211996 | 0.65 | 0.042 | 0 | EPI | SULT1C2 |
| 54 | 0 | 2.653207966 | 0.357 | 0.042 | 0 | EPI | MSMB |
| 55 | 0 | 2.585229362 | 0.598 | 0.123 | 0 | EPI | SYTL2 |
| 56 | 0.00E+00 | 2.572925665 | 0.547 | 0.039 | 0.00E+00 | EPI | RASEF |
| 57 | 0.00E+00 | 2.545856785 | 0.596 | 0.06 | 0.00E+00 | EPI | CYP3A5 |
| 58 | 2.34E-172 | 5.551961336 | 0.253 | 0.085 | 7.69E-168 | EPI | LIPF |
| 59 | 6.32E-151 | 4.147207672 | 0.233 | 0.081 | 2.08E-146 | EPI | MUC6 |
| 60 | 4.36E-20 | 4.557576723 | 0.128 | 0.078 | 1.43E-15 | EPI | PGA3 |
| 61 | 0.00E+00 | 4.688961822 | 0.867 | 0.017 | 0.00E+00 | Endo | PLVAP |
| 62 | 0.00E+00 | 3.866170172 | 0.884 | 0.039 | 0.00E+00 | Endo | RAMP2 |
| 63 | 0.00E+00 | 3.664445942 | 0.781 | 0.008 | 0.00E+00 | Endo | VWF |
| 64 | 0.00E+00 | 3.520215322 | 0.783 | 0.097 | 0.00E+00 | Endo | PECAM1 |
| 65 | 0.00E+00 | 3.432684838 | 0.974 | 0.153 | 0.00E+00 | Endo | IGFBP7 |
| 66 | 0.00E+00 | 3.411687919 | 0.834 | 0.109 | 0.00E+00 | Endo | HSPG2 |
| 67 | 0.00E+00 | 3.399554099 | 0.9 | 0.095 | 0.00E+00 | Endo | SPARCL1 |
| 68 | 0.00E+00 | 3.305634497 | 0.842 | 0.068 | 0.00E+00 | Endo | GNG11 |
| 69 | 0.00E+00 | 3.285060182 | 0.896 | 0.109 | 0.00E+00 | Endo | CRIP2 |
| 70 | 0.00E+00 | 3.247295427 | 0.634 | 0.151 | 0.00E+00 | Endo | CD320 |
| 71 | 0.00E+00 | 3.131868474 | 0.345 | 0.005 | 0.00E+00 | Endo | ACKR1 |
| 72 | 0.00E+00 | 3.049271534 | 0.688 | 0.056 | 0.00E+00 | Endo | COL4A1 |
| 73 | 0.00E+00 | 3.039092548 | 0.531 | 0.046 | 0.00E+00 | Endo | SERPINE1 |
| 74 | 0.00E+00 | 3.016249005 | 0.729 | 0.075 | 0.00E+00 | Endo | COL4A2 |
| 75 | 0.00E+00 | 3.014284677 | 0.594 | 0.005 | 0.00E+00 | Endo | CLDN5 |
| 76 | 0.00E+00 | 2.926019168 | 0.716 | 0.131 | 0.00E+00 | Endo | SLC9A3R2 |
| 77 | 0.00E+00 | 2.861359326 | 0.748 | 0.007 | 0.00E+00 | Endo | RAMP3 |
| 78 | 0.00E+00 | 2.861325013 | 0.782 | 0.065 | 0.00E+00 | Endo | ENG |
| 79 | 0.00E+00 | 2.812056402 | 0.753 | 0.026 | 0.00E+00 | Endo | EGFL7 |
| 80 | 0.00E+00 | 2.790500103 | 0.76 | 0.096 | 0.00E+00 | Endo | PRSS23 |
| 81 | 0.00E+00 | 2.781733328 | 0.755 | 0.055 | 0.00E+00 | Endo | HYAL2 |
| 82 | 0.00E+00 | 2.776178272 | 0.641 | 0.032 | 0.00E+00 | Endo | COL15A1 |
| 83 | 0.00E+00 | 2.747515682 | 0.849 | 0.103 | 0.00E+00 | Endo | A2M |
| 84 | 0.00E+00 | 2.631352416 | 0.851 | 0.114 | 0.00E+00 | Endo | SPARC |
| 85 | 0.00E+00 | 2.555698977 | 0.777 | 0.155 | 0.00E+00 | Endo | TM4SF1 |
| 86 | 0.00E+00 | 2.547594219 | 0.735 | 0.081 | 0.00E+00 | Endo | CAV1 |

| | | | | | | | |
|---|---|---|---|---|---|---|---|
| 87 | 0.00E+00 | 2.542027971 | 0.81 | 0.121 | 0.00E+00 | Endo | NPDC1 |
| 88 | 0.00E+00 | 2.531688339 | 0.973 | 0.369 | 0.00E+00 | Endo | IFITM3 |
| 89 | 0.00E+00 | 2.505860873 | 0.604 | 0.005 | 0.00E+00 | Endo | SOX18 |
| 90 | 0.00E+00 | 2.496679934 | 0.681 | 0.006 | 0.00E+00 | Endo | CLEC14A |
| 91 | 0.00E+00 | 5.259601243 | 0.555 | 0.104 | 0.00E+00 | Fib | CFD |
| 92 | 0.00E+00 | 5.122024731 | 0.863 | 0.036 | 0.00E+00 | Fib | DCN |
| 93 | 0.00E+00 | 4.903396286 | 0.508 | 0.034 | 0.00E+00 | Fib | CXCL14 |
| 94 | 0.00E+00 | 4.399875457 | 0.695 | 0.025 | 0.00E+00 | Fib | LUM |
| 95 | 0.00E+00 | 4.334972938 | 0.603 | 0.059 | 0.00E+00 | Fib | TAGLN |
| 96 | 0.00E+00 | 4.278053354 | 0.827 | 0.047 | 0.00E+00 | Fib | COL1A1 |
| 97 | 0.00E+00 | 4.223399416 | 0.828 | 0.029 | 0.00E+00 | Fib | COL3A1 |
| 98 | 0.00E+00 | 4.21026988 | 0.887 | 0.031 | 0.00E+00 | Fib | COL1A2 |
| 99 | 0.00E+00 | 4.18328619 | 0.637 | 0.085 | 0.00E+00 | Fib | MGP |
| 100 | 0.00E+00 | 4.143043101 | 0.473 | 0.019 | 0.00E+00 | Fib | APOD |
| 101 | 0 | 4.057834087 | 0.444 | 0.019 | 0 | Fib | POSTN |
| 102 | 0 | 3.941096578 | 0.304 | 0.025 | 0 | Fib | RGS5 |
| 103 | 0 | 3.927302404 | 0.746 | 0.012 | 0 | Fib | MFAP4 |
| 104 | 0 | 3.855965177 | 0.959 | 0.065 | 0 | Fib | CALD1 |
| 105 | 0 | 3.84441083 | 0.857 | 0.039 | 0 | Fib | RARRES2 |
| 106 | 0 | 3.812917298 | 0.695 | 0.029 | 0 | Fib | IGFBP5 |
| 107 | 0 | 3.789768794 | 0.568 | 0.057 | 0 | Fib | ACTA2 |
| 108 | 0 | 3.750973313 | 0.819 | 0.075 | 0 | Fib | MYL9 |
| 109 | 0 | 3.70682157 | 0.913 | 0.048 | 0 | Fib | COL6A2 |
| 110 | 0 | 3.683196773 | 0.624 | 0.021 | 0 | Fib | FBLN1 |
| 111 | 0 | 3.677016573 | 0.655 | 0.058 | 0 | Fib | CTGF |
| 112 | 0 | 3.632355462 | 0.857 | 0.029 | 0 | Fib | C1S |
| 113 | 0 | 3.5750177 | 0.584 | 0.038 | 0 | Fib | IGFBP6 |
| 114 | 0 | 3.511967663 | 0.876 | 0.056 | 0 | Fib | SERPING1 |
| 115 | 0 | 3.476485315 | 0.783 | 0.021 | 0 | Fib | SOD3 |
| 116 | 0 | 3.473831465 | 0.28 | 0.013 | 0 | Fib | SFRP2 |
| 117 | 0 | 3.416296971 | 0.941 | 0.127 | 0 | Fib | IGFBP7 |
| 118 | 0 | 3.376201742 | 0.698 | 0.03 | 0 | Fib | MMP2 |
| 119 | 0 | 3.337303032 | 0.807 | 0.068 | 0 | Fib | TPM2 |
| 120 | 0 | 3.322711177 | 0.735 | 0.038 | 0 | Fib | SERPINF1 |
| 121 | 0 | 4.547836218 | 0.495 | 0.056 | 0 | Mono/Mac | G0S2 |
| 122 | 0 | 4.344339009 | 0.964 | 0.092 | 0 | Mono/Mac | TYROBP |
| 123 | 0 | 4.178034395 | 0.491 | 0.025 | 0 | Mono/Mac | C1QA |
| 124 | 0 | 4.17306984 | 0.47 | 0.024 | 0 | Mono/Mac | C1QB |
| 125 | 0 | 4.157518963 | 0.472 | 0.052 | 0 | Mono/Mac | CXCL8 |
| 126 | 0 | 4.008809844 | 0.453 | 0.021 | 0 | Mono/Mac | C1QC |
| 127 | 0 | 3.768284307 | 0.844 | 0.032 | 0 | Mono/Mac | AIF1 |
| 128 | 0 | 3.712453028 | 0.877 | 0.068 | 0 | Mono/Mac | FCER1G |
| 129 | 0 | 3.659871653 | 0.992 | 0.918 | 0 | Mono/Mac | FTL |
| 130 | 0 | 3.561760873 | 0.386 | 0.021 | 0 | Mono/Mac | IL1B |

| | | | | | | | |
|---|---|---|---|---|---|---|---|
| 131 | 0 | 3.450910508 | 0.996 | 0.959 | 0 | Mono/Mac | FTH1 |
| 132 | 0 | 3.297717903 | 0.369 | 0.051 | 0 | Mono/Mac | APOC1 |
| 133 | 0 | 3.292526102 | 0.78 | 0.28 | 0 | Mono/Mac | SOD2 |
| 134 | 0 | 3.281307914 | 0.414 | 0.056 | 0 | Mono/Mac | CCL3 |
| 135 | 0 | 3.184517548 | 0.776 | 0.322 | 0 | Mono/Mac | CTSB |
| 136 | 0 | 3.127607516 | 0.496 | 0.069 | 0 | Mono/Mac | S100A8 |
| 137 | 0 | 2.93034274 | 0.753 | 0.386 | 0 | Mono/Mac | GPX1P1 |
| 138 | 0 | 2.922745195 | 0.741 | 0.246 | 0 | Mono/Mac | NAMPT |
| 139 | 0 | 2.807421661 | 0.833 | 0.418 | 0 | Mono/Mac | HLA-DRB1 |
| 140 | 0 | 2.801857909 | 0.309 | 0.04 | 0 | Mono/Mac | CCL3L1 |
| 141 | 0 | 2.748556222 | 0.977 | 0.738 | 0 | Mono/Mac | SAT1 |
| 142 | 0 | 2.738095452 | 0.765 | 0.062 | 0 | Mono/Mac | SPI1 |
| 143 | 0 | 2.731997337 | 0.852 | 0.343 | 0 | Mono/Mac | HLA-DRA |
| 144 | 0 | 2.708851011 | 0.627 | 0.115 | 0 | Mono/Mac | BCL2A1 |
| 145 | 0 | 2.693023243 | 0.581 | 0.011 | 0 | Mono/Mac | MS4A6A |
| 146 | 0.00E+00 | 2.690093564 | 0.785 | 0.386 | 0.00E+00 | Mono/Mac | HLA-DPA1 |
| 147 | 1.15E-290 | 2.778280491 | 0.156 | 0.01 | 3.77E-286 | Mono/Mac | SPP1 |
| 148 | 2.91E-267 | 3.210249262 | 0.367 | 0.08 | 9.58E-263 | Mono/Mac | APOE |
| 149 | 2.80E-204 | 2.72620728 | 0.716 | 0.377 | 9.21E-200 | Mono/Mac | TIMP1 |
| 150 | 4.84E-98 | 2.92353523 | 0.107 | 0.017 | 1.59E-93 | Mono/Mac | CXCL5 |
| 151 | 0 | 7.59572634 | 1 | 0.019 | 0 | Mast | TPSAB1 |
| 152 | 0 | 6.382788133 | 0.771 | 0.007 | 0 | Mast | TPSB2 |
| 153 | 0 | 5.973748945 | 0.953 | 0.007 | 0 | Mast | CPA3 |
| 154 | 0 | 5.222454927 | 0.689 | 0.003 | 0 | Mast | TPSD1 |
| 155 | 0 | 4.206454752 | 0.893 | 0.013 | 0 | Mast | GATA2 |
| 156 | 0 | 4.035471579 | 0.842 | 0.013 | 0 | Mast | KIT |
| 157 | 0 | 3.491360869 | 0.873 | 0.049 | 0 | Mast | LTC4S |
| 158 | 0 | 3.290300119 | 0.86 | 0.005 | 0 | Mast | HPGDS |
| 159 | 0 | 3.193257706 | 0.851 | 0.042 | 0 | Mast | VWA5A |
| 160 | 0 | 2.874894577 | 0.687 | 0.001 | 0 | Mast | MS4A2 |
| 161 | 0 | 2.860959021 | 0.884 | 0.153 | 0 | Mast | CLU |
| 162 | 0 | 2.860240823 | 0.76 | 0.005 | 0 | Mast | SLC18A2 |
| 163 | 0 | 2.822214216 | 0.729 | 0.007 | 0 | Mast | HDC |
| 164 | 0 | 2.652889342 | 0.522 | 0.062 | 0 | Mast | CSF1 |
| 165 | 0 | 2.538732757 | 0.927 | 0.239 | 0 | Mast | ALOX5AP |
| 166 | 0 | 2.518784599 | 0.571 | 0.004 | 0 | Mast | RHEX |
| 167 | 0 | 2.50881192 | 0.716 | 0.008 | 0 | Mast | IL1RL1 |
| 168 | 0 | 2.39788218 | 0.56 | 0.016 | 0 | Mast | SLC24A3 |
| 169 | 0 | 2.35741531 | 0.707 | 0.07 | 0 | Mast | ALOX5 |
| 170 | 4.4316E-264 | 2.324922241 | 0.862 | 0.263 | 1.4576E-259 | Mast | CD9 |
| 171 | 1.9946E-235 | 2.156266767 | 0.876 | 0.303 | 6.5603E-231 | Mast | CAPG |
| 172 | 2.7219E-224 | 2.380694401 | 0.891 | 0.329 | 8.9524E-220 | Mast | GLUL |
| 173 | 3.2354E-219 | 2.342072947 | 1 | 0.65 | 1.0642E-214 | Mast | VIM |
| 174 | 1.1729E-184 | 2.514230371 | 0.724 | 0.234 | 3.8577E-180 | Mast | SOCS1 |

| | | | | | | | |
|---|---|---|---|---|---|---|---|
| 175 | 1.0708E-180 | 2.217248343 | 0.902 | 0.398 | 3.5221E-176 | Mast | ANXA1 |
| 176 | 8.7068E-180 | 2.373028048 | 0.831 | 0.353 | 2.8637E-175 | Mast | NFKBIZ |
| 177 | 4.2719E-165 | 2.920594081 | 0.693 | 0.22 | 1.4051E-160 | Mast | AREG |
| 178 | 4.6319E-162 | 2.514675601 | 0.858 | 0.426 | 1.5235E-157 | Mast | LMNA |
| 179 | 1.015E-116 | 2.169238097 | 0.804 | 0.43 | 3.3383E-112 | Mast | TNFAIP3 |
| 180 | 1.81867E-87 | 2.171201366 | 0.713 | 0.362 | 5.98177E-83 | Mast | TUBA1A |
| 181 | 0 | 6.765640317 | 0.872 | 0.181 | 0 | Plasma | JCHAIN |
| 182 | 0 | 6.306404222 | 0.555 | 0.096 | 0 | Plasma | IGLV6-57 |
| 183 | 0 | 6.221141815 | 0.238 | 0.033 | 0 | Plasma | IGKV1-39 |
| 184 | 0 | 6.166206289 | 0.784 | 0.104 | 0 | Plasma | IGHA1 |
| 185 | 0 | 5.808893678 | 0.647 | 0.108 | 0 | Plasma | IGKC |
| 186 | 0 | 5.761954903 | 0.29 | 0.05 | 0 | Plasma | IGKV1-12 |
| 187 | 0 | 5.749367666 | 0.788 | 0.09 | 0 | Plasma | IGLL5 |
| 188 | 0 | 5.49677199 | 0.243 | 0.03 | 0 | Plasma | IGKV1-9 |
| 189 | 9.648E-249 | 6.183568841 | 0.301 | 0.068 | 3.1733E-244 | Plasma | IGHV4-34 |
| 190 | 5.6051E-233 | 5.807679713 | 0.189 | 0.029 | 1.8436E-228 | Plasma | IGKV1D-39 |
| 191 | 5.67E-219 | 6.48531046 | 0.491 | 0.172 | 1.86E-214 | Plasma | IGKV1-5 |
| 192 | 3.60E-216 | 5.629917656 | 0.162 | 0.022 | 1.18E-211 | Plasma | IGKV1-27 |
| 193 | 2.68E-190 | 6.277270488 | 0.358 | 0.105 | 8.81E-186 | Plasma | IGKV3-11 |
| 194 | 5.76E-185 | 6.064430403 | 0.218 | 0.047 | 1.90E-180 | Plasma | IGHV1-69D |
| 195 | 6.16E-184 | 5.501041015 | 0.251 | 0.061 | 2.03E-179 | Plasma | IGHV3-30 |
| 196 | 2.17E-173 | 5.535749234 | 0.484 | 0.174 | 7.15E-169 | Plasma | IGKV4-1 |
| 197 | 2.06E-168 | 5.706270513 | 0.517 | 0.197 | 6.78E-164 | Plasma | IGKV3-20 |
| 198 | 6.12E-166 | 5.599775575 | 0.203 | 0.045 | 2.01E-161 | Plasma | IGHV3-7 |
| 199 | 6.29E-164 | 5.543893441 | 0.182 | 0.037 | 2.07E-159 | Plasma | IGHV3-48 |
| 200 | 1.79E-163 | 5.61081759 | 0.151 | 0.026 | 5.89E-159 | Plasma | IGKV2D-28 |
| 201 | 2.7828E-151 | 5.470551056 | 0.287 | 0.084 | 9.1528E-147 | Plasma | IGKV3-15 |
| 202 | 4.7982E-131 | 5.684557706 | 0.241 | 0.07 | 1.5782E-126 | Plasma | IGLV2-11 |
| 203 | 1.2405E-115 | 6.368835916 | 0.301 | 0.103 | 4.0802E-111 | Plasma | IGLV1-40 |
| 204 | 3.864E-107 | 5.999689152 | 0.384 | 0.156 | 1.2709E-102 | Plasma | IGLV2-14 |
| 205 | 4.7519E-102 | 5.906791755 | 0.218 | 0.069 | 1.56296E-97 | Plasma | IGLV1-47 |
| 206 | 1.4669E-101 | 5.475757025 | 0.156 | 0.04 | 4.82491E-97 | Plasma | IGLV3-10 |
| 207 | 2.65088E-77 | 5.66802791 | 0.162 | 0.051 | 8.719E-73 | Plasma | IGLV7-46 |
| 208 | 3.30102E-72 | 5.727515216 | 0.203 | 0.074 | 1.08574E-67 | Plasma | IGLV1-51 |
| 209 | 2.269E-37 | 5.816434064 | 0.103 | 0.039 | 7.46296E-33 | Plasma | IGHV2-5 |
| 210 | 4.46914E-05 | 6.615027491 | 0.112 | 0.094 | 1 | Plasma | IGJ |
| 211 | 0 | 3.557263712 | 0.895 | 0.011 | 0 | B | MS4A1 |
| 212 | 0 | 3.07981699 | 0.992 | 0.308 | 0 | B | HLA-DRA |
| 213 | 0 | 3.065014073 | 0.81 | 0.121 | 0 | B | CD83 |
| 214 | 0 | 3.0017772 | 0.932 | 0.123 | 0 | B | CD79A |
| 215 | 0 | 2.873596996 | 0.723 | 0.016 | 0 | B | VPREB3 |
| 216 | 0 | 2.794145244 | 0.999 | 0.718 | 0 | B | CD74 |
| 217 | 0 | 2.47223076 | 0.944 | 0.345 | 0 | B | CD37 |
| 218 | 0 | 2.468477932 | 0.904 | 0.208 | 0 | B | HLA-DQB1 |

| | | | | | | | |
|---|---|---|---|---|---|---|---|
| 219 | 0 | 2.365823874 | 0.899 | 0.172 | 0 | B | HLA-DQA1 |
| 220 | 0 | 2.356116207 | 0.665 | 0.024 | 0 | B | BANK1 |
| 221 | 0 | 2.202495054 | 0.956 | 0.301 | 0 | B | HLA-DPB1 |
| 222 | 0 | 2.173817414 | 0.54 | 0.076 | 0 | B | CD79B |
| 223 | 0 | 2.163566252 | 0.965 | 0.351 | 0 | B | HLA-DPA1 |
| 224 | 0 | 2.137560557 | 0.972 | 0.386 | 0 | B | HLA-DRB1 |
| 225 | 0 | 2.075998211 | 0.308 | 0.002 | 0 | B | TCL1A |
| 226 | 0 | 2.028077416 | 0.66 | 0.133 | 0 | B | MEF2C |
| 227 | 0 | 1.992703977 | 0.607 | 0.069 | 0 | B | IRF8 |
| 228 | 0 | 1.859152019 | 0.905 | 0.357 | 0 | B | CXCR4 |
| 229 | 0 | 1.784929604 | 0.524 | 0.039 | 0 | B | CD19 |
| 230 | 0 | 1.717802635 | 0.742 | 0.173 | 0 | B | LTB |
| 231 | 0 | 1.715112414 | 0.389 | 0.016 | 0 | B | LINC00926 |
| 232 | 0 | 1.712392697 | 0.448 | 0.091 | 0 | B | LY9 |
| 233 | 0 | 1.709452685 | 0.732 | 0.226 | 0 | B | HLA-DMA |
| 234 | 0 | 1.697332736 | 0.571 | 0.185 | 0 | B | SMIM14 |
| 235 | 0 | 1.66144718 | 0.914 | 0.378 | 0 | B | LAPTM5 |
| 236 | 0 | 1.657851588 | 0.412 | 0.027 | 0 | B | TNFRSF13C |
| 237 | 0 | 1.637698369 | 0.469 | 0.025 | 0 | B | FCRLA |
| 238 | 0 | 1.602090752 | 0.895 | 0.326 | 0 | B | CD52 |
| 239 | 0 | 1.58862688 | 0.508 | 0.067 | 0 | B | HLA-DMB |
| 240 | 0 | 1.581181582 | 0.585 | 0.186 | 0 | B | HLA-DRB5 |
| 241 | 0 | 4.230426471 | 0.64 | 0.072 | 0 | T | CCL5 |
| 242 | 0 | 3.535315164 | 0.782 | 0.044 | 0 | T | CD7 |
| 243 | 0 | 3.505606155 | 0.499 | 0.111 | 0 | T | CCL4 |
| 244 | 0 | 3.467505972 | 0.888 | 0.031 | 0 | T | CD3E |
| 245 | 0 | 3.440005543 | 0.5 | 0.025 | 0 | T | GZMA |
| 246 | 0 | 3.378254803 | 0.606 | 0.05 | 0 | T | IL7R |
| 247 | 0 | 3.376013791 | 0.197 | 0.031 | 0 | T | GNLY |
| 248 | 0 | 3.374470948 | 0.805 | 0.016 | 0 | T | CD3D |
| 249 | 0 | 3.318268364 | 0.522 | 0.045 | 0 | T | NKG7 |
| 250 | 0 | 2.997979212 | 0.916 | 0.277 | 0 | T | IL32 |
| 251 | 0 | 2.912521846 | 0.748 | 0.021 | 0 | T | CD2 |
| 252 | 0 | 2.838831376 | 0.295 | 0.081 | 0 | T | CCL4L2 |
| 253 | 0 | 2.820022029 | 0.392 | 0.023 | 0 | T | GZMB |
| 254 | 0 | 2.661377176 | 0.434 | 0.016 | 0 | T | KLRB1 |
| 255 | 0 | 2.554579372 | 0.574 | 0.041 | 0 | T | CST7 |
| 256 | 0 | 2.271205239 | 0.855 | 0.221 | 0 | T | PTPRC |
| 257 | 0 | 2.249765312 | 0.726 | 0.134 | 0 | T | HCST |
| 258 | 0 | 2.232746803 | 0.385 | 0.011 | 0 | T | CD8A |
| 259 | 0 | 2.220892417 | 0.654 | 0.217 | 0 | T | DUSP2 |
| 260 | 0 | 2.218173149 | 0.597 | 0.015 | 0 | T | CD3G |
| 261 | 0 | 2.188651098 | 0.224 | 0.007 | 0 | T | GZMK |
| 262 | 0 | 2.158233434 | 0.68 | 0.12 | 0 | T | FYN |

| | | | | | | | |
|---|---|---|---|---|---|---|---|
| 263 | 0 | 2.134606266 | 0.398 | 0.013 | 0 | T | PRF1 |
| 264 | 0 | 2.119064192 | 0.735 | 0.173 | 0 | T | EVL |
| 265 | 0 | 2.085778058 | 0.715 | 0.208 | 0 | T | LEPROTL1 |
| 266 | 0 | 2.078349615 | 0.633 | 0.039 | 0 | T | LCK |
| 267 | 0 | 2.049736337 | 0.689 | 0.168 | 0 | T | ETS1 |
| 268 | 0 | 2.035081687 | 0.296 | 0.012 | 0 | T | GZMH |
| 269 | 0 | 2.001877469 | 0.672 | 0.471 | 0 | T | DNAJB1 |
| 270 | 0 | 1.994019097 | 0.729 | 0.467 | 0 | T | SARAF |

Supplementary Table S2 DEGs of 6 malignant epithelium subclusters

| | p_val | avg_log2FC | pct.1 | pct.2 | p_val_adj | cluster | gene |
|---|---|---|---|---|---|---|---|
| 1 | 6.96E-251 | 0.769026606 | 0.434 | 0.069 | 2.05E-246 | M1 | ASCL2 |
| 2 | 3.43E-249 | 1.232585432 | 0.77 | 0.286 | 1.01E-244 | M1 | CLDN3 |
| 3 | 3.54E-235 | 0.686406905 | 0.642 | 0.18 | 1.04E-230 | M1 | PRAP1 |
| 4 | 2.26E-232 | 0.670613821 | 0.359 | 0.046 | 6.66E-228 | M1 | DPEP1 |
| 5 | 1.17E-228 | 0.964736562 | 0.913 | 0.424 | 3.44E-224 | M1 | GPX2 |
| 6 | 1.31E-221 | 1.461702181 | 0.838 | 0.386 | 3.86E-217 | M1 | MDK |
| 7 | 2.22E-213 | 0.826910663 | 0.656 | 0.212 | 6.55E-209 | M1 | CES2 |
| 8 | 4.13E-203 | 1.548636946 | 0.984 | 0.845 | 1.22E-198 | M1 | RPL7 |
| 9 | 1.50E-197 | 0.939325645 | 0.913 | 0.463 | 4.42E-193 | M1 | TSPAN8 |
| 10 | 2.02E-197 | 0.72431177 | 0.59 | 0.19 | 5.96E-193 | M1 | GGH |
| 11 | 3.93E-196 | 0.899125075 | 0.66 | 0.227 | 1.16E-191 | M1 | TMEM176A |
| 12 | 2.10E-165 | 0.672716497 | 0.651 | 0.238 | 6.19E-161 | M1 | TMEM176B |
| 13 | 1.42E-155 | 1.444460783 | 0.902 | 0.501 | 4.20E-151 | M1 | RPS20 |
| 14 | 5.56E-155 | 0.814201166 | 0.903 | 0.491 | 1.64E-150 | M1 | CLDN4 |
| 15 | 1.45E-153 | 1.100381684 | 0.488 | 0.156 | 4.26E-149 | M1 | WFDC2 |
| 16 | 2.58E-151 | 1.298933098 | 0.687 | 0.297 | 7.62E-147 | M1 | LYZ |
| 17 | 2.72E-147 | 0.923150361 | 0.731 | 0.33 | 8.01E-143 | M1 | SPINK1 |
| 18 | 2.06E-141 | 2.000490944 | 0.425 | 0.138 | 6.07E-137 | M1 | OLFM4 |
| 19 | 1.15E-138 | 1.212814328 | 0.397 | 0.115 | 3.40E-134 | M1 | DMBT1 |
| 20 | 6.59E-125 | 0.757335193 | 0.298 | 0.072 | 1.94E-120 | M1 | RBP4 |
| 21 | 9.24E-115 | 0.738948073 | 0.387 | 0.122 | 2.72E-110 | M1 | PCK1 |
| 22 | 2.60E-109 | 0.71074746 | 0.906 | 0.588 | 7.67E-105 | M1 | IFI27 |
| 23 | 1.78E-108 | 0.744010559 | 0.351 | 0.111 | 5.24E-104 | M1 | PSMA2-1 |
| 24 | 7.47E-107 | 0.685471769 | 0.34 | 0.105 | 2.20E-102 | M1 | ATP5C1 |
| 25 | 3.12E-106 | 0.785150536 | 0.952 | 0.647 | 9.21E-102 | M1 | RPL27A |
| 26 | 3.27E-106 | 0.675422502 | 0.224 | 0.043 | 9.64E-102 | M1 | KRT23 |
| 27 | 3.27E-103 | 0.63053219 | 0.342 | 0.105 | 9.64E-99 | M1 | ATP5F1 |
| 28 | 7.85E-100 | 0.703362996 | 0.349 | 0.114 | 2.31E-95 | M1 | ATP5B |
| 29 | 2.30E-99 | 0.674398984 | 0.342 | 0.113 | 6.77E-95 | M1 | C14orf166 |
| 30 | 1.03E-97 | 0.936016032 | 0.376 | 0.142 | 3.04E-93 | M1 | ATP5G2 |
| 31 | 8.38E-92 | 1.210580823 | 0.957 | 0.721 | 2.47E-87 | M1 | LGALS3 |
| 32 | 9.56E-92 | 0.904457523 | 0.366 | 0.134 | 2.82E-87 | M1 | ATP5O |
| 33 | 1.11E-87 | 0.963665711 | 0.379 | 0.155 | 3.28E-83 | M1 | ATP5L |

| | | | | | | | |
|---|---|---|---|---|---|---|---|
| 34 | 5.24E-87 | 1.299720329 | 0.444 | 0.187 | 1.54E-82 | M1 | REG1A |
| 35 | 1.63E-85 | 0.740654917 | 0.35 | 0.127 | 4.80E-81 | M1 | ATP5G1 |
| 36 | 2.74E-85 | 1.281190956 | 0.387 | 0.171 | 8.08E-81 | M1 | GNB2L1 |
| 37 | 3.24E-84 | 0.976957298 | 0.518 | 0.243 | 9.55E-80 | M1 | PIGR |
| 38 | 1.08E-83 | 0.937880934 | 0.371 | 0.142 | 3.20E-79 | M1 | ATP5G3 |
| 39 | 4.53E-81 | 0.864703138 | 0.306 | 0.097 | 1.34E-76 | M1 | PGC |
| 40 | 4.20E-80 | 0.788152034 | 0.366 | 0.148 | 1.24E-75 | M1 | SHFM1 |
| 41 | 2.92E-78 | 0.689534014 | 0.931 | 0.659 | 8.60E-74 | M1 | RPS29 |
| 42 | 1.95E-77 | 0.932861115 | 0.704 | 0.377 | 5.76E-73 | M1 | LCN2 |
| 43 | 1.82E-75 | 0.665263077 | 0.372 | 0.152 | 5.37E-71 | M1 | C14orf2 |
| 44 | 8.77E-73 | 0.943817598 | 0.303 | 0.107 | 2.59E-68 | M1 | UBD |
| 45 | 4.31E-72 | 0.732238568 | 0.371 | 0.156 | 1.27E-67 | M1 | USMG5 |
| 46 | 1.27E-65 | 0.630404319 | 0.376 | 0.157 | 3.75E-61 | M1 | ATP5J2 |
| 47 | 1.75E-63 | 1.199664954 | 0.121 | 0.02 | 5.16E-59 | M1 | REG3A |
| 48 | 2.74E-57 | 0.739596253 | 0.388 | 0.178 | 8.09E-53 | M1 | ATP5E |
| 49 | 4.77E-57 | 0.712521891 | 0.958 | 0.79 | 1.41E-52 | M1 | RPL13A |
| 50 | 3.89E-17 | 0.651642263 | 0.331 | 0.209 | 1.15E-12 | M1 | PLA2G2A |
| 51 | 0 | 3.513970594 | 0.728 | 0.146 | 0 | M2 | TFF2 |
| 52 | 0 | 2.839064322 | 0.748 | 0.191 | 0 | M2 | REG4 |
| 53 | 0 | 2.699879105 | 0.557 | 0.083 | 0 | M2 | LEFTY1 |
| 54 | 0 | 2.535232578 | 0.81 | 0.127 | 0 | M2 | CLDN18 |
| 55 | 0 | 2.295917875 | 0.991 | 0.661 | 0 | M2 | KRT19 |
| 56 | 0 | 2.1763819 | 0.575 | 0.063 | 0 | M2 | CXCL17 |
| 57 | 0 | 2.071506481 | 0.858 | 0.11 | 0 | M2 | CTSE |
| 58 | 0 | 2.041916245 | 0.687 | 0.121 | 0 | M2 | ARL14 |
| 59 | 0 | 1.99138795 | 0.667 | 0.115 | 0 | M2 | PSCA |
| 60 | 0 | 1.948844912 | 0.974 | 0.523 | 0 | M2 | FXYD3 |
| 61 | 0 | 1.657296184 | 0.76 | 0.172 | 0 | M2 | ANXA10 |
| 62 | 0 | 1.571153987 | 0.893 | 0.255 | 0 | M2 | TSPAN1 |
| 63 | 0 | 1.5709594 | 0.907 | 0.276 | 0 | M2 | TMC5 |
| 64 | 0 | 1.479212404 | 0.687 | 0.119 | 0 | M2 | AKR1B10 |
| 65 | 0 | 1.473860928 | 0.85 | 0.211 | 0 | M2 | PIGR |
| 66 | 0 | 1.386773816 | 0.346 | 0.019 | 0 | M2 | DPCR1 |
| 67 | 0 | 1.284528836 | 0.6 | 0.064 | 0 | M2 | MIA |
| 68 | 0 | 1.227959585 | 0.791 | 0.159 | 0 | M2 | LAMB3 |
| 69 | 0 | 1.201847531 | 0.51 | 0.068 | 0 | M2 | B3GNT7 |
| 70 | 0 | 1.186793729 | 0.759 | 0.176 | 0 | M2 | MUC13 |
| 71 | 8.97E-308 | 1.384087896 | 0.813 | 0.231 | 2.64E-303 | M2 | PLAC8 |
| 72 | 2.06E-303 | 1.465056845 | 0.824 | 0.222 | 6.08E-299 | M2 | PDZK1IP1 |
| 73 | 3.58E-293 | 1.734816845 | 0.977 | 0.427 | 1.06E-288 | M2 | LGALS4 |
| 74 | 4.26E-284 | 1.920741642 | 0.868 | 0.298 | 1.26E-279 | M2 | LYZ |
| 75 | 1.93E-281 | 2.562952862 | 0.816 | 0.272 | 5.68E-277 | M2 | TFF3 |
| 76 | 4.36E-276 | 1.318748429 | 0.693 | 0.176 | 1.29E-271 | M2 | SRD5A3 |
| 77 | 3.87E-256 | 1.504143645 | 0.837 | 0.298 | 1.14E-251 | M2 | AGR3 |

| | | | | | | | |
|---|---|---|---|---|---|---|---|
| 78 | 4.84E-254 | 1.268363697 | 0.9 | 0.343 | 1.43E-249 | M2 | C12orf75 |
| 79 | 1.43E-250 | 1.647842532 | 0.961 | 0.49 | 4.21E-246 | M2 | TSPAN8 |
| 80 | 1.39E-249 | 1.362349542 | 0.914 | 0.39 | 4.09E-245 | M2 | NQO1 |
| 81 | 1.57E-243 | 2.091263579 | 0.909 | 0.369 | 4.63E-239 | M2 | LCN2 |
| 82 | 1.48E-237 | 3.022852519 | 0.868 | 0.456 | 4.35E-233 | M2 | TFF1 |
| 83 | 7.67E-218 | 1.167039198 | 0.878 | 0.351 | 2.26E-213 | M2 | TACSTD2 |
| 84 | 1.47E-208 | 1.496404338 | 0.822 | 0.277 | 4.34E-204 | M2 | C15orf48 |
| 85 | 1.54E-190 | 1.424288256 | 0.759 | 0.269 | 4.55E-186 | M2 | MT1E |
| 86 | 1.93E-184 | 1.662481889 | 0.967 | 0.825 | 5.70E-180 | M2 | AGR2 |
| 87 | 5.99E-178 | 1.22435433 | 0.463 | 0.114 | 1.77E-173 | M2 | SELK |
| 88 | 1.21E-168 | 1.35731472 | 0.476 | 0.126 | 3.56E-164 | M2 | HN1 |
| 89 | 1.16E-165 | 1.197561039 | 0.713 | 0.264 | 3.41E-161 | M2 | CA2 |
| 90 | 1.35E-146 | 1.604930361 | 0.725 | 0.301 | 3.98E-142 | M2 | MT1X |
| 91 | 2.63E-145 | 1.281835498 | 0.664 | 0.244 | 7.75E-141 | M2 | MT1G |
| 92 | 3.14E-133 | 1.293422927 | 0.811 | 0.347 | 9.26E-129 | M2 | SPINK1 |
| 93 | 4.73E-131 | 1.305881599 | 0.685 | 0.276 | 1.40E-126 | M2 | GDF15 |
| 94 | 6.72E-126 | 1.29454084 | 0.277 | 0.05 | 1.98E-121 | M2 | IL8 |
| 95 | 2.18E-103 | 1.156792057 | 0.825 | 0.452 | 6.44E-99 | M2 | CCT2 |
| 96 | 4.21E-100 | 1.549436859 | 0.375 | 0.111 | 1.24E-95 | M2 | CCL20 |
| 97 | 6.78E-94 | 1.48405792 | 0.182 | 0.027 | 2.00E-89 | M2 | SAA1 |
| 98 | 6.23E-59 | 1.253523883 | 0.441 | 0.208 | 1.84E-54 | M2 | CEACAM6 |
| 99 | 8.00E-28 | 1.211857165 | 0.253 | 0.122 | 2.36E-23 | M2 | PGC |
| 100 | 1.98E-14 | 1.388838828 | 0.105 | 0.045 | 5.85E-10 | M2 | LYPD2 |
| 101 | 9.65E-207 | 1.799653198 | 0.705 | 0.353 | 2.85E-202 | M3 | PHGR1 |
| 102 | 1.74E-125 | 1.755826249 | 0.167 | 0.016 | 5.15E-121 | M3 | REG1B |
| 103 | 4.11E-82 | 1.366381975 | 0.407 | 0.19 | 1.21E-77 | M3 | REG1A |
| 104 | 5.93E-77 | 1.028156086 | 0.846 | 0.818 | 1.75E-72 | M3 | RPL13A |
| 105 | 5.83E-76 | 1.017602249 | 0.813 | 0.685 | 1.72E-71 | M3 | RPS29 |
| 106 | 8.92E-62 | 1.836188983 | 0.333 | 0.168 | 2.63E-57 | M3 | SPINK4 |
| 107 | 4.81E-45 | 0.87116966 | 0.753 | 0.696 | 1.42E-40 | M3 | RPL27A |
| 108 | 5.59E-43 | 0.570114194 | 0.65 | 0.459 | 1.65E-38 | M3 | LGALS4 |
| 109 | 1.60E-36 | 0.687582201 | 0.627 | 0.493 | 4.71E-32 | M3 | GPX2 |
| 110 | 3.86E-36 | 0.405559993 | 0.996 | 0.997 | 1.14E-31 | M3 | S100A6 |
| 111 | 2.17E-34 | 0.465502174 | 0.532 | 0.376 | 6.40E-30 | M3 | SPINK1 |
| 112 | 2.07E-27 | 0.660437716 | 0.121 | 0.048 | 6.12E-23 | M3 | SNORD3A |
| 113 | 5.62E-27 | 0.625331083 | 0.641 | 0.566 | 1.66E-22 | M3 | RPS20 |
| 114 | 8.07E-26 | 1.071986683 | 0.563 | 0.545 | 2.38E-21 | M3 | HIST1H4C |
| 115 | 2.97E-24 | 1.224017658 | 0.318 | 0.238 | 8.75E-20 | M3 | CENPW |
| 116 | 4.80E-24 | 0.978101181 | 0.296 | 0.195 | 1.42E-19 | M3 | TCEB2 |
| 117 | 2.86E-23 | 0.837131011 | 0.301 | 0.198 | 8.43E-19 | M3 | ATP5E |
| 118 | 1.69E-22 | 0.952123853 | 0.254 | 0.165 | 4.98E-18 | M3 | ATP5D |
| 119 | 1.57E-21 | 1.044955653 | 0.19 | 0.115 | 4.62E-17 | M3 | SNORA76 |
| 120 | 2.50E-20 | 1.040420091 | 0.273 | 0.189 | 7.37E-16 | M3 | ATP5I |
| 121 | 8.02E-19 | 0.816804709 | 0.567 | 0.566 | 2.37E-14 | M3 | TMEM141 |

| | | | | | | | |
|---|---|---|---|---|---|---|---|
| 122 | 1.36E-15 | 1.234405159 | 0.356 | 0.286 | 4.02E-11 | M3 | MT1G |
| 123 | 2.75E-14 | 0.716542665 | 0.361 | 0.311 | 8.12E-10 | M3 | CDC42EP5 |
| 124 | 1.44E-13 | 0.559669047 | 0.432 | 0.371 | 4.25E-09 | M3 | CLDN3 |
| 125 | 1.51E-12 | 0.701385842 | 0.281 | 0.213 | 4.45E-08 | M3 | FABP1 |
| 126 | 3.25E-11 | 0.58932616 | 0.26 | 0.192 | 9.59E-07 | M3 | UQCR11-1 |
| 127 | 3.26E-10 | 0.847041344 | 0.304 | 0.266 | 9.62E-06 | M3 | PRAP1 |
| 128 | 3.72E-10 | 0.661714351 | 0.143 | 0.097 | 1.10E-05 | M3 | FABP2 |
| 129 | 1.19E-09 | 0.721048862 | 0.223 | 0.177 | 3.50E-05 | M3 | GLTSCR2 |
| 130 | 1.21E-09 | 0.55622143 | 0.343 | 0.509 | 3.58E-05 | M3 | JUP |
| 131 | 1.21E-08 | 0.76232288 | 0.152 | 0.106 | 0.000355811 | M3 | KRT20 |
| 132 | 8.96E-08 | 0.738412857 | 0.578 | 0.635 | 0.00264186 | M3 | FABP5 |
| 133 | 4.37E-07 | 0.651930006 | 0.211 | 0.176 | 0.01288322 | M3 | ATPIF1 |
| 134 | 8.30E-07 | 0.443842845 | 0.236 | 0.189 | 0.024493137 | M3 | USMG5 |
| 135 | 1.33E-06 | 1.103691294 | 0.156 | 0.121 | 0.039080573 | M3 | MT1H |
| 136 | 1.65E-06 | 0.498006715 | 0.12 | 0.085 | 0.048586562 | M3 | NPW |
| 137 | 1.85E-06 | 1.434569078 | 0.3 | 0.282 | 0.054663534 | M3 | PPP1R1B |
| 138 | 2.62E-06 | 0.598190795 | 0.139 | 0.104 | 0.077182203 | M3 | ANPEP |
| 139 | 2.77E-06 | 0.739867314 | 0.355 | 0.331 | 0.081769695 | M3 | MT1E |
| 140 | 5.78E-06 | 1.03091256 | 0.224 | 0.323 | 0.170527528 | M3 | RF00100.4 |
| 141 | 8.54E-06 | 0.756735018 | 0.495 | 0.536 | 0.25199773 | M3 | SOX4 |
| 142 | 1.00E-05 | 0.505744797 | 0.224 | 0.189 | 0.295838582 | M3 | C14orf2 |
| 143 | 8.19E-05 | 0.419875509 | 0.212 | 0.183 | 1 | M3 | C11orf31 |
| 144 | 0.000124961 | 0.414426924 | 0.49 | 0.669 | 1 | M3 | ASS1 |
| 145 | 0.003149441 | 0.632938445 | 0.116 | 0.099 | 1 | M3 | KLK1 |
| 146 | 0.008749685 | 0.69815094 | 0.18 | 0.241 | 1 | M3 | MT1F |
| 147 | 4.85E-161 | 1.269683952 | 0.597 | 0.116 | 1.43E-156 | M4 | FXYD4 |
| 148 | 3.07E-152 | 0.887048216 | 0.655 | 0.141 | 9.05E-148 | M4 | RTEL1-TNFRSF6B |
| 149 | 3.38E-146 | 1.091110729 | 0.717 | 0.185 | 9.98E-142 | M4 | AC023043.1 |
| 150 | 2.50E-136 | 1.106171557 | 0.752 | 0.203 | 7.37E-132 | M4 | KLK8 |
| 151 | 1.83E-132 | 0.961326759 | 0.711 | 0.176 | 5.41E-128 | M4 | FGL1 |
| 152 | 1.97E-128 | 0.880786351 | 0.726 | 0.193 | 5.82E-124 | M4 | RIMKLB |
| 153 | 2.44E-126 | 0.910770114 | 0.707 | 0.216 | 7.18E-122 | M4 | TM7SF2 |
| 154 | 1.92E-121 | 1.554283888 | 0.942 | 0.618 | 5.67E-117 | M4 | SLPI |
| 155 | 7.36E-121 | 0.974155646 | 0.816 | 0.271 | 2.17E-116 | M4 | RHOV |
| 156 | 1.08E-110 | 0.815478862 | 0.82 | 0.331 | 3.19E-106 | M4 | C18orf32 |
| 157 | 3.59E-102 | 1.138806486 | 0.692 | 0.197 | 1.06E-97 | M4 | INSL4 |
| 158 | 1.04E-101 | 0.899722178 | 0.842 | 0.356 | 3.07E-97 | M4 | PRSS22 |
| 159 | 8.47E-101 | 1.020719143 | 0.859 | 0.427 | 2.50E-96 | M4 | NAXE |
| 160 | 5.01E-98 | 1.134574286 | 0.835 | 0.371 | 1.48E-93 | M4 | CFD |
| 161 | 1.23E-95 | 0.946295972 | 0.315 | 0.053 | 3.64E-91 | M4 | S100A12 |
| 162 | 5.77E-93 | 1.522226621 | 0.792 | 0.332 | 1.70E-88 | M4 | AQP3 |
| 163 | 8.19E-93 | 1.228738384 | 0.884 | 0.597 | 2.42E-88 | M4 | GPX1P1 |
| 164 | 6.73E-90 | 0.926117312 | 0.814 | 0.383 | 1.99E-85 | M4 | CD320 |
| 165 | 4.63E-83 | 0.910687057 | 0.854 | 0.43 | 1.36E-78 | M4 | MYDGF |

| | | | | | | | |
|---|---|---|---|---|---|---|---|
| 166 | 1.01E-82 | 0.826534544 | 0.775 | 0.313 | 2.99E-78 | M4 | PI3 |
| 167 | 1.58E-79 | 1.038862112 | 0.961 | 0.653 | 4.67E-75 | M4 | S100A14 |
| 168 | 6.87E-75 | 1.291598591 | 0.929 | 0.653 | 2.03E-70 | M4 | CD55 |
| 169 | 2.64E-66 | 0.888222531 | 0.797 | 0.422 | 7.79E-62 | M4 | FAM207A |
| 170 | 8.40E-66 | 1.057617747 | 0.996 | 0.838 | 2.48E-61 | M4 | CSTB |
| 171 | 6.15E-62 | 1.47010296 | 0.857 | 0.506 | 1.81E-57 | M4 | KRT17 |
| 172 | 1.35E-59 | 1.041566025 | 0.897 | 0.537 | 3.97E-55 | M4 | PHLDA2 |
| 173 | 1.48E-57 | 1.246523298 | 0.606 | 0.265 | 4.37E-53 | M4 | SFTA2 |
| 174 | 1.09E-54 | 0.802156533 | 0.998 | 0.997 | 3.22E-50 | M4 | S100A6 |
| 175 | 3.11E-54 | 0.990483957 | 0.863 | 0.477 | 9.17E-50 | M4 | S100A9 |
| 176 | 4.20E-52 | 0.901905433 | 0.831 | 0.515 | 1.24E-47 | M4 | NR4A1 |
| 177 | 4.14E-51 | 0.868158831 | 0.246 | 0.058 | 1.22E-46 | M4 | KRT6B |
| 178 | 5.25E-51 | 1.608137479 | 0.632 | 0.281 | 1.55E-46 | M4 | SERPINB3 |
| 179 | 5.48E-51 | 1.553414877 | 0.319 | 0.091 | 1.62E-46 | M4 | S100A2 |
| 180 | 1.70E-49 | 0.937407325 | 0.842 | 0.499 | 5.02E-45 | M4 | AREG |
| 181 | 5.30E-49 | 0.913349389 | 0.807 | 0.487 | 1.56E-44 | M4 | SERPINA1 |
| 182 | 2.39E-46 | 0.966871075 | 0.869 | 0.505 | 7.05E-42 | M4 | ANXA1 |
| 183 | 2.56E-46 | 1.065156804 | 0.522 | 0.213 | 7.55E-42 | M4 | MSMB |
| 184 | 1.22E-45 | 1.333880856 | 0.824 | 0.564 | 3.60E-41 | M4 | CXCL3 |
| 185 | 2.63E-45 | 1.050602258 | 0.882 | 0.605 | 7.76E-41 | M4 | FDPS |
| 186 | 6.82E-44 | 1.012604942 | 0.961 | 0.831 | 2.01E-39 | M4 | S100P |
| 187 | 1.65E-42 | 0.876318568 | 0.876 | 0.521 | 4.87E-38 | M4 | SFN |
| 188 | 2.99E-38 | 1.042861837 | 0.737 | 0.438 | 8.81E-34 | M4 | CXCL1 |
| 189 | 4.61E-37 | 1.883381778 | 0.54 | 0.29 | 1.36E-32 | M4 | NTS |
| 190 | 6.19E-36 | 1.053036977 | 0.452 | 0.189 | 1.83E-31 | M4 | FGB |
| 191 | 2.18E-34 | 0.833392275 | 0.869 | 0.616 | 6.42E-30 | M4 | TM4SF1 |
| 192 | 4.64E-34 | 0.873229362 | 0.989 | 0.958 | 1.37E-29 | M4 | KRT18 |
| 193 | 1.48E-32 | 1.673834443 | 0.625 | 0.33 | 4.37E-28 | M4 | S100A8 |
| 194 | 1.90E-26 | 0.902161458 | 0.893 | 0.698 | 5.61E-22 | M4 | HSPA1A |
| 195 | 2.42E-17 | 1.176353709 | 0.428 | 0.228 | 7.13E-13 | M4 | TCN1 |
| 196 | 3.02E-10 | 1.013389971 | 0.328 | 0.194 | 8.90E-06 | M4 | FGG |
| 197 | 0 | 2.407649178 | 0.945 | 0.305 | 0 | M5 | S100A9 |
| 198 | 0 | 2.105259697 | 0.752 | 0.108 | 0 | M5 | NTS |
| 199 | 0 | 2.103242082 | 0.49 | 0.075 | 0 | M5 | FGG |
| 200 | 0 | 2.018221103 | 0.714 | 0.122 | 0 | M5 | SERPINB3 |
| 201 | 0 | 2.002583723 | 0.739 | 0.216 | 0 | M5 | VIM |
| 202 | 0 | 1.968499457 | 0.922 | 0.354 | 0 | M5 | KRT17 |
| 203 | 0 | 1.839913913 | 0.683 | 0.201 | 0 | M5 | S100A8 |
| 204 | 0 | 1.771068183 | 0.407 | 0.05 | 0 | M5 | GNLY |
| 205 | 0 | 1.695029008 | 0.472 | 0.079 | 0 | M5 | CCL5 |
| 206 | 0 | 1.302067155 | 0.899 | 0.57 | 0 | M5 | CD55 |
| 207 | 0 | 1.121130235 | 0.906 | 0.624 | 0 | M5 | HSPA1A |
| 208 | 1.52E-305 | 1.85082548 | 0.472 | 0.088 | 4.48E-301 | M5 | FGB |
| 209 | 3.11E-285 | 1.940515631 | 0.438 | 0.078 | 9.18E-281 | M5 | FGA |

| | | | | | | | |
|---|---|---|---|---|---|---|---|
| 210 | 6.36E-254 | 1.569490256 | 0.706 | 0.418 | 1.87E-249 | M5 | SCD |
| 211 | 6.79E-250 | 1.078838444 | 0.838 | 0.518 | 2.00E-245 | M5 | GPX1P1 |
| 212 | 1.71E-243 | 1.174003613 | 0.744 | 0.404 | 5.03E-239 | M5 | SERPINA1 |
| 213 | 2.36E-242 | 2.233679339 | 0.476 | 0.137 | 6.96E-238 | M5 | TCN1 |
| 214 | 2.30E-241 | 1.525584259 | 0.528 | 0.179 | 6.79E-237 | M5 | SRGN |
| 215 | 5.41E-238 | 1.10481294 | 0.447 | 0.092 | 1.60E-233 | M5 | SPARC |
| 216 | 2.53E-206 | 1.471325255 | 0.425 | 0.118 | 7.46E-202 | M5 | FGL1 |
| 217 | 4.76E-202 | 1.449064492 | 0.555 | 0.228 | 1.40E-197 | M5 | LGALS1 |
| 218 | 1.89E-185 | 1.341814857 | 0.575 | 0.269 | 5.58E-181 | M5 | AQP3 |
| 219 | 1.08E-182 | 1.310924805 | 0.441 | 0.137 | 3.20E-178 | M5 | INSL4 |
| 220 | 6.86E-160 | 1.190169745 | 0.243 | 0.036 | 2.02E-155 | M5 | GZMB |
| 221 | 9.72E-160 | 1.349148776 | 0.477 | 0.188 | 2.87E-155 | M5 | CXCL8 |
| 222 | 3.18E-157 | 1.323679982 | 0.333 | 0.086 | 9.37E-153 | M5 | CCL4 |
| 223 | 2.86E-154 | 1.383869451 | 0.387 | 0.134 | 8.42E-150 | M5 | GLIPR1 |
| 224 | 2.52E-146 | 0.957842074 | 0.302 | 0.065 | 7.42E-142 | M5 | COL4A1 |
| 225 | 4.59E-145 | 1.507471695 | 0.502 | 0.28 | 1.35E-140 | M5 | PLK2 |
| 226 | 9.73E-139 | 1.220265654 | 0.256 | 0.05 | 2.87E-134 | M5 | RGS5 |
| 227 | 8.05E-132 | 1.118514176 | 0.273 | 0.063 | 2.37E-127 | M5 | MGP |
| 228 | 5.02E-129 | 1.305751899 | 0.5 | 0.283 | 1.48E-124 | M5 | FURIN |
| 229 | 6.01E-126 | 1.055758312 | 0.618 | 0.387 | 1.77E-121 | M5 | CXCL1 |
| 230 | 8.22E-126 | 1.126340299 | 0.306 | 0.084 | 2.42E-121 | M5 | CALD1 |
| 231 | 2.47E-125 | 1.014787874 | 0.568 | 0.328 | 7.28E-121 | M5 | CFD |
| 232 | 1.10E-122 | 1.259475228 | 0.483 | 0.274 | 3.26E-118 | M5 | CTSL |
| 233 | 2.09E-110 | 1.09275568 | 0.521 | 0.314 | 6.17E-106 | M5 | MUC5B |
| 234 | 9.24E-103 | 1.017698325 | 0.24 | 0.065 | 2.72E-98 | M5 | VTN |
| 235 | 1.47E-101 | 1.043398289 | 0.264 | 0.081 | 4.35E-97 | M5 | TYROBP |
| 236 | 2.79E-95 | 1.009779839 | 0.448 | 0.246 | 8.22E-91 | M5 | RHOV |
| 237 | 1.09E-91 | 1.075973849 | 0.362 | 0.171 | 3.20E-87 | M5 | RIMKLB |
| 238 | 5.62E-89 | 1.129266733 | 0.433 | 0.258 | 1.66E-84 | M5 | PHLDA1 |
| 239 | 2.06E-84 | 1.11244836 | 0.445 | 0.285 | 6.08E-80 | M5 | AL135905.2 |
| 240 | 1.70E-77 | 1.010426735 | 0.309 | 0.139 | 5.02E-73 | M5 | NNMT |
| 241 | 1.65E-57 | 1.118947967 | 0.249 | 0.113 | 4.87E-53 | M5 | GOS2 |
| 242 | 2.19E-50 | 0.98949623 | 0.396 | 0.291 | 6.45E-46 | M5 | SGK1 |
| 243 | 2.26E-48 | 0.984920524 | 0.207 | 0.089 | 6.68E-44 | M5 | ACTA2 |
| 244 | 5.24E-37 | 0.97581293 | 0.414 | 0.358 | 1.55E-32 | M5 | NR4A2 |
| 245 | 4.25E-27 | 1.000875241 | 0.323 | 0.255 | 1.25E-22 | M5 | PDK4 |
| 246 | 3.83E-24 | 1.132629927 | 0.516 | 0.571 | 1.13E-19 | M5 | ODC1 |
| 247 | 6.25E-284 | 3.014140273 | 0.742 | 0.105 | 1.84E-279 | M6 | COL1A2 |
| 248 | 4.53E-283 | 2.754749194 | 0.745 | 0.102 | 1.33E-278 | M6 | COL3A1 |
| 249 | 8.65E-238 | 2.437590694 | 0.636 | 0.086 | 2.55E-233 | M6 | FN1 |
| 250 | 6.80E-227 | 2.798753506 | 0.814 | 0.169 | 2.01E-222 | M6 | SPARC |
| 251 | 5.64E-225 | 3.396688613 | 0.768 | 0.162 | 1.66E-220 | M6 | COL1A1 |
| 252 | 2.12E-205 | 1.791604218 | 0.596 | 0.081 | 6.26E-201 | M6 | COL6A2 |
| 253 | 3.75E-197 | 2.148521938 | 0.653 | 0.111 | 1.10E-192 | M6 | COL4A1 |

| | | | | | | | |
|---|---|---|---|---|---|---|---|
| 254 | 1.37E-191 | 1.905799539 | 0.63 | 0.102 | 4.04E-187 | M6 | SPARCL1 |
| 255 | 6.54E-187 | 1.628919785 | 0.272 | 0.014 | 1.93E-182 | M6 | CCL11 |
| 256 | 1.08E-182 | 1.544059655 | 0.387 | 0.035 | 3.19E-178 | M6 | MMP2 |
| 257 | 2.47E-181 | 1.946678835 | 0.673 | 0.125 | 7.28E-177 | M6 | CALD1 |
| 258 | 2.08E-172 | 2.056509718 | 0.499 | 0.068 | 6.14E-168 | M6 | LUM |
| 259 | 3.62E-157 | 1.557965147 | 0.427 | 0.054 | 1.07E-152 | M6 | SERPINF1 |
| 260 | 1.71E-155 | 1.604624921 | 0.467 | 0.065 | 5.03E-151 | M6 | BGN |
| 261 | 1.82E-152 | 2.006928221 | 0.579 | 0.114 | 5.37E-148 | M6 | THBS1 |
| 262 | 6.27E-152 | 2.013103015 | 0.464 | 0.065 | 1.85E-147 | M6 | DCN |
| 263 | 2.18E-143 | 2.246862912 | 0.564 | 0.111 | 6.42E-139 | M6 | TAGLN |
| 264 | 3.80E-136 | 1.834463127 | 0.897 | 0.35 | 1.12E-131 | M6 | VIM |
| 265 | 3.34E-135 | 1.528045986 | 0.372 | 0.046 | 9.84E-131 | M6 | COL6A3 |
| 266 | 7.78E-131 | 1.077431669 | 0.229 | 0.015 | 2.29E-126 | M6 | POSTN |
| 267 | 1.47E-123 | 2.134330467 | 0.501 | 0.095 | 4.33E-119 | M6 | A2M |
| 268 | 1.68E-123 | 1.837106601 | 0.622 | 0.172 | 4.95E-119 | M6 | CTGF |
| 269 | 6.41E-115 | 1.94424728 | 0.9 | 0.542 | 1.89E-110 | M6 | TIMP1 |
| 270 | 1.43E-109 | 1.38526085 | 0.441 | 0.081 | 4.22E-105 | M6 | SERPINE1 |
| 271 | 7.78E-109 | 1.647731415 | 0.493 | 0.106 | 2.29E-104 | M6 | ACTA2 |
| 272 | 4.44E-108 | 1.515214681 | 0.424 | 0.08 | 1.31E-103 | M6 | C1S |
| 273 | 1.21E-104 | 1.589844729 | 0.788 | 0.305 | 3.58E-100 | M6 | LGALS1 |
| 274 | 1.12E-92 | 1.419264927 | 0.484 | 0.109 | 3.29E-88 | M6 | MGP |
| 275 | 4.20E-87 | 1.512620814 | 0.404 | 0.089 | 1.24E-82 | M6 | RARRES2 |
| 276 | 8.65E-79 | 1.256739665 | 0.384 | 0.086 | 2.55E-74 | M6 | COL6A1 |
| 277 | 1.80E-75 | 1.356051044 | 0.401 | 0.1 | 5.32E-71 | M6 | CYR61 |
| 278 | 8.53E-73 | 0.994609958 | 0.252 | 0.038 | 2.52E-68 | M6 | MAP1B |
| 279 | 1.35E-65 | 1.525392993 | 0.39 | 0.099 | 3.99E-61 | M6 | RGS5 |
| 280 | 8.75E-64 | 1.58814367 | 0.55 | 0.218 | 2.58E-59 | M6 | MYL9 |
| 281 | 1.69E-56 | 1.251455714 | 0.312 | 0.077 | 4.98E-52 | M6 | TIMP3 |
| 282 | 1.30E-53 | 1.226235959 | 0.352 | 0.103 | 3.83E-49 | M6 | SPON2 |
| 283 | 1.50E-52 | 1.156965922 | 0.372 | 0.11 | 4.43E-48 | M6 | SERPING1 |
| 284 | 2.43E-49 | 0.947831495 | 0.289 | 0.071 | 7.17E-45 | M6 | SOD3 |
| 285 | 5.05E-48 | 1.628700024 | 0.278 | 0.072 | 1.49E-43 | M6 | SERPINA3 |
| 286 | 7.16E-36 | 1.027543726 | 0.828 | 0.664 | 2.11E-31 | M6 | CD55 |
| 287 | 1.99E-31 | 0.959948388 | 0.708 | 0.496 | 5.88E-27 | M6 | SCD |
| 288 | 5.53E-30 | 1.046599233 | 0.395 | 0.185 | 1.63E-25 | M6 | ICAM1 |
| 289 | 2.40E-29 | 1.072884591 | 0.378 | 0.174 | 7.08E-25 | M6 | TIMP2 |
| 290 | 8.53E-24 | 1.599676784 | 0.117 | 0.025 | 2.52E-19 | M6 | SFRP2 |
| 291 | 3.97E-21 | 1.060366322 | 0.447 | 0.282 | 1.17E-16 | M6 | TPM2 |
| 292 | 4.29E-21 | 1.049809802 | 0.33 | 0.166 | 1.27E-16 | M6 | ACSL4 |
| 293 | 9.30E-21 | 1.124720087 | 0.347 | 0.183 | 2.74E-16 | M6 | GEM |
| 294 | 1.47E-20 | 0.943485195 | 0.395 | 0.22 | 4.35E-16 | M6 | PRSS23 |
| 295 | 1.71E-14 | 0.949142575 | 0.433 | 0.316 | 5.03E-10 | M6 | HMGCS1 |
| 296 | 0.002168965 | 1.029468184 | 0.08 | 0.143 | 1 | M6 | PGC |

Supplementary Table S3 DEGs of 9 non-malignant epithelium subclusters

| | p_val | avg_log2FC | pct.1 | pct.2 | p_val_adj | cluster | gene |
|---|---|---|---|---|---|---|---|
| 1 | 0 | 5.501156904 | 0.854 | 0.057 | 0 | PMC | MUCL3 |
| 2 | 0 | 5.259691328 | 0.999 | 0.314 | 0 | PMC | GKN1 |
| 3 | 0 | 4.149060154 | 0.998 | 0.598 | 0 | PMC | MUC5AC |
| 4 | 0.00E+00 | 3.722427122 | 0.988 | 0.332 | 0.00E+00 | PMC | GKN2 |
| 5 | 0.00E+00 | 3.709989608 | 0.91 | 0.32 | 0.00E+00 | PMC | CAPN8 |
| 6 | 0.00E+00 | 3.233673531 | 0.793 | 0.056 | 0.00E+00 | PMC | CEACAM5 |
| 7 | 0.00E+00 | 3.177162137 | 1 | 0.653 | 0.00E+00 | PMC | TFF1 |
| 8 | 0.00E+00 | 3.058567506 | 0.751 | 0.043 | 0.00E+00 | PMC | PSAPL1 |
| 9 | 0.00E+00 | 3.013214041 | 0.868 | 0.23 | 0.00E+00 | PMC | SYTL2 |
| 10 | 0.00E+00 | 2.829748142 | 0.812 | 0.219 | 0.00E+00 | PMC | RASEF |
| 11 | 0.00E+00 | 2.806484668 | 0.669 | 0.012 | 0.00E+00 | PMC | SLC5A5 |
| 12 | 0.00E+00 | 2.701852544 | 0.78 | 0.083 | 0.00E+00 | PMC | LGALS9C |
| 13 | 0.00E+00 | 2.639791287 | 0.989 | 0.574 | 0.00E+00 | PMC | PSCA |
| 14 | 0.00E+00 | 2.506656701 | 0.447 | 0.001 | 0.00E+00 | PMC | BNIP5 |
| 15 | 0.00E+00 | 2.384064048 | 0.774 | 0.176 | 0.00E+00 | PMC | B4GALNT3 |
| 16 | 0.00E+00 | 2.333939135 | 0.545 | 0.001 | 0.00E+00 | PMC | PLAAT2 |
| 17 | 0.00E+00 | 2.329638516 | 0.681 | 0.063 | 0.00E+00 | PMC | LGALS9B |
| 18 | 0.00E+00 | 2.320367239 | 0.828 | 0.304 | 0.00E+00 | PMC | LINC01133 |
| 19 | 0.00E+00 | 2.31519893 | 0.625 | 0.103 | 0.00E+00 | PMC | DDX60 |
| 20 | 0.00E+00 | 2.309318451 | 0.542 | 0.16 | 0.00E+00 | PMC | ANKRD36C |
| 21 | 0.00E+00 | 2.277214183 | 0.783 | 0.229 | 0.00E+00 | PMC | VILL |
| 22 | 0.00E+00 | 2.247010027 | 0.705 | 0.294 | 0.00E+00 | PMC | RNF213 |
| 23 | 0.00E+00 | 2.233740136 | 0.864 | 0.554 | 0.00E+00 | PMC | AHNAK |
| 24 | 0.00E+00 | 2.227986076 | 0.423 | 0.035 | 0.00E+00 | PMC | DUOX2 |
| 25 | 0.00E+00 | 2.203523938 | 0.977 | 0.749 | 0.00E+00 | PMC | MUC1 |
| 26 | 0.00E+00 | 2.145287069 | 0.879 | 0.522 | 0.00E+00 | PMC | MLPH |
| 27 | 0.00E+00 | 2.121256307 | 0.979 | 0.665 | 0.00E+00 | PMC | IFI27 |
| 28 | 0.00E+00 | 2.034330471 | 0.835 | 0.299 | 0.00E+00 | PMC | CYP3A5 |
| 29 | 0.00E+00 | 2.032327277 | 0.596 | 0.082 | 0.00E+00 | PMC | RFLNA |
| 30 | 0.00E+00 | 2.025062158 | 0.888 | 0.483 | 0.00E+00 | PMC | GSN |
| 31 | 0.00E+00 | 3.981576048 | 0.999 | 0.523 | 0.00E+00 | pre-PMC | RPS29 |
| 32 | 0.00E+00 | 3.578325587 | 0.985 | 0.345 | 0.00E+00 | pre-PMC | RPS20 |
| 33 | 0.00E+00 | 3.537214815 | 0.975 | 0.042 | 0.00E+00 | pre-PMC | GNB2L1 |
| 34 | 0.00E+00 | 3.432285774 | 0.998 | 0.556 | 0.00E+00 | pre-PMC | RPL27A |
| 35 | 0.00E+00 | 3.327478694 | 0.987 | 0.375 | 0.00E+00 | pre-PMC | RPL36A |
| 36 | 0.00E+00 | 3.287626429 | 0.999 | 0.667 | 0.00E+00 | pre-PMC | RPL13A |
| 37 | 0.00E+00 | 3.250567564 | 0.993 | 0.519 | 0.00E+00 | pre-PMC | RPL37A |
| 38 | 0.00E+00 | 3.185678176 | 0.973 | 0.426 | 0.00E+00 | pre-PMC | RPL7 |
| 39 | 0.00E+00 | 3.180602665 | 0.989 | 0.511 | 0.00E+00 | pre-PMC | RPL21 |
| 40 | 0.00E+00 | 3.177585341 | 0.996 | 0.524 | 0.00E+00 | pre-PMC | RPL31 |
| 41 | 0.00E+00 | 3.121266464 | 0.999 | 0.785 | 0.00E+00 | pre-PMC | RPL41 |
| 42 | 0.00E+00 | 3.014875059 | 0.984 | 0.352 | 0.00E+00 | pre-PMC | RPS10 |

| | | | | | | | |
|---|---|---|---|---|---|---|---|
| 43 | 0.00E+00 | 2.879213572 | 0.983 | 0.072 | 0.00E+00 | pre-PMC | ATP5E |
| 44 | 0.00E+00 | 2.848690813 | 0.966 | 0.362 | 0.00E+00 | pre-PMC | RPL23 |
| 45 | 0.00E+00 | 2.801755584 | 0.971 | 0.052 | 0.00E+00 | pre-PMC | ATP5I |
| 46 | 0.00E+00 | 2.800330979 | 0.992 | 0.559 | 0.00E+00 | pre-PMC | RPL23A |
| 47 | 0.00E+00 | 2.750287367 | 0.993 | 0.529 | 0.00E+00 | pre-PMC | RPL38 |
| 48 | 0.00E+00 | 2.744466353 | 0.935 | 0.031 | 0.00E+00 | pre-PMC | ATP5G2 |
| 49 | 0.00E+00 | 2.737446825 | 1 | 0.804 | 0.00E+00 | pre-PMC | RPS27 |
| 50 | 0 | 2.713716027 | 0.982 | 0.457 | 0 | pre-PMC | RPL27 |
| 51 | 0 | 2.706075722 | 0.999 | 0.709 | 0 | pre-PMC | RPLP2 |
| 52 | 0 | 2.702529138 | 0.839 | 0.113 | 0 | pre-PMC | LIPF |
| 53 | 0 | 2.598541504 | 0.989 | 0.546 | 0 | pre-PMC | RPS11 |
| 54 | 0 | 2.566401066 | 0.99 | 0.543 | 0 | pre-PMC | RPS16 |
| 55 | 0 | 2.524412934 | 0.994 | 0.642 | 0 | pre-PMC | RPL35 |
| 56 | 0 | 2.496275666 | 0.883 | 0.025 | 0 | pre-PMC | ATP5O |
| 57 | 0 | 2.458668555 | 0.94 | 0.053 | 0 | pre-PMC | UQCR11-1 |
| 58 | 0 | 2.439831916 | 0.994 | 0.681 | 0 | pre-PMC | RPL39 |
| 59 | 0 | 2.43009451 | 0.995 | 0.62 | 0 | pre-PMC | RPL34 |
| 60 | 0 | 2.428624899 | 0.93 | 0.042 | 0 | pre-PMC | ATP5L |
| 61 | 0 | 9.381766862 | 0.977 | 0.102 | 0 | Chief cell | PGA3 |
| 62 | 0 | 5.414393925 | 0.675 | 0.003 | 0 | Chief cell | PGA5 |
| 63 | 0 | 3.285568178 | 0.823 | 0.053 | 0 | Chief cell | PGA4 |
| 64 | 1.283E-249 | 3.479472818 | 0.997 | 0.262 | 4.22E-245 | Chief cell | LIPF |
| 65 | 4.7914E-170 | 3.221063449 | 1 | 0.524 | 1.5759E-165 | Chief cell | PGC |
| 66 | 1.4451E-143 | 1.315497809 | 0.639 | 0.114 | 4.7531E-139 | Chief cell | LTF |
| 67 | 2.7592E-137 | 1.366495416 | 0.846 | 0.225 | 9.0754E-133 | Chief cell | REG1A |
| 68 | 1.4984E-126 | 1.940916015 | 0.882 | 0.326 | 4.9283E-122 | Chief cell | MT1F |
| 69 | 2.8062E-122 | 1.317904373 | 0.757 | 0.179 | 9.23E-118 | Chief cell | LRRC75A-AS1 |
| 70 | 5.3645E-120 | 1.66989506 | 0.82 | 0.245 | 1.7644E-115 | Chief cell | MT1M |
| 71 | 9.72E-118 | 1.253496396 | 0.715 | 0.185 | 3.20E-113 | Chief cell | PDIA2 |
| 72 | 6.36E-113 | 1.26139527 | 0.515 | 0.099 | 2.09E-108 | Chief cell | HLA-DPA1 |
| 73 | 1.36E-112 | 1.323561378 | 0.151 | 0.007 | 4.48E-108 | Chief cell | CHIA |
| 74 | 5.74E-98 | 1.385656868 | 0.951 | 0.434 | 1.89E-93 | Chief cell | CXCL17 |
| 75 | 1.29E-91 | 1.399464679 | 0.692 | 0.225 | 4.25E-87 | Chief cell | AZGP1 |
| 76 | 4.21E-91 | 1.283050243 | 0.816 | 0.288 | 1.38E-86 | Chief cell | ATP5MC2 |
| 77 | 1.97E-88 | 2.096586373 | 0.885 | 0.453 | 6.47E-84 | Chief cell | CD74 |
| 78 | 1.37E-86 | 1.572232518 | 0.492 | 0.114 | 4.49E-82 | Chief cell | MIA |
| 79 | 1.39E-84 | 1.170440972 | 0.866 | 0.383 | 4.58E-80 | Chief cell | DCXR |
| 80 | 1.08E-80 | 1.913437427 | 0.97 | 0.602 | 3.54E-76 | Chief cell | MT1G |
| 81 | 1.81E-78 | 1.399025022 | 0.816 | 0.365 | 5.94E-74 | Chief cell | RACK1 |
| 82 | 1.97E-70 | 1.671668442 | 0.879 | 0.454 | 6.49E-66 | Chief cell | MT1X |
| 83 | 2.77E-64 | 1.922561839 | 0.748 | 0.402 | 9.10E-60 | Chief cell | HSPA1A |
| 84 | 1.17E-60 | 1.217320897 | 0.436 | 0.116 | 3.85E-56 | Chief cell | ADH1C |
| 85 | 2.57E-57 | 1.181331755 | 0.692 | 0.28 | 8.44E-53 | Chief cell | HLA-DRA |
| 86 | 9.49E-55 | 1.229550863 | 0.639 | 0.257 | 3.12E-50 | Chief cell | HLA-DRB1 |

| | | | | | | | |
|---|---|---|---|---|---|---|---|
| 87 | 1.25E-47 | 1.454943751 | 0.931 | 0.672 | 4.12E-43 | Chief cell | MT2A |
| 88 | 9.20E-47 | 1.111932995 | 0.954 | 0.685 | 3.03E-42 | Chief cell | MT1E |
| 89 | 2.13E-42 | 1.119011414 | 0.734 | 0.413 | 6.99E-38 | Chief cell | HSPA1B |
| 90 | 7.57E-07 | 1.348558995 | 0.239 | 0.132 | 2.49E-02 | Chief cell | PRR4 |
| 91 | 0.00E+00 | 4.635784654 | 0.938 | 0.076 | 0.00E+00 | Neck cell | LTF |
| 92 | 0.00E+00 | 4.588943743 | 0.953 | 0.145 | 0.00E+00 | Neck cell | BPIFB1 |
| 93 | 0.00E+00 | 4.587671877 | 0.707 | 0.048 | 0.00E+00 | Neck cell | REG3A |
| 94 | 0.00E+00 | 4.327166743 | 0.985 | 0.196 | 0.00E+00 | Neck cell | REG1A |
| 95 | 0.00E+00 | 3.733842264 | 0.994 | 0.2 | 0.00E+00 | Neck cell | MUC6 |
| 96 | 0.00E+00 | 3.684187815 | 0.856 | 0.085 | 0.00E+00 | Neck cell | PRR4 |
| 97 | 0.00E+00 | 3.586263126 | 0.804 | 0.05 | 0.00E+00 | Neck cell | MTCO1P12 |
| 98 | 0.00E+00 | 3.477327777 | 0.948 | 0.299 | 0.00E+00 | Neck cell | MSMB |
| 99 | 0.00E+00 | 3.368776007 | 0.925 | 0.266 | 0.00E+00 | Neck cell | LCN2 |
| 100 | 0.00E+00 | 3.294468007 | 0.826 | 0.049 | 0.00E+00 | Neck cell | GP2 |
| 101 | 0 | 3.213451996 | 0.879 | 0.14 | 0 | Neck cell | SERPINA1 |
| 102 | 0 | 2.903605873 | 0.953 | 0.12 | 0 | Neck cell | AQP5 |
| 103 | 0 | 2.881267087 | 0.794 | 0.047 | 0 | Neck cell | AC139749.1 |
| 104 | 0 | 2.719381625 | 0.824 | 0.087 | 0 | Neck cell | C6orf58 |
| 105 | 0 | 2.342179384 | 0.693 | 0.01 | 0 | Neck cell | SERPINA3 |
| 106 | 0 | 2.315704325 | 0.708 | 0.033 | 0 | Neck cell | TCN1 |
| 107 | 0 | 2.242591848 | 0.869 | 0.21 | 0 | Neck cell | ZG16B |
| 108 | 0 | 2.233205137 | 0.95 | 0.273 | 0 | Neck cell | NME2 |
| 109 | 0 | 2.175272891 | 0.673 | 0.091 | 0 | Neck cell | CXCL2 |
| 110 | 0 | 1.894458933 | 0.865 | 0.153 | 0 | Neck cell | LRRC75A-AS1 |
| 111 | 0 | 1.841640693 | 0.75 | 0.123 | 0 | Neck cell | C2CD4B |
| 112 | 0 | 1.764595695 | 0.622 | 0.029 | 0 | Neck cell | S100A1 |
| 113 | 1.5565E-296 | 3.081107541 | 1 | 0.509 | 5.1196E-292 | Neck cell | PGC |
| 114 | 4.7324E-290 | 3.295561186 | 1 | 0.771 | 1.5565E-285 | Neck cell | LYZ |
| 115 | 1.2353E-273 | 1.848925239 | 0.985 | 0.233 | 4.063E-269 | Neck cell | MT-RNR2 |
| 116 | 1.5306E-268 | 1.783408503 | 0.92 | 0.264 | 5.0342E-264 | Neck cell | ATP5MC2 |
| 117 | 2.13E-265 | 2.147350751 | 0.985 | 0.433 | 7.0058E-261 | Neck cell | CD74 |
| 118 | 6.1209E-254 | 1.852950406 | 0.961 | 0.341 | 2.0132E-249 | Neck cell | RACK1 |
| 119 | 4.3224E-176 | 2.233333848 | 0.32 | 0.035 | 1.4217E-171 | Neck cell | AC090498.1 |
| 120 | 4.80314E-84 | 2.438978587 | 0.241 | 0.044 | 1.5798E-79 | Neck cell | OLFM4 |
| 121 | 0 | 3.947541129 | 0.883 | 0.156 | 0 | SPEM | TFF3 |
| 122 | 0 | 3.835237501 | 0.781 | 0.103 | 0 | SPEM | C6orf58 |
| 123 | 0 | 3.295718642 | 0.993 | 0.214 | 0 | SPEM | MUC6 |
| 124 | 0 | 2.835057077 | 0.633 | 0.064 | 0 | SPEM | IGKV1-5 |
| 125 | 0 | 2.717085227 | 0.706 | 0.072 | 0 | SPEM | AC020656.1 |
| 126 | 0 | 2.285589686 | 0.85 | 0.14 | 0 | SPEM | AQP5 |
| 127 | 0 | 2.271078423 | 0.474 | 0.022 | 0 | SPEM | SOD3 |
| 128 | 0 | 1.692759837 | 0.377 | 0.018 | 0 | SPEM | LINC00632 |
| 129 | 1.2019E-251 | 2.105564329 | 0.613 | 0.086 | 3.9531E-247 | SPEM | EEF1A1P5 |
| 130 | 2.3561E-242 | 2.165421129 | 0.586 | 0.076 | 7.7495E-238 | SPEM | GP2 |

| | | | | | | | |
|---|---|---|---|---|---|---|---|
| 131 | 2.3211E-238 | 1.789078162 | 0.663 | 0.112 | 7.6342E-234 | SPEM | PPP1R1B |
| 132 | 1.458E-235 | 1.860354461 | 0.584 | 0.078 | 4.7954E-231 | SPEM | JCHAIN |
| 133 | 2.8939E-205 | 2.09849725 | 0.521 | 0.078 | 9.5184E-201 | SPEM | RARRES2 |
| 134 | 1.0814E-196 | 2.801756322 | 0.995 | 0.775 | 3.5569E-192 | SPEM | LYZ |
| 135 | 2.8828E-194 | 1.986596619 | 0.885 | 0.288 | 9.4817E-190 | SPEM | NME2 |
| 136 | 2.7904E-189 | 1.858958971 | 0.948 | 0.352 | 9.1778E-185 | SPEM | RACK1 |
| 137 | 6.1918E-183 | 1.761481232 | 0.743 | 0.172 | 2.0366E-178 | SPEM | LRRC75A-AS1 |
| 138 | 3.2816E-179 | 1.801197676 | 0.633 | 0.135 | 1.0793E-174 | SPEM | C16orf89 |
| 139 | 6.4348E-179 | 2.164452373 | 0.349 | 0.034 | 2.1165E-174 | SPEM | IGHV4-34 |
| 140 | 6.4068E-176 | 1.720950467 | 0.92 | 0.25 | 2.1073E-171 | SPEM | MT-RNR2 |
| 141 | 5.2227E-174 | 2.850622088 | 0.252 | 0.016 | 1.7178E-169 | SPEM | SCGB3A1 |
| 142 | 1.9734E-172 | 1.694324486 | 0.87 | 0.278 | 6.4906E-168 | SPEM | ATP5MC2 |
| 143 | 1.9112E-135 | 1.733352093 | 0.753 | 0.222 | 6.2862E-131 | SPEM | REG1A |
| 144 | 3.1729E-134 | 1.701070034 | 0.923 | 0.502 | 1.0436E-129 | SPEM | S100A10 |
| 145 | 9.4697E-124 | 2.08884927 | 0.314 | 0.041 | 3.1147E-119 | SPEM | IGHA1 |
| 146 | 5.757E-113 | 1.782706099 | 0.945 | 0.657 | 1.8935E-108 | SPEM | GOLM1 |
| 147 | 7.60E-98 | 2.018652474 | 0.641 | 0.234 | 2.50E-93 | SPEM | ZG16B |
| 148 | 3.64E-56 | 1.727903197 | 0.945 | 0.854 | 1.20E-51 | SPEM | TFF2 |
| 149 | 6.73E-43 | 2.424852516 | 0.175 | 0.035 | 2.21E-38 | SPEM | IGLV1-44 |
| 150 | 4.11E-34 | 1.790646987 | 0.185 | 0.048 | 1.35E-29 | SPEM | AC090498.1 |
| 151 | 0.00E+00 | 7.744223194 | 0.953 | 0.029 | 0.00E+00 | IM | SPINK4 |
| 152 | 0.00E+00 | 6.562675741 | 0.899 | 0.019 | 0.00E+00 | IM | MUC2 |
| 153 | 0.00E+00 | 5.229259248 | 0.838 | 0.027 | 0.00E+00 | IM | ITLN1 |
| 154 | 0.00E+00 | 5.198867601 | 0.595 | 0.004 | 0.00E+00 | IM | CLCA1 |
| 155 | 0.00E+00 | 4.522103776 | 0.588 | 0.018 | 0.00E+00 | IM | ZG16 |
| 156 | 0.00E+00 | 2.824855199 | 0.797 | 0.029 | 0.00E+00 | IM | KLK1 |
| 157 | 0.00E+00 | 1.626272965 | 0.622 | 0.018 | 0.00E+00 | IM | HEPACAM2 |
| 158 | 3.58E-252 | 5.476184288 | 0.939 | 0.104 | 1.18E-247 | IM | REG4 |
| 159 | 1.11E-244 | 1.99780831 | 0.743 | 0.059 | 3.66E-240 | IM | LRRC26 |
| 160 | 1.31E-221 | 1.607018417 | 0.669 | 0.045 | 4.32E-217 | IM | FABP2 |
| 161 | 2.11E-197 | 2.217947559 | 0.838 | 0.089 | 6.94E-193 | IM | CLDN3 |
| 162 | 5.93E-191 | 5.114808298 | 1 | 0.178 | 1.95E-186 | IM | TFF3 |
| 163 | 2.51E-156 | 1.566672265 | 0.541 | 0.044 | 8.26E-152 | IM | MUC4 |
| 164 | 3.22E-122 | 1.700574611 | 0.838 | 0.14 | 1.06E-117 | IM | CAMK2N1 |
| 165 | 3.64E-111 | 1.557917695 | 0.791 | 0.136 | 1.20E-106 | IM | CLDN7 |
| 166 | 1.97E-108 | 1.994115464 | 0.615 | 0.086 | 6.47E-104 | IM | HES6 |
| 167 | 6.28E-101 | 1.966649638 | 0.953 | 0.226 | 2.07E-96 | IM | CLDN4 |
| 168 | 2.40E-94 | 2.02171273 | 0.818 | 0.218 | 7.88E-90 | IM | ST6GALNAC1 |
| 169 | 6.13E-89 | 3.792273138 | 0.973 | 0.453 | 2.02E-84 | IM | FCGBP |
| 170 | 1.00E-83 | 1.981496071 | 0.98 | 0.27 | 3.29E-79 | IM | MT-RNR2 |
| 171 | 7.04E-81 | 1.616610901 | 0.973 | 0.267 | 2.31E-76 | IM | MT-RNR1 |
| 172 | 1.17E-79 | 2.48965076 | 0.709 | 0.16 | 3.85E-75 | IM | WFDC2 |
| 173 | 1.91E-74 | 1.7235603 | 0.784 | 0.19 | 6.30E-70 | IM | LRRC75A-AS1 |
| 174 | 1.03E-67 | 1.900420938 | 0.892 | 0.372 | 3.40E-63 | IM | CDC42EP5 |

| | | | | | | | |
|---|---|---|---|---|---|---|---|
| 175 | 4.53E-61 | 2.834733391 | 0.946 | 0.681 | 1.49E-56 | IM | LGALS4 |
| 176 | 1.08E-55 | 1.782788133 | 0.851 | 0.311 | 3.55E-51 | IM | NR4A1 |
| 177 | 1.07E-52 | 1.755279632 | 0.946 | 0.746 | 3.52E-48 | IM | SH3BGRL3 |
| 178 | 1.09E-52 | 2.834267021 | 0.142 | 0.009 | 3.57E-48 | IM | PRSS2 |
| 179 | 1.43E-22 | 1.732502524 | 0.284 | 0.068 | 4.71E-18 | IM | IGKV4-1 |
| 180 | 2.35E-10 | 2.022438835 | 0.216 | 0.077 | 7.74E-06 | IM | FP671120.1 |
| 181 | 0.00E+00 | 6.036722927 | 0.436 | 0.016 | 0.00E+00 | Enterocyte | APOA4 |
| 182 | 0.00E+00 | 5.731430216 | 0.464 | 0.04 | 0.00E+00 | Enterocyte | APOA1 |
| 183 | 0.00E+00 | 5.460841632 | 0.661 | 0.026 | 0.00E+00 | Enterocyte | FABP1 |
| 184 | 0.00E+00 | 4.907357086 | 0.783 | 0.02 | 0.00E+00 | Enterocyte | ANPEP |
| 185 | 0.00E+00 | 4.509950023 | 0.336 | 0.009 | 0.00E+00 | Enterocyte | APOC3 |
| 186 | 0.00E+00 | 4.430649397 | 0.723 | 0.033 | 0.00E+00 | Enterocyte | PRAP1 |
| 187 | 0.00E+00 | 4.393130123 | 0.416 | 0.01 | 0.00E+00 | Enterocyte | RBP2 |
| 188 | 0.00E+00 | 4.04359656 | 0.618 | 0.027 | 0.00E+00 | Enterocyte | ALDOB |
| 189 | 0.00E+00 | 3.772352578 | 0.576 | 0.03 | 0.00E+00 | Enterocyte | PCK1 |
| 190 | 0.00E+00 | 3.668494007 | 0.65 | 0.079 | 0.00E+00 | Enterocyte | RF00100.4 |
| 191 | 0.00E+00 | 3.388848019 | 0.978 | 0.219 | 0.00E+00 | Enterocyte | MT-RNR1 |
| 192 | 0.00E+00 | 3.277022923 | 0.484 | 0.019 | 0.00E+00 | Enterocyte | FABP2 |
| 193 | 0.00E+00 | 3.008398445 | 0.387 | 0.005 | 0.00E+00 | Enterocyte | APOB |
| 194 | 0.00E+00 | 2.898338191 | 0.991 | 0.221 | 0.00E+00 | Enterocyte | MT-RNR2 |
| 195 | 0.00E+00 | 2.750436251 | 0.402 | 0.003 | 0.00E+00 | Enterocyte | CYP3A4 |
| 196 | 0.00E+00 | 2.290281761 | 0.385 | 0.004 | 0.00E+00 | Enterocyte | C3orf85 |
| 197 | 0.00E+00 | 2.273160726 | 0.317 | 0.002 | 0.00E+00 | Enterocyte | TMPRSS15 |
| 198 | 0.00E+00 | 2.245408089 | 0.536 | 0.065 | 0 | Enterocyte | CLDN3 |
| 199 | 0.00E+00 | 2.19627005 | 0.361 | 0.009 | 0 | Enterocyte | SLC6A19 |
| 200 | 2.3266E-306 | 2.593244942 | 0.428 | 0.043 | 7.6526E-302 | Enterocyte | SLC5A1 |
| 201 | 2.6087E-306 | 2.20069978 | 0.613 | 0.107 | 8.5802E-302 | Enterocyte | CLDN7 |
| 202 | 4.0651E-256 | 5.430493073 | 0.411 | 0.049 | 1.3371E-251 | Enterocyte | FP236383.1 |
| 203 | 7.8691E-243 | 3.735316795 | 0.407 | 0.05 | 2.5882E-238 | Enterocyte | FP671120.1 |
| 204 | 1.9494E-221 | 2.371386088 | 0.569 | 0.124 | 6.4119E-217 | Enterocyte | MUC17 |
| 205 | 9.8554E-218 | 2.711960409 | 0.345 | 0.039 | 3.2415E-213 | Enterocyte | RPL41P1 |
| 206 | 7.4465E-184 | 2.706532233 | 0.643 | 0.23 | 2.4492E-179 | Enterocyte | CDHR5 |
| 207 | 6.3937E-144 | 2.771950672 | 0.362 | 0.069 | 2.103E-139 | Enterocyte | IGKV1-5 |
| 208 | 1.3816E-125 | 2.384441054 | 0.485 | 0.161 | 4.5441E-121 | Enterocyte | SELENOP |
| 209 | 4.3502E-108 | 2.449173031 | 0.297 | 0.06 | 1.4308E-103 | Enterocyte | IGLV2-14 |
| 210 | 3.5171E-105 | 2.199582642 | 0.616 | 0.308 | 1.15682E-100 | Enterocyte | ADIRF |
| 211 | 0 | 6.254053638 | 0.737 | 0.041 | 0 | Endocrine | CHGA |
| 212 | 0 | 5.977284667 | 0.905 | 0.022 | 0 | Endocrine | PCSK1N |
| 213 | 0 | 3.7510905 | 0.658 | 0.004 | 0 | Endocrine | CHGB |
| 214 | 0 | 3.720223205 | 0.77 | 0.03 | 0 | Endocrine | SCG5 |
| 215 | 0 | 3.611843779 | 0.473 | 0.012 | 0 | Endocrine | TTR |
| 216 | 0 | 3.433401196 | 0.724 | 0.036 | 0 | Endocrine | TUBA1A |
| 217 | 0 | 3.181139149 | 0.642 | 0.006 | 0 | Endocrine | CPE |
| 218 | 0 | 2.913444161 | 0.498 | 0.003 | 0 | Endocrine | CRYBA2 |

| | | | | | | | |
|---|---|---|---|---|---|---|---|
| 219 | 0 | 2.728281921 | 0.313 | 0.001 | 0 | Endocrine | SHISAL2B |
| 220 | 0 | 2.667991503 | 0.79 | 0.073 | 0 | Endocrine | MS4A8 |
| 221 | 0 | 2.590946362 | 0.654 | 0.015 | 0 | Endocrine | CACNA1A |
| 222 | 0 | 2.383143523 | 0.671 | 0.002 | 0 | Endocrine | SCG3 |
| 223 | 0 | 2.360349726 | 0.601 | 0.011 | 0 | Endocrine | SCGN |
| 224 | 0 | 2.273883501 | 0.543 | 0.001 | 0 | Endocrine | FEV |
| 225 | 0 | 2.094881068 | 0.305 | 0.004 | 0 | Endocrine | HAP1 |
| 226 | 3.1643E-273 | 2.889700407 | 0.671 | 0.065 | 1.0408E-268 | Endocrine | ERO1B |
| 227 | 1.4196E-248 | 2.354797365 | 0.267 | 0.007 | 4.6691E-244 | Endocrine | CES1 |
| 228 | 7.2242E-248 | 2.434787583 | 0.23 | 0.004 | 2.3761E-243 | Endocrine | TPH1 |
| 229 | 1.9261E-127 | 4.181022651 | 0.226 | 0.013 | 6.3351E-123 | Endocrine | RBP4 |
| 230 | 3.064E-112 | 2.100494303 | 0.535 | 0.103 | 1.0078E-107 | Endocrine | PAM |
| 231 | 5.0666E-107 | 2.757755243 | 0.626 | 0.15 | 1.6665E-102 | Endocrine | DEPP1 |
| 232 | 1.73308E-71 | 8.574841741 | 0.321 | 0.055 | 5.70027E-67 | Endocrine | GAST |
| 233 | 2.76E-66 | 2.046432431 | 0.844 | 0.442 | 9.07E-62 | Endocrine | MDK |
| 234 | 9.83E-59 | 2.045411008 | 0.523 | 0.154 | 3.23E-54 | Endocrine | C2CD4B |
| 235 | 9.52E-55 | 2.081318195 | 0.519 | 0.16 | 3.13E-50 | Endocrine | CDKN1C |
| 236 | 9.87E-55 | 2.229851924 | 0.798 | 0.481 | 3.25E-50 | Endocrine | BTG2 |
| 237 | 6.98E-46 | 2.153053661 | 0.646 | 0.31 | 2.30E-41 | Endocrine | NR4A1 |
| 238 | 3.32E-35 | 2.057830434 | 0.494 | 0.197 | 1.09E-30 | Endocrine | RASD1 |
| 239 | 9.00E-12 | 2.043771641 | 0.111 | 0.031 | 2.96E-07 | Endocrine | IGLV2-11 |
| 240 | 4.97E-11 | 2.047339961 | 0.3 | 0.174 | 1.63E-06 | Endocrine | SCGB2A1 |
| 241 | 0.00E+00 | 6.246903703 | 0.964 | 0.015 | 0.00E+00 | Parietal cell | GIF |
| 242 | 0.00E+00 | 6.014580794 | 0.936 | 0.007 | 0.00E+00 | Parietal cell | ATP4B |
| 243 | 0.00E+00 | 4.227695542 | 0.791 | 0.004 | 0.00E+00 | Parietal cell | ATP4A |
| 244 | 0.00E+00 | 3.627040158 | 0.709 | 0.029 | 0.00E+00 | Parietal cell | GOS2 |
| 245 | 0.00E+00 | 2.885106962 | 0.573 | 0.002 | 0.00E+00 | Parietal cell | CPA2 |
| 246 | 2.44E-144 | 3.431257333 | 0.945 | 0.159 | 8.03E-140 | Parietal cell | SLC25A4 |
| 247 | 1.52E-143 | 6.175694292 | 1 | 0.195 | 5.00E-139 | Parietal cell | CKB |
| 248 | 2.77E-135 | 4.169220008 | 0.973 | 0.19 | 9.11E-131 | Parietal cell | IDH2 |
| 249 | 6.14E-114 | 4.090169712 | 0.973 | 0.241 | 2.02E-109 | Parietal cell | KCNE2 |
| 250 | 7.35E-108 | 3.874810028 | 0.982 | 0.269 | 2.42E-103 | Parietal cell | LDHB |
| 251 | 7.66E-101 | 3.145163788 | 0.909 | 0.223 | 2.51946E-96 | Parietal cell | MPC1 |
| 252 | 2.02228E-93 | 3.148263708 | 0.955 | 0.277 | 6.65149E-89 | Parietal cell | GSTA1 |
| 253 | 4.19036E-92 | 3.35298438 | 0.918 | 0.264 | 1.37825E-87 | Parietal cell | MDH1 |
| 254 | 1.50849E-79 | 3.065817536 | 0.882 | 0.274 | 4.96157E-75 | Parietal cell | GHITM |
| 255 | 3.48984E-76 | 3.257687589 | 0.973 | 0.459 | 1.14784E-71 | Parietal cell | AKR7A3 |
| 256 | 4.02706E-74 | 3.708033369 | 0.982 | 0.543 | 1.32454E-69 | Parietal cell | SLC25A5 |
| 257 | 6.25436E-73 | 3.19288588 | 0.982 | 0.559 | 2.05712E-68 | Parietal cell | NDUFA4 |
| 258 | 1.53692E-70 | 3.445323257 | 0.955 | 0.432 | 5.05508E-66 | Parietal cell | ALDH1A1 |
| 259 | 1.49E-68 | 3.105350369 | 0.927 | 0.397 | 4.89E-64 | Parietal cell | CYCS |
| 260 | 1.01E-65 | 2.887617309 | 0.973 | 0.538 | 3.33E-61 | Parietal cell | UQCRB |
| 261 | 1.78E-64 | 2.966115168 | 0.936 | 0.507 | 5.85E-60 | Parietal cell | COX5A |
| 262 | 6.06E-64 | 3.118348662 | 0.973 | 0.652 | 1.99E-59 | Parietal cell | UQCRQ |

| | | | | | | | |
|---|---|---|---|---|---|---|---|
| 263 | 3.96E-61 | 3.168596888 | 0.945 | 0.601 | 1.30E-56 | Parietal cell | COX7A2 |
| 264 | 7.05E-60 | 3.080306037 | 0.927 | 0.58 | 2.32E-55 | Parietal cell | COX7B |
| 265 | 7.45E-41 | 2.953950644 | 0.591 | 0.182 | 2.45E-36 | Parietal cell | ATP5A1 |
| 266 | 3.00E-39 | 3.320370649 | 0.609 | 0.21 | 9.86E-35 | Parietal cell | ATP5J |
| 267 | 1.91E-38 | 2.911285564 | 0.591 | 0.195 | 6.28E-34 | Parietal cell | ATP5G1 |
| 268 | 1.79E-37 | 3.782683524 | 0.609 | 0.223 | 5.89E-33 | Parietal cell | ATP5G3 |
| 269 | 4.48E-37 | 3.3249527 | 0.582 | 0.196 | 1.47E-32 | Parietal cell | ATP5B |
| 270 | 2.11E-31 | 3.063541002 | 0.6 | 0.243 | 6.94E-27 | Parietal cell | USMG5 |

Supplementary Table S4 DEGs of T/NK subclusters

| | p_val | avg_log2FC | pct.1 | pct.2 | p_val_adj | cluster | gene |
|---|---|---|---|---|---|---|---|
| 1 | 0 | 0.882161129 | 0.157 | 0.001 | 0 | ILC | PCDH9 |
| 2 | 8.20E-306 | 0.968271076 | 0.142 | 0.002 | 2.42E-301 | ILC | TNFSF11 |
| 3 | 5.59E-300 | 1.658469101 | 0.331 | 0.015 | 1.65E-295 | ILC | KIT |
| 4 | 2.82E-249 | 1.362099874 | 0.394 | 0.026 | 8.31E-245 | ILC | KRT81 |
| 5 | 1.21E-213 | 1.763697545 | 0.63 | 0.077 | 3.57E-209 | ILC | FCER1G |
| 6 | 2.96E-205 | 1.612757812 | 0.484 | 0.049 | 8.72E-201 | ILC | KRT86 |
| 7 | 9.56E-171 | 0.941361472 | 0.331 | 0.027 | 2.82E-166 | ILC | FES |
| 8 | 6.70E-157 | 1.657185698 | 0.669 | 0.112 | 1.97E-152 | ILC | TYROBP |
| 9 | 6.91E-149 | 0.875041368 | 0.299 | 0.025 | 2.04E-144 | ILC | B3GNT7 |
| 10 | 2.95E-141 | 1.048108797 | 0.362 | 0.039 | 8.71E-137 | ILC | SH2D1B |
| 11 | 8.58E-141 | 1.322568403 | 0.378 | 0.043 | 2.53E-136 | ILC | GSN |
| 12 | 2.33E-135 | 0.890487603 | 0.343 | 0.036 | 6.88E-131 | ILC | CXXC5 |
| 13 | 5.81E-117 | 1.494978637 | 0.701 | 0.166 | 1.71E-112 | ILC | TMIGD2 |
| 14 | 9.50E-114 | 2.689128142 | 0.717 | 0.192 | 2.80E-109 | ILC | AREG |
| 15 | 1.20E-97 | 1.077280629 | 0.283 | 0.034 | 3.53E-93 | ILC | SPINK2 |
| 16 | 2.39E-83 | 1.698738332 | 0.303 | 0.046 | 7.06E-79 | ILC | LST1 |
| 17 | 4.37E-81 | 1.957109807 | 0.602 | 0.164 | 1.29E-76 | ILC | XCL1 |
| 18 | 6.57E-65 | 1.087704892 | 0.374 | 0.078 | 1.94E-60 | ILC | CD160 |
| 19 | 9.16E-65 | 0.878340528 | 0.429 | 0.099 | 2.70E-60 | ILC | TXK |
| 20 | 2.00E-63 | 1.021635497 | 0.26 | 0.043 | 5.91E-59 | ILC | TLE1 |
| 21 | 3.58E-63 | 1.025920697 | 0.307 | 0.056 | 1.06E-58 | ILC | TRDC |
| 22 | 4.12E-62 | 1.334978727 | 0.732 | 0.29 | 1.21E-57 | ILC | MAP3K8 |
| 23 | 8.78E-59 | 1.124360447 | 0.453 | 0.12 | 2.59E-54 | ILC | ENTPD1 |
| 24 | 1.36E-58 | 0.986765107 | 0.5 | 0.139 | 4.02E-54 | ILC | SLC16A3 |
| 25 | 2.17E-57 | 0.885269334 | 0.382 | 0.089 | 6.39E-53 | ILC | ZBTB16 |
| 26 | 3.71E-57 | 1.731562869 | 0.626 | 0.22 | 1.10E-52 | ILC | XCL2 |
| 27 | 1.31E-56 | 1.177280672 | 0.236 | 0.039 | 3.85E-52 | ILC | IL4I1 |
| 28 | 1.48E-53 | 0.912718932 | 0.35 | 0.08 | 4.37E-49 | ILC | MAFF |
| 29 | 1.80E-49 | 1.361253146 | 0.661 | 0.283 | 5.32E-45 | ILC | LDLRAD4 |
| 30 | 1.68E-46 | 0.930757824 | 0.598 | 0.219 | 4.94E-42 | ILC | TNFRSF18 |
| 31 | 4.36E-36 | 1.04017271 | 0.543 | 0.225 | 1.29E-31 | ILC | TIPARP |
| 32 | 4.86E-36 | 0.881527141 | 0.343 | 0.101 | 1.43E-31 | ILC | KLRC2 |
| 33 | 5.51E-36 | 1.232514586 | 0.764 | 0.47 | 1.62E-31 | ILC | NR4A2 |

| | | | | | | | |
|---|---|---|---|---|---|---|---|
| 34 | 8.34E-36 | 0.891936884 | 0.449 | 0.163 | 2.46E-31 | ILC | NSMCE1 |
| 35 | 3.72E-35 | 1.129279788 | 0.681 | 0.349 | 1.10E-30 | ILC | ZNF331 |
| 36 | 8.51E-35 | 0.887705633 | 0.602 | 0.261 | 2.51E-30 | ILC | SCML4 |
| 37 | 4.09E-34 | 1.153271267 | 0.52 | 0.206 | 1.21E-29 | ILC | NR4A1 |
| 38 | 1.17E-32 | 0.89384877 | 0.378 | 0.129 | 3.47E-28 | ILC | CAT |
| 39 | 4.44E-31 | 0.895351893 | 0.528 | 0.235 | 1.31E-26 | ILC | MPG |
| 40 | 1.86E-29 | 1.283282806 | 0.823 | 0.577 | 5.49E-25 | ILC | ZFP36L1 |
| 41 | 5.64E-26 | 0.908114342 | 0.685 | 0.414 | 1.66E-21 | ILC | FAM177A1 |
| 42 | 3.50E-25 | 0.882204498 | 0.898 | 0.598 | 1.03E-20 | ILC | FOS |
| 43 | 2.00E-24 | 0.903220057 | 0.52 | 0.259 | 5.90E-20 | ILC | PFKFB3 |
| 44 | 2.39E-24 | 0.997195551 | 0.846 | 0.819 | 7.06E-20 | ILC | CD7 |
| 45 | 5.01E-23 | 1.002387698 | 0.291 | 0.104 | 1.48E-18 | ILC | CD83 |
| 46 | 1.73E-16 | 0.88979258 | 0.701 | 0.449 | 5.09E-12 | ILC | ID2 |
| 47 | 6.87E-15 | 0.940138211 | 0.165 | 0.054 | 2.03E-10 | ILC | SOX4 |
| 48 | 3.33E-14 | 1.139473889 | 0.417 | 0.234 | 9.83E-10 | ILC | IGKV1-5 |
| 49 | 5.45E-11 | 0.906142972 | 0.528 | 0.409 | 1.61E-06 | ILC | EEF1A1P5 |
| 50 | 1.29E-10 | 1.062002124 | 0.189 | 0.08 | 3.81E-06 | ILC | MIR24-2 |
| 51 | 0 | 3.47263987 | 0.922 | 0.041 | 0 | NK1 | FCER1G |
| 52 | 0 | 3.2155761 | 0.943 | 0.078 | 0 | NK1 | TYROBP |
| 53 | 0 | 3.082327184 | 0.895 | 0.516 | 0 | NK1 | GZMA |
| 54 | 0 | 3.039164374 | 0.686 | 0.19 | 0 | NK1 | GNLY |
| 55 | 0 | 2.46789827 | 0.775 | 0.17 | 0 | NK1 | AREG |
| 56 | 0 | 2.189269264 | 0.612 | 0.031 | 0 | NK1 | TRDC |
| 57 | 0 | 2.098981493 | 0.717 | 0.081 | 0 | NK1 | KIR2DL4 |
| 58 | 0 | 1.938661086 | 0.567 | 0.028 | 0 | NK1 | KRT86 |
| 59 | 0 | 1.861008115 | 0.992 | 0.81 | 0 | NK1 | CD7 |
| 60 | 0 | 1.844043201 | 0.791 | 0.195 | 0 | NK1 | TNFRSF18 |
| 61 | 0 | 1.830241957 | 0.917 | 0.508 | 0 | NK1 | GSTP1 |
| 62 | 0 | 1.806118087 | 0.711 | 0.145 | 0 | NK1 | TMIGD2 |
| 63 | 0 | 1.76292385 | 0.892 | 0.469 | 0 | NK1 | CD247 |
| 64 | 0 | 1.748729287 | 0.679 | 0.192 | 0 | NK1 | CLIC3 |
| 65 | 0 | 1.66414472 | 0.857 | 0.314 | 0 | NK1 | HOPX |
| 66 | 0 | 1.621683576 | 0.567 | 0.016 | 0 | NK1 | SH2D1B |
| 67 | 0 | 1.610583011 | 0.433 | 0.01 | 0 | NK1 | KRT81 |
| 68 | 0 | 1.604267744 | 0.492 | 0.061 | 0 | NK1 | CD160 |
| 69 | 0 | 1.526189394 | 0.566 | 0.036 | 0 | NK1 | LAT2 |
| 70 | 0 | 1.458617264 | 0.615 | 0.12 | 0 | NK1 | SLC16A3 |
| 71 | 0 | 1.417740695 | 0.549 | 0.103 | 0 | NK1 | ENTPD1 |
| 72 | 0 | 1.296746367 | 0.678 | 0.169 | 0 | NK1 | MATK |
| 73 | 0 | 1.268419727 | 0.563 | 0.079 | 0 | NK1 | TXK |
| 74 | 0 | 1.263835388 | 0.384 | 0.011 | 0 | NK1 | B3GNT7 |
| 75 | 0 | 1.106835009 | 0.379 | 0.025 | 0 | NK1 | IRF8 |
| 76 | 0 | 1.086544236 | 0.455 | 0.056 | 0 | NK1 | MCTP2 |
| 77 | 0 | 1.067691119 | 0.377 | 0.019 | 0 | NK1 | FGR |

| | | | | | | | |
|---|---|---|---|---|---|---|---|
| 78 | 0 | 1.055212435 | 0.322 | 0.003 | 0 | NK1 | ADGRG3 |
| 79 | 7.85E-294 | 1.382738395 | 0.748 | 0.242 | 2.32E-289 | NK1 | IL2RB |
| 80 | 1.72E-272 | 1.377352268 | 0.764 | 0.272 | 5.09E-268 | NK1 | MAP3K8 |
| 81 | 1.93E-267 | 1.400769429 | 0.693 | 0.212 | 5.70E-263 | NK1 | CEBPD |
| 82 | 3.12E-242 | 1.077705049 | 0.425 | 0.087 | 9.19E-238 | NK1 | KLRC2 |
| 83 | 7.50E-229 | 1.30882876 | 0.878 | 0.412 | 2.21E-224 | NK1 | PRF1 |
| 84 | 2.69E-215 | 1.143777399 | 0.881 | 0.46 | 7.95E-211 | NK1 | CD63 |
| 85 | 1.86E-212 | 1.09131562 | 0.349 | 0.068 | 5.50E-208 | NK1 | GEM |
| 86 | 4.03E-202 | 1.198419255 | 0.719 | 0.277 | 1.19E-197 | NK1 | FOSL2 |
| 87 | 4.14E-202 | 1.062491172 | 0.961 | 0.546 | 1.22E-197 | NK1 | NKG7 |
| 88 | 1.75E-193 | 1.222267446 | 0.951 | 0.663 | 5.15E-189 | NK1 | CCL5 |
| 89 | 2.23E-173 | 1.201068606 | 0.703 | 0.286 | 6.57E-169 | NK1 | KLRD1 |
| 90 | 1.62E-172 | 1.194025303 | 0.351 | 0.083 | 4.78E-168 | NK1 | PLPP1 |
| 91 | 3.06E-172 | 1.350266213 | 0.747 | 0.34 | 9.03E-168 | NK1 | CTSW |
| 92 | 3.59E-169 | 1.084294343 | 0.745 | 0.334 | 1.06E-164 | NK1 | CAPG |
| 93 | 2.87E-152 | 1.15100238 | 0.852 | 0.422 | 8.46E-148 | NK1 | GZMB |
| 94 | 4.72E-148 | 1.323497225 | 0.489 | 0.168 | 1.39E-143 | NK1 | KLRC1 |
| 95 | 2.28E-136 | 1.100479464 | 0.623 | 0.27 | 6.73E-132 | NK1 | LDLRAD4 |
| 96 | 1.39E-118 | 1.103322282 | 0.53 | 0.205 | 4.11E-114 | NK1 | GZMK |
| 97 | 1.14E-98 | 1.076794433 | 0.337 | 0.109 | 3.35E-94 | NK1 | CCL3 |
| 98 | 7.64E-84 | 1.142485324 | 0.479 | 0.212 | 2.25E-79 | NK1 | XCL2 |
| 99 | 1.88E-75 | 1.152223912 | 0.383 | 0.159 | 5.56E-71 | NK1 | XCL1 |
| 100 | 1.07E-11 | 1.160699372 | 0.326 | 0.232 | 3.17E-07 | NK1 | IGKV1-5 |
| 101 | 0 | 2.908811895 | 0.58 | 0.005 | 0 | NK2 | FGFBP2 |
| 102 | 0 | 2.65056364 | 0.647 | 0.026 | 0 | NK2 | FCGR3A |
| 103 | 0 | 2.574956524 | 0.938 | 0.188 | 0 | NK2 | GNLY |
| 104 | 0 | 2.301544382 | 0.999 | 0.551 | 0 | NK2 | NKG7 |
| 105 | 0 | 2.28484436 | 0.689 | 0.099 | 0 | NK2 | TYROBP |
| 106 | 0 | 1.984461144 | 0.889 | 0.305 | 0 | NK2 | GZMH |
| 107 | 0 | 1.769812021 | 0.414 | 0.013 | 0 | NK2 | KLRF1 |
| 108 | 0 | 1.544695233 | 0.374 | 0.009 | 0 | NK2 | S1PR5 |
| 109 | 0 | 1.380111209 | 0.155 | 0.004 | 0 | NK2 | MYOM2 |
| 110 | 0 | 1.054885534 | 0.242 | 0.009 | 0 | NK2 | PRSS23 |
| 111 | 0 | 0.924845047 | 0.171 | 0.001 | 0 | NK2 | CX3CR1 |
| 112 | 6.62E-266 | 1.73615463 | 0.839 | 0.287 | 1.95E-261 | NK2 | KLRD1 |
| 113 | 8.25E-246 | 1.882916132 | 0.902 | 0.418 | 2.43E-241 | NK2 | PRF1 |
| 114 | 2.69E-237 | 1.372693384 | 0.399 | 0.067 | 7.92E-233 | NK2 | PLEK |
| 115 | 7.74E-220 | 1.607934977 | 0.957 | 0.601 | 2.28E-215 | NK2 | CST7 |
| 116 | 6.81E-218 | 0.951618847 | 0.239 | 0.025 | 2.01E-213 | NK2 | C1orf21 |
| 117 | 2.01E-210 | 1.489796685 | 0.935 | 0.426 | 5.93E-206 | NK2 | GZMB |
| 118 | 4.24E-204 | 1.594716136 | 0.82 | 0.343 | 1.25E-199 | NK2 | CTSW |
| 119 | 2.29E-182 | 1.001062242 | 0.26 | 0.036 | 6.76E-178 | NK2 | CD300A |
| 120 | 1.68E-176 | 1.657749844 | 0.732 | 0.267 | 4.96E-172 | NK2 | KLF2 |
| 121 | 2.26E-176 | 0.970327059 | 0.153 | 0.012 | 6.67E-172 | NK2 | KIR3DL1 |

| | | | | | | | |
|---|---|---|---|---|---|---|---|
| 122 | 2.59E-158 | 0.994413989 | 0.252 | 0.038 | 7.65E-154 | NK2 | ADRB2 |
| 123 | 4.00E-155 | 1.898945642 | 0.524 | 0.176 | 1.18E-150 | NK2 | SPON2 |
| 124 | 2.33E-142 | 0.854658694 | 0.155 | 0.016 | 6.88E-138 | NK2 | KIR2DS4 |
| 125 | 5.13E-141 | 0.955037376 | 0.252 | 0.043 | 1.51E-136 | NK2 | ADGRG1 |
| 126 | 4.66E-133 | 1.343797889 | 0.666 | 0.318 | 1.37E-128 | NK2 | EFHD2 |
| 127 | 1.41E-129 | 1.135929654 | 0.38 | 0.1 | 4.17E-125 | NK2 | PLAC8 |
| 128 | 3.10E-129 | 0.861072013 | 0.242 | 0.042 | 9.13E-125 | NK2 | FCRL6 |
| 129 | 3.36E-112 | 2.402099165 | 0.377 | 0.111 | 9.90E-108 | NK2 | CCL3 |
| 130 | 1.76E-105 | 1.125732442 | 0.906 | 0.525 | 5.18E-101 | NK2 | CCL4 |
| 131 | 1.82E-100 | 1.11303714 | 0.685 | 0.37 | 5.37E-96 | NK2 | LITAF |
| 132 | 3.62E-89 | 1.341320127 | 0.494 | 0.206 | 1.07E-84 | NK2 | CLIC3 |
| 133 | 4.26E-81 | 1.206556063 | 0.658 | 0.413 | 1.26E-76 | NK2 | ITGB2 |
| 134 | 1.81E-78 | 0.984620533 | 0.386 | 0.144 | 5.35E-74 | NK2 | TBX21 |
| 135 | 1.35E-76 | 1.061172306 | 0.494 | 0.225 | 3.99E-72 | NK2 | ZEB2 |
| 136 | 1.53E-73 | 1.106247579 | 0.588 | 0.335 | 4.50E-69 | NK2 | FLNA |
| 137 | 3.71E-69 | 1.216318609 | 0.429 | 0.191 | 1.09E-64 | NK2 | CMC1 |
| 138 | 7.56E-63 | 0.982474449 | 0.54 | 0.298 | 2.23E-58 | NK2 | XBP1 |
| 139 | 4.98E-62 | 0.896844699 | 0.256 | 0.077 | 1.47E-57 | NK2 | FCER1G |
| 140 | 7.70E-62 | 0.905145702 | 0.465 | 0.232 | 2.27E-57 | NK2 | APMAP |
| 141 | 1.99E-61 | 0.903249341 | 0.347 | 0.139 | 5.88E-57 | NK2 | UPP1 |
| 142 | 5.36E-61 | 1.013404318 | 0.501 | 0.251 | 1.58E-56 | NK2 | METRNL |
| 143 | 5.49E-58 | 0.878946729 | 0.689 | 0.482 | 1.62E-53 | NK2 | CD247 |
| 144 | 2.84E-57 | 0.904738121 | 0.406 | 0.188 | 8.38E-53 | NK2 | ITGB1 |
| 145 | 1.70E-56 | 0.863883676 | 0.62 | 0.404 | 5.00E-52 | NK2 | ABHD17A |
| 146 | 1.42E-40 | 1.552123745 | 0.242 | 0.091 | 4.18E-36 | NK2 | HSPA6 |
| 147 | 2.86E-34 | 0.865893088 | 0.419 | 0.243 | 8.44E-30 | NK2 | GADD45B |
| 148 | 1.85E-24 | 1.133253242 | 0.255 | 0.132 | 5.47E-20 | NK2 | RHOB |
| 149 | 2.21E-16 | 0.863564446 | 0.524 | 0.402 | 6.50E-12 | NK2 | HSPA1A |
| 150 | 5.32E-07 | 0.875335416 | 0.263 | 0.197 | 0.015696499 | NK2 | AREG |
| 151 | 0 | 3.562689353 | 0.953 | 0.141 | 0 | CD8_MKI67 | STMN1 |
| 152 | 0 | 3.048340838 | 0.978 | 0.544 | 0 | CD8_MKI67 | TUBA1B |
| 153 | 0 | 2.857990915 | 0.946 | 0.433 | 0 | CD8_MKI67 | TUBB |
| 154 | 0 | 2.497290366 | 0.742 | 0.022 | 0 | CD8_MKI67 | TYMS |
| 155 | 0 | 2.227523468 | 0.966 | 0.505 | 0 | CD8_MKI67 | HMGN2 |
| 156 | 0 | 2.21579281 | 0.698 | 0.02 | 0 | CD8_MKI67 | MKI67 |
| 157 | 0 | 2.195000333 | 0.589 | 0.009 | 0 | CD8_MKI67 | UBE2C |
| 158 | 0 | 1.923945295 | 0.4 | 0.015 | 0 | CD8_MKI67 | HIST1H1B |
| 159 | 0 | 1.802667434 | 0.624 | 0.045 | 0 | CD8_MKI67 | NUSAP1 |
| 160 | 0 | 1.741850229 | 0.555 | 0.013 | 0 | CD8_MKI67 | TOP2A |
| 161 | 0 | 1.734611733 | 0.563 | 0.028 | 0 | CD8_MKI67 | CENPF |
| 162 | 0 | 1.69769852 | 0.543 | 0.006 | 0 | CD8_MKI67 | RRM2 |
| 163 | 0 | 1.544662922 | 0.631 | 0.05 | 0 | CD8_MKI67 | CKS1B |
| 164 | 0 | 1.542276587 | 0.49 | 0.008 | 0 | CD8_MKI67 | ASPM |
| 165 | 0 | 1.535704928 | 0.364 | 0.035 | 0 | CD8_MKI67 | HIST1H2AI |

| | | | | | | | |
|---|---|---|---|---|---|---|---|
| 166 | 0 | 1.52455105 | 0.572 | 0.023 | 0 | CD8_MKI67 | TPX2 |
| 167 | 0 | 1.515639626 | 0.589 | 0.016 | 0 | CD8_MKI67 | TK1 |
| 168 | 0 | 1.480648632 | 0.61 | 0.101 | 0 | CD8_MKI67 | PCNA |
| 169 | 0 | 1.457423696 | 0.568 | 0.023 | 0 | CD8_MKI67 | CDK1 |
| 170 | 0 | 1.451092953 | 0.425 | 0.005 | 0 | CD8_MKI67 | CDC20 |
| 171 | 0 | 1.448247244 | 0.501 | 0.012 | 0 | CD8_MKI67 | PCLAF |
| 172 | 0 | 1.440354354 | 0.602 | 0.06 | 0 | CD8_MKI67 | MCM7 |
| 173 | 0 | 1.440204972 | 0.557 | 0.019 | 0 | CD8_MKI67 | CDKN3 |
| 174 | 0 | 1.38938156 | 0.658 | 0.072 | 0 | CD8_MKI67 | CENPM |
| 175 | 0 | 1.363513205 | 0.608 | 0.012 | 0 | CD8_MKI67 | ZWINT |
| 176 | 0 | 1.360518414 | 0.734 | 0.149 | 0 | CD8_MKI67 | SMC4 |
| 177 | 0 | 1.270434368 | 0.462 | 0.006 | 0 | CD8_MKI67 | CCNB2 |
| 178 | 0 | 1.257881556 | 0.479 | 0.005 | 0 | CD8_MKI67 | BIRC5 |
| 179 | 0 | 1.256836715 | 0.644 | 0.085 | 0 | CD8_MKI67 | TMEM106C |
| 180 | 8.09E-286 | 2.137409565 | 0.964 | 0.519 | 2.39E-281 | CD8_MKI67 | HMGB2 |
| 181 | 2.14E-275 | 1.472667805 | 1 | 0.951 | 6.32E-271 | CD8_MKI67 | GAPDH |
| 182 | 1.87E-267 | 1.982384705 | 0.964 | 0.587 | 5.53E-263 | CD8_MKI67 | H2AFZ |
| 183 | 3.86E-264 | 1.729503245 | 0.829 | 0.272 | 1.14E-259 | CD8_MKI67 | DUT |
| 184 | 1.88E-263 | 1.54009075 | 0.784 | 0.211 | 5.55E-259 | CD8_MKI67 | FABP5 |
| 185 | 2.43E-263 | 1.44602529 | 0.751 | 0.194 | 7.17E-259 | CD8_MKI67 | PTTG1 |
| 186 | 1.71E-241 | 1.583813741 | 0.986 | 0.804 | 5.05E-237 | CD8_MKI67 | HMGB1 |
| 187 | 3.72E-232 | 1.276901487 | 0.75 | 0.201 | 1.10E-227 | CD8_MKI67 | CKS2 |
| 188 | 1.79E-221 | 1.31319483 | 0.781 | 0.237 | 5.27E-217 | CD8_MKI67 | TMPO |
| 189 | 2.89E-218 | 1.262582302 | 0.83 | 0.282 | 8.53E-214 | CD8_MKI67 | RANBP1 |
| 190 | 2.79E-203 | 1.516530995 | 0.921 | 0.479 | 8.24E-199 | CD8_MKI67 | H2AFV |
| 191 | 3.64E-202 | 1.372514311 | 0.969 | 0.615 | 1.07E-197 | CD8_MKI67 | TPI1 |
| 192 | 1.62E-199 | 1.59372772 | 0.767 | 0.263 | 4.77E-195 | CD8_MKI67 | HLA-DRA |
| 193 | 5.91E-170 | 1.408386568 | 0.932 | 0.472 | 1.74E-165 | CD8_MKI67 | MT2A |
| 194 | 3.35E-151 | 1.258714821 | 0.802 | 0.335 | 9.87E-147 | CD8_MKI67 | TUBB4B |
| 195 | 1.49E-147 | 2.754402115 | 0.835 | 0.461 | 4.41E-143 | CD8_MKI67 | HIST1H4C |
| 196 | 6.29E-143 | 1.384720616 | 0.561 | 0.165 | 1.85E-138 | CD8_MKI67 | HIST1H1C |
| 197 | 1.01E-133 | 1.351241157 | 0.827 | 0.402 | 2.97E-129 | CD8_MKI67 | HLA-DRB1 |
| 198 | 7.73E-130 | 1.471093909 | 0.359 | 0.079 | 2.28E-125 | CD8_MKI67 | CXCL13 |
| 199 | 1.51E-122 | 1.442192926 | 0.76 | 0.362 | 4.45E-118 | CD8_MKI67 | UBE2S |
| 200 | 4.05E-74 | 1.668776369 | 0.589 | 0.276 | 1.19E-69 | CD8_MKI67 | HIST1H1E |
| 201 | 0 | 1.791875943 | 0.9 | 0.378 | 0 | CD8_CXCL13 | GZMB |
| 202 | 0 | 1.747187712 | 0.706 | 0.261 | 0 | CD8_CXCL13 | LAG3 |
| 203 | 0 | 1.746461156 | 0.318 | 0.055 | 0 | CD8_CXCL13 | CXCL13 |
| 204 | 0 | 1.711385368 | 0.427 | 0.107 | 0 | CD8_CXCL13 | LINC02446 |
| 205 | 0 | 1.654496679 | 0.779 | 0.26 | 0 | CD8_CXCL13 | GZMH |
| 206 | 0 | 1.628802264 | 0.334 | 0.029 | 0 | CD8_CXCL13 | VCAM1 |
| 207 | 0 | 1.57739018 | 0.529 | 0.169 | 0 | CD8_CXCL13 | GNLY |
| 208 | 0 | 1.482047616 | 0.951 | 0.511 | 0 | CD8_CXCL13 | NKG7 |
| 209 | 0 | 1.452312798 | 0.811 | 0.358 | 0 | CD8_CXCL13 | HLA-DRB1 |

| | | | | | | | |
|---|---|---|---|---|---|---|---|
| 210 | 0 | 1.382582626 | 0.735 | 0.244 | 0 | CD8_CXCL13 | KLRD1 |
| 211 | 0 | 1.329286889 | 0.78 | 0.385 | 0 | CD8_CXCL13 | PRF1 |
| 212 | 0 | 1.322312205 | 0.408 | 0.085 | 0 | CD8_CXCL13 | HAVCR2 |
| 213 | 0 | 1.204813107 | 0.379 | 0.074 | 0 | CD8_CXCL13 | KIR2DL4 |
| 214 | 0 | 1.197644161 | 0.983 | 0.632 | 0 | CD8_CXCL13 | CCL5 |
| 215 | 0 | 1.17229413 | 0.786 | 0.401 | 0 | CD8_CXCL13 | HLA-DPA1 |
| 216 | 0 | 1.083056177 | 0.657 | 0.295 | 0 | CD8_CXCL13 | DUSP4 |
| 217 | 0 | 1.013602469 | 0.734 | 0.367 | 0 | CD8_CXCL13 | HLA-DPB1 |
| 218 | 0 | 0.977602161 | 0.775 | 0.356 | 0 | CD8_CXCL13 | CD8A |
| 219 | 0 | 0.877371395 | 0.902 | 0.482 | 0 | CD8_CXCL13 | GZMA |
| 220 | 1.83E-305 | 0.941028216 | 0.695 | 0.305 | 5.39E-301 | CD8_CXCL13 | CD8B |
| 221 | 1.75E-286 | 0.972210385 | 0.333 | 0.087 | 5.16E-282 | CD8_CXCL13 | TNFRSF9 |
| 222 | 4.50E-286 | 1.115199385 | 0.501 | 0.191 | 1.33E-281 | CD8_CXCL13 | FABP5 |
| 223 | 1.50E-274 | 1.154755399 | 0.559 | 0.239 | 4.44E-270 | CD8_CXCL13 | HLA-DRA |
| 224 | 9.67E-268 | 1.064346758 | 0.706 | 0.363 | 2.85E-263 | CD8_CXCL13 | HSPA1A |
| 225 | 4.85E-266 | 1.072892739 | 0.492 | 0.2 | 1.43E-261 | CD8_CXCL13 | LYST |
| 226 | 1.37E-264 | 1.007386778 | 0.502 | 0.204 | 4.03E-260 | CD8_CXCL13 | PTMS |
| 227 | 1.17E-256 | 1.028739818 | 0.464 | 0.184 | 3.45E-252 | CD8_CXCL13 | ACP5 |
| 228 | 7.94E-250 | 0.973970049 | 0.527 | 0.222 | 2.34E-245 | CD8_CXCL13 | METRNL |
| 229 | 1.26E-240 | 0.876309189 | 0.848 | 0.58 | 3.71E-236 | CD8_CXCL13 | CST7 |
| 230 | 5.10E-234 | 0.838834266 | 0.936 | 0.789 | 1.50E-229 | CD8_CXCL13 | CD74 |
| 231 | 1.40E-232 | 0.926965941 | 0.447 | 0.171 | 4.13E-228 | CD8_CXCL13 | HLA-DRB5 |
| 232 | 4.46E-213 | 0.989087172 | 0.605 | 0.311 | 1.32E-208 | CD8_CXCL13 | HSPH1 |
| 233 | 5.87E-211 | 0.821854384 | 0.725 | 0.445 | 1.73E-206 | CD8_CXCL13 | CD63 |
| 234 | 3.78E-202 | 0.888387083 | 0.27 | 0.081 | 1.11E-197 | CD8_CXCL13 | SNAP47 |
| 235 | 3.29E-192 | 0.787716616 | 0.336 | 0.117 | 9.71E-188 | CD8_CXCL13 | OASL |
| 236 | 6.72E-190 | 0.766563053 | 0.249 | 0.07 | 1.98E-185 | CD8_CXCL13 | KLRC3 |
| 237 | 1.47E-188 | 0.825943349 | 0.811 | 0.626 | 4.35E-184 | CD8_CXCL13 | RARRES3 |
| 238 | 1.93E-188 | 0.805566426 | 0.438 | 0.184 | 5.70E-184 | CD8_CXCL13 | CLIC3 |
| 239 | 2.95E-185 | 0.800593489 | 0.252 | 0.073 | 8.71E-181 | CD8_CXCL13 | PLPP1 |
| 240 | 2.79E-174 | 0.993436304 | 0.348 | 0.14 | 8.22E-170 | CD8_CXCL13 | FAM3C |
| 241 | 3.41E-173 | 0.855503061 | 0.397 | 0.164 | 1.00E-168 | CD8_CXCL13 | HLA-DQA1 |
| 242 | 8.07E-159 | 0.789108534 | 0.564 | 0.306 | 2.38E-154 | CD8_CXCL13 | CXCR6 |
| 243 | 7.14E-148 | 0.77320762 | 0.746 | 0.528 | 2.11E-143 | CD8_CXCL13 | CLEC2B |
| 244 | 4.50E-146 | 0.793256126 | 0.535 | 0.298 | 1.33E-141 | CD8_CXCL13 | DUSP5 |
| 245 | 4.48E-142 | 0.899074896 | 0.284 | 0.105 | 1.32E-137 | CD8_CXCL13 | CRTAM |
| 246 | 6.41E-138 | 0.878600605 | 0.434 | 0.228 | 1.89E-133 | CD8_CXCL13 | SRRT |
| 247 | 1.75E-132 | 0.797213373 | 0.519 | 0.292 | 5.15E-128 | CD8_CXCL13 | TIGIT |
| 248 | 3.73E-116 | 0.857031038 | 0.632 | 0.436 | 1.10E-111 | CD8_CXCL13 | HSPD1 |
| 249 | 1.57E-70 | 0.813580114 | 0.485 | 0.342 | 4.62E-66 | CD8_CXCL13 | CTSW |
| 250 | 1.85E-52 | 1.168260387 | 0.108 | 0.039 | 5.46E-48 | CD8_CXCL13 | TRBV28 |
| 251 | 0 | 3.117503707 | 0.972 | 0.165 | 0 | CD8_GZMK | GZMK |
| 252 | 0 | 1.417754166 | 0.934 | 0.59 | 0 | CD8_GZMK | CST7 |
| 253 | 5.66E-294 | 1.437466976 | 0.92 | 0.638 | 1.67E-289 | CD8_GZMK | DUSP2 |

| | | | | | | | |
|---|---|---|---|---|---|---|---|
| 254 | 1.90E-290 | 1.735560887 | 0.9 | 0.511 | 5.61E-286 | CD8_GZMK | CCL4 |
| 255 | 1.30E-184 | 1.402831513 | 0.399 | 0.13 | 3.84E-180 | CD8_GZMK | TNFSF9 |
| 256 | 3.84E-170 | 1.309506827 | 0.452 | 0.18 | 1.13E-165 | CD8_GZMK | CMC1 |
| 257 | 4.63E-170 | 1.03605001 | 0.933 | 0.798 | 1.37E-165 | CD8_GZMK | CD74 |
| 258 | 1.88E-167 | 1.520227688 | 0.601 | 0.284 | 5.56E-163 | CD8_GZMK | CCL4L2 |
| 259 | 4.33E-167 | 0.970995146 | 0.281 | 0.072 | 1.28E-162 | CD8_GZMK | KLRG1 |
| 260 | 2.15E-166 | 1.10228519 | 0.357 | 0.111 | 6.35E-162 | CD8_GZMK | CRTAM |
| 261 | 2.12E-154 | 0.773863037 | 0.865 | 0.544 | 6.26E-150 | CD8_GZMK | NKG7 |
| 262 | 1.15E-137 | 0.935334508 | 0.669 | 0.395 | 3.40E-133 | CD8_GZMK | HLA-DPB1 |
| 263 | 7.44E-135 | 1.018530479 | 0.474 | 0.223 | 2.19E-130 | CD8_GZMK | SH2D1A |
| 264 | 2.98E-114 | 0.885469426 | 0.414 | 0.177 | 8.79E-110 | CD8_GZMK | HLA-DQA1 |
| 265 | 8.33E-114 | 0.950953058 | 0.625 | 0.405 | 2.46E-109 | CD8_GZMK | GZMM |
| 266 | 4.36E-110 | 0.712784683 | 0.212 | 0.058 | 1.29E-105 | CD8_GZMK | EOMES |
| 267 | 2.86E-107 | 0.863168986 | 0.625 | 0.407 | 8.44E-103 | CD8_GZMK | ITGB2 |
| 268 | 7.32E-106 | 1.106656123 | 0.579 | 0.377 | 2.16E-101 | CD8_GZMK | CLDND1 |
| 269 | 7.47E-99 | 0.811721414 | 0.657 | 0.398 | 2.20E-94 | CD8_GZMK | HLA-DRB1 |
| 270 | 4.51E-97 | 0.967554906 | 0.701 | 0.536 | 1.33E-92 | CD8_GZMK | TAGAP |
| 271 | 1.02E-91 | 0.830353247 | 0.309 | 0.127 | 3.02E-87 | CD8_GZMK | SAMD3 |
| 272 | 2.66E-86 | 0.72198783 | 0.911 | 0.789 | 7.85E-82 | CD8_GZMK | CXCR4 |
| 273 | 1.82E-85 | 0.756080289 | 0.643 | 0.435 | 5.37E-81 | CD8_GZMK | HLA-DPA1 |
| 274 | 8.26E-76 | 0.677966821 | 0.226 | 0.083 | 2.44E-71 | CD8_GZMK | DTHD1 |
| 275 | 2.14E-75 | 0.967434235 | 0.375 | 0.193 | 6.32E-71 | CD8_GZMK | HLA-DRB5 |
| 276 | 2.21E-72 | 0.794675262 | 0.523 | 0.311 | 6.51E-68 | CD8_GZMK | GZMH |
| 277 | 6.31E-65 | 0.919358166 | 0.291 | 0.134 | 1.86E-60 | CD8_GZMK | TNF |
| 278 | 2.27E-63 | 0.832750556 | 0.467 | 0.269 | 6.70E-59 | CD8_GZMK | KLF2 |
| 279 | 2.78E-62 | 0.742295938 | 0.348 | 0.181 | 8.20E-58 | CD8_GZMK | HLA-DQB1 |
| 280 | 8.13E-61 | 0.697355692 | 0.643 | 0.507 | 2.40E-56 | CD8_GZMK | CD81 |
| 281 | 1.31E-60 | 0.76217303 | 0.394 | 0.231 | 3.85E-56 | CD8_GZMK | TRAT1 |
| 282 | 4.94E-59 | 0.58277118 | 0.188 | 0.071 | 1.46E-54 | CD8_GZMK | PLEK |
| 283 | 1.35E-58 | 0.540796054 | 0.736 | 0.614 | 3.97E-54 | CD8_GZMK | CD44 |
| 284 | 2.10E-57 | 0.696898203 | 0.229 | 0.096 | 6.18E-53 | CD8_GZMK | CCL3L1 |
| 285 | 5.11E-49 | 0.841890657 | 0.394 | 0.253 | 1.51E-44 | CD8_GZMK | ITM2C |
| 286 | 2.34E-45 | 0.662272912 | 0.317 | 0.182 | 6.90E-41 | CD8_GZMK | LYAR |
| 287 | 1.53E-44 | 0.692874154 | 0.531 | 0.401 | 4.51E-40 | CD8_GZMK | PIK3R1 |
| 288 | 1.82E-43 | 0.548542371 | 0.633 | 0.51 | 5.38E-39 | CD8_GZMK | IER2 |
| 289 | 9.30E-41 | 0.846062931 | 0.239 | 0.12 | 2.74E-36 | CD8_GZMK | IFNG |
| 290 | 1.58E-40 | 0.582415322 | 0.238 | 0.124 | 4.67E-36 | CD8_GZMK | F2R |
| 291 | 3.10E-38 | 0.620486542 | 0.254 | 0.138 | 9.16E-34 | CD8_GZMK | LINC00861 |
| 292 | 9.68E-37 | 0.545004148 | 0.462 | 0.324 | 2.86E-32 | CD8_GZMK | CD27 |
| 293 | 5.34E-36 | 0.593498011 | 0.558 | 0.454 | 1.57E-31 | CD8_GZMK | TUBA4A |
| 294 | 1.80E-33 | 0.608606029 | 0.399 | 0.27 | 5.31E-29 | CD8_GZMK | HLA-DRA |
| 295 | 5.76E-32 | 0.556648046 | 0.138 | 0.06 | 1.70E-27 | CD8_GZMK | IGLV7-46 |
| 296 | 3.80E-31 | 0.558842795 | 0.424 | 0.313 | 1.12E-26 | CD8_GZMK | APOBEC3G |
| 297 | 8.46E-28 | 0.603110601 | 0.34 | 0.242 | 2.50E-23 | CD8_GZMK | NFATC2 |

| | | | | | | | |
|---|---|---|---|---|---|---|---|
| 298 | 1.60E-22 | 0.9254489 | 0.178 | 0.101 | 4.73E-18 | CD8_GZMK | IGLV1-44 |
| 299 | 2.09E-13 | 0.589472086 | 0.133 | 0.08 | 6.15E-09 | CD8_GZMK | IGLV2-11 |
| 300 | 2.99E-09 | 1.088447882 | 0.131 | 0.087 | 8.81E-05 | CD8_GZMK | IGLV6-57 |
| 301 | 0 | 3.167712221 | 0.488 | 0.072 | 0 | CD8_IFNG | HSPA6 |
| 302 | 0 | 2.891250535 | 0.953 | 0.273 | 0 | CD8_IFNG | HSPA1B |
| 303 | 0 | 2.606298533 | 0.978 | 0.636 | 0 | CD8_IFNG | DNAJB1 |
| 304 | 0 | 2.440181697 | 0.967 | 0.37 | 0 | CD8_IFNG | HSPA1A |
| 305 | 0 | 2.34529392 | 0.219 | 0.021 | 0 | CD8_IFNG | PGA3 |
| 306 | 0 | 1.993636953 | 0.983 | 0.877 | 0 | CD8_IFNG | HSP90AA1 |
| 307 | 5.08E-290 | 1.729056181 | 0.969 | 0.882 | 1.50E-285 | CD8_IFNG | HSPA8 |
| 308 | 9.00E-288 | 2.08355341 | 0.748 | 0.322 | 2.66E-283 | CD8_IFNG | HSPH1 |
| 309 | 3.10E-265 | 1.859882224 | 0.86 | 0.56 | 9.15E-261 | CD8_IFNG | HSPE1 |
| 310 | 3.76E-250 | 1.68607567 | 0.713 | 0.28 | 1.11E-245 | CD8_IFNG | CCL4L2 |
| 311 | 7.43E-237 | 1.471188328 | 0.595 | 0.197 | 2.19E-232 | CD8_IFNG | GZMK |
| 312 | 1.06E-207 | 1.558201453 | 0.889 | 0.628 | 3.12E-203 | CD8_IFNG | JUN |
| 313 | 5.90E-190 | 1.287792513 | 0.348 | 0.09 | 1.74E-185 | CD8_IFNG | DNAJB4 |
| 314 | 1.46E-185 | 1.914966681 | 0.498 | 0.191 | 4.29E-181 | CD8_IFNG | NR4A1 |
| 315 | 5.85E-167 | 1.336834831 | 0.871 | 0.644 | 1.72E-162 | CD8_IFNG | DUSP2 |
| 316 | 7.06E-167 | 1.743702056 | 0.427 | 0.147 | 2.08E-162 | CD8_IFNG | BAG3 |
| 317 | 5.23E-166 | 1.924259552 | 0.373 | 0.113 | 1.54E-161 | CD8_IFNG | IFNG |
| 318 | 4.52E-161 | 1.318270231 | 0.297 | 0.076 | 1.33E-156 | CD8_IFNG | DNAJA4 |
| 319 | 2.66E-151 | 1.507951524 | 0.767 | 0.561 | 7.84E-147 | CD8_IFNG | DNAJA1 |
| 320 | 3.45E-151 | 1.635228241 | 0.708 | 0.445 | 1.02E-146 | CD8_IFNG | HSPD1 |
| 321 | 1.54E-149 | 1.503705031 | 0.678 | 0.409 | 4.53E-145 | CD8_IFNG | HSPB1 |
| 322 | 2.82E-143 | 1.794169923 | 0.369 | 0.127 | 8.33E-139 | CD8_IFNG | ZFAND2A |
| 323 | 5.41E-138 | 0.920466821 | 0.952 | 0.829 | 1.60E-133 | CD8_IFNG | JUND |
| 324 | 3.03E-130 | 1.250662479 | 0.68 | 0.424 | 8.94E-126 | CD8_IFNG | CACYBP |
| 325 | 8.99E-127 | 1.733127357 | 0.372 | 0.134 | 2.65E-122 | CD8_IFNG | TNFSF9 |
| 326 | 4.31E-124 | 1.576673902 | 0.253 | 0.068 | 1.27E-119 | CD8_IFNG | SERPINH1 |
| 327 | 8.24E-118 | 1.304729213 | 0.875 | 0.815 | 2.43E-113 | CD8_IFNG | HSP90AB1 |
| 328 | 1.02E-115 | 1.284289988 | 0.342 | 0.123 | 3.02E-111 | CD8_IFNG | RHOB |
| 329 | 1.71E-107 | 1.446568253 | 0.413 | 0.186 | 5.05E-103 | CD8_IFNG | NEU1 |
| 330 | 1.96E-107 | 1.418534146 | 0.409 | 0.18 | 5.78E-103 | CD8_IFNG | FKBP4 |
| 331 | 1.00E-104 | 1.160762079 | 0.883 | 0.788 | 2.96E-100 | CD8_IFNG | DUSP1 |
| 332 | 6.44E-86 | 1.055389754 | 0.271 | 0.095 | 1.90E-81 | CD8_IFNG | CCL3L1 |
| 333 | 2.38E-80 | 0.997321068 | 0.573 | 0.358 | 7.01E-76 | CD8_IFNG | TSPYL2 |
| 334 | 5.28E-80 | 0.850741944 | 0.816 | 0.677 | 1.56E-75 | CD8_IFNG | TXNIP |
| 335 | 6.94E-80 | 0.990545384 | 0.957 | 0.956 | 2.05E-75 | CD8_IFNG | UBC |
| 336 | 2.13E-60 | 0.865347095 | 0.218 | 0.084 | 6.29E-56 | CD8_IFNG | IER5L |
| 337 | 1.53E-58 | 0.976095085 | 0.25 | 0.108 | 4.52E-54 | CD8_IFNG | TNFSF14 |
| 338 | 6.05E-57 | 0.895999289 | 0.398 | 0.231 | 1.78E-52 | CD8_IFNG | CHORDC1 |
| 339 | 2.10E-55 | 0.96299858 | 0.31 | 0.151 | 6.20E-51 | CD8_IFNG | ATF3 |
| 340 | 6.62E-51 | 0.849284907 | 0.872 | 0.862 | 1.95E-46 | CD8_IFNG | UBB |
| 341 | 9.23E-49 | 0.966773572 | 0.157 | 0.055 | 2.72E-44 | CD8_IFNG | LIPF |

| | | | | | | | |
|---|---|---|---|---|---|---|---|
| 342 | 3.54E-46 | 0.8757715 | 0.508 | 0.365 | 1.04E-41 | CD8_IFNG | DOK2 |
| 343 | 1.67E-43 | 0.873355172 | 0.189 | 0.079 | 4.93E-39 | CD8_IFNG | EGR1 |
| 344 | 3.43E-40 | 0.998887244 | 0.432 | 0.273 | 1.01E-35 | CD8_IFNG | KLF2 |
| 345 | 1.57E-33 | 1.245552204 | 0.253 | 0.138 | 4.62E-29 | CD8_IFNG | TNF |
| 346 | 2.71E-33 | 0.955199686 | 0.26 | 0.14 | 8.01E-29 | CD8_IFNG | LINC02446 |
| 347 | 7.52E-33 | 0.965688014 | 0.363 | 0.242 | 2.22E-28 | CD8_IFNG | GADD45B |
| 348 | 1.36E-22 | 1.142874899 | 0.216 | 0.131 | 4.00E-18 | CD8_IFNG | PGC |
| 349 | 1.24E-06 | 0.88965862 | 0.16 | 0.117 | 0.036540391 | CD8_IFNG | IGKV3-11 |
| 350 | 0.004453665 | 0.99922895 | 0.092 | 0.121 | 1 | CD8_IFNG | IGLV3-1 |
| 351 | 0 | 1.705005479 | 0.416 | 0.108 | 0 | CD8_GZMB | XCL1 |
| 352 | 0 | 1.686520827 | 0.879 | 0.452 | 0 | CD8_GZMB | CCL4 |
| 353 | 0 | 1.543760307 | 0.479 | 0.11 | 0 | CD8_GZMB | KLRC1 |
| 354 | 0 | 1.533265768 | 0.523 | 0.15 | 0 | CD8_GZMB | XCL2 |
| 355 | 0 | 1.459336867 | 0.82 | 0.305 | 0 | CD8_GZMB | CD8A |
| 356 | 0 | 1.405536643 | 0.756 | 0.253 | 0 | CD8_GZMB | CD8B |
| 357 | 0 | 0.957382063 | 0.439 | 0.166 | 0 | CD8_GZMB | ITGA1 |
| 358 | 0 | 0.888283381 | 0.952 | 0.865 | 0 | CD8_GZMB | ZFP36L2 |
| 359 | 0 | 0.862967884 | 0.99 | 0.598 | 0 | CD8_GZMB | CCL5 |
| 360 | 8.21E-296 | 0.726038903 | 0.592 | 0.277 | 2.42E-291 | CD8_GZMB | HOPX |
| 361 | 4.84E-273 | 1.124987564 | 0.131 | 0.013 | 1.43E-268 | CD8_GZMB | TRBV4-2 |
| 362 | 1.76E-254 | 0.488327535 | 0.996 | 0.981 | 5.18E-250 | CD8_GZMB | MT-CO3 |
| 363 | 9.55E-236 | 0.827533619 | 0.24 | 0.067 | 2.82E-231 | CD8_GZMB | ZNF683 |
| 364 | 5.35E-212 | 0.513154381 | 0.936 | 0.825 | 1.58E-207 | CD8_GZMB | S100A6 |
| 365 | 6.42E-211 | 0.729408057 | 0.548 | 0.305 | 1.89E-206 | CD8_GZMB | CAPG |
| 366 | 5.10E-208 | 0.832242102 | 0.361 | 0.157 | 1.50E-203 | CD8_GZMB | AUTS2 |
| 367 | 1.22E-195 | 0.825882086 | 0.546 | 0.335 | 3.60E-191 | CD8_GZMB | CD55 |
| 368 | 2.24E-191 | 0.781947384 | 0.601 | 0.39 | 6.61E-187 | CD8_GZMB | PTGER4 |
| 369 | 3.27E-191 | 0.815918745 | 0.324 | 0.136 | 9.64E-187 | CD8_GZMB | SPRY1 |
| 370 | 1.90E-177 | 2.060084437 | 0.27 | 0.109 | 5.60E-173 | CD8_GZMB | MTCO1P12 |
| 371 | 1.27E-164 | 0.702290987 | 0.862 | 0.751 | 3.74E-160 | CD8_GZMB | CD52 |
| 372 | 1.39E-164 | 0.536864899 | 0.902 | 0.772 | 4.11E-160 | CD8_GZMB | CXCR4 |
| 373 | 1.42E-149 | 0.611423907 | 0.742 | 0.549 | 4.18E-145 | CD8_GZMB | IL7R |
| 374 | 3.17E-146 | 0.590150338 | 0.692 | 0.524 | 9.35E-142 | CD8_GZMB | CKLF |
| 375 | 1.45E-138 | 0.837541381 | 0.406 | 0.233 | 4.26E-134 | CD8_GZMB | TOB1 |
| 376 | 7.75E-135 | 0.523189427 | 0.754 | 0.596 | 2.29E-130 | CD8_GZMB | CD3G |
| 377 | 2.13E-119 | 0.648798705 | 0.593 | 0.417 | 6.30E-115 | CD8_GZMB | ID2 |
| 378 | 1.77E-118 | 0.612629427 | 0.544 | 0.381 | 5.21E-114 | CD8_GZMB | PARP8 |
| 379 | 1.12E-109 | 0.496965335 | 0.62 | 0.445 | 3.31E-105 | CD8_GZMB | CD63 |
| 380 | 7.83E-107 | 0.53110228 | 0.23 | 0.104 | 2.31E-102 | CD8_GZMB | SMIM3 |
| 381 | 1.66E-106 | 0.854910033 | 0.227 | 0.102 | 4.91E-102 | CD8_GZMB | AC092580.4 |
| 382 | 7.32E-105 | 0.584672418 | 0.215 | 0.095 | 2.16E-100 | CD8_GZMB | MYBL1 |
| 383 | 8.66E-100 | 0.502440147 | 0.84 | 0.723 | 2.55E-95 | CD8_GZMB | CD69 |
| 384 | 3.84E-96 | 0.512494132 | 0.443 | 0.289 | 1.13E-91 | CD8_GZMB | FKBP11 |
| 385 | 6.60E-96 | 1.549735664 | 0.423 | 0.277 | 1.95E-91 | CD8_GZMB | CCL4L2 |

| | | | | | | | |
|---|---|---|---|---|---|---|---|
| 386 | 3.50E-94 | 0.507209384 | 0.486 | 0.335 | 1.03E-89 | CD8_GZMB | ITGAE |
| 387 | 2.07E-91 | 0.582090613 | 0.873 | 0.83 | 6.12E-87 | CD8_GZMB | ZFP36 |
| 388 | 2.40E-91 | 0.532663637 | 0.458 | 0.312 | 7.08E-87 | CD8_GZMB | RUNX3 |
| 389 | 2.76E-91 | 0.492554131 | 0.344 | 0.203 | 8.15E-87 | CD8_GZMB | THEMIS |
| 390 | 1.10E-90 | 0.541958982 | 0.378 | 0.237 | 3.26E-86 | CD8_GZMB | SCML4 |
| 391 | 1.25E-82 | 0.508392641 | 0.256 | 0.135 | 3.68E-78 | CD8_GZMB | PTGER2 |
| 392 | 2.29E-82 | 0.515004621 | 0.211 | 0.102 | 6.77E-78 | CD8_GZMB | TFF2 |
| 393 | 1.90E-79 | 0.52311923 | 0.293 | 0.172 | 5.60E-75 | CD8_GZMB | ANKRD28 |
| 394 | 1.09E-68 | 0.702162028 | 0.488 | 0.373 | 3.22E-64 | CD8_GZMB | FOSB |
| 395 | 1.40E-53 | 0.657885834 | 0.667 | 0.585 | 4.12E-49 | CD8_GZMB | FOS |
| 396 | 3.84E-53 | 0.598358682 | 0.305 | 0.206 | 1.13E-48 | CD8_GZMB | AC020916.1 |
| 397 | 1.22E-49 | 0.510772968 | 0.34 | 0.243 | 3.60E-45 | CD8_GZMB | PFKFB3 |
| 398 | 5.30E-37 | 0.51518577 | 0.16 | 0.092 | 1.56E-32 | CD8_GZMB | CCL3L1 |
| 399 | 7.93E-16 | 0.512949572 | 0.289 | 0.246 | 2.34E-11 | CD8_GZMB | ATP5E |
| 400 | 1.15E-11 | 0.552499549 | 0.118 | 0.085 | 3.39E-07 | CD8_GZMB | IGKV3-15 |
| 401 | 0 | 1.480871419 | 0.834 | 0.404 | 0 | CD4_IL7R | KLRB1 |
| 402 | 0 | 1.455039715 | 0.894 | 0.543 | 0 | CD4_IL7R | IL7R |
| 403 | 0 | 1.243561564 | 0.446 | 0.098 | 0 | CD4_IL7R | CD40LG |
| 404 | 0 | 1.091571243 | 0.746 | 0.401 | 0 | CD4_IL7R | RORA |
| 405 | 0 | 0.978326079 | 0.36 | 0.09 | 0 | CD4_IL7R | CCR6 |
| 406 | 0 | 0.825855748 | 0.979 | 0.798 | 0 | CD4_IL7R | S100A4 |
| 407 | 0 | 0.560817737 | 1 | 1 | 0 | CD4_IL7R | RPLP1 |
| 408 | 1.58E-307 | 0.675878131 | 0.179 | 0.022 | 4.66E-303 | CD4_IL7R | IL4I1 |
| 409 | 1.40E-248 | 1.109945234 | 0.295 | 0.077 | 4.13E-244 | CD4_IL7R | CCL20 |
| 410 | 4.32E-234 | 0.514818469 | 0.998 | 0.989 | 1.28E-229 | CD4_IL7R | TPT1 |
| 411 | 4.40E-198 | 0.634661805 | 0.2 | 0.044 | 1.30E-193 | CD4_IL7R | TNFSF13B |
| 412 | 2.01E-187 | 0.523443706 | 0.172 | 0.035 | 5.94E-183 | CD4_IL7R | DPP4 |
| 413 | 3.56E-182 | 0.847994769 | 0.701 | 0.405 | 1.05E-177 | CD4_IL7R | LTB |
| 414 | 1.49E-176 | 0.722050393 | 0.389 | 0.161 | 4.41E-172 | CD4_IL7R | AQP3 |
| 415 | 1.93E-165 | 0.568739137 | 0.951 | 0.832 | 5.70E-161 | CD4_IL7R | S100A6 |
| 416 | 3.80E-150 | 0.773999046 | 0.404 | 0.187 | 1.12E-145 | CD4_IL7R | IFNGR1 |
| 417 | 3.63E-144 | 0.783334223 | 0.611 | 0.362 | 1.07E-139 | CD4_IL7R | GPR183 |
| 418 | 3.74E-137 | 0.81780641 | 0.852 | 0.733 | 1.10E-132 | CD4_IL7R | TNFAIP3 |
| 419 | 3.66E-135 | 0.61684906 | 0.527 | 0.289 | 1.08E-130 | CD4_IL7R | FKBP11 |
| 420 | 1.07E-128 | 0.59602474 | 0.31 | 0.129 | 3.16E-124 | CD4_IL7R | TNFRSF25 |
| 421 | 2.09E-128 | 0.530407594 | 0.237 | 0.083 | 6.16E-124 | CD4_IL7R | RUNX2 |
| 422 | 8.97E-125 | 0.552734636 | 0.245 | 0.09 | 2.65E-120 | CD4_IL7R | IFI44 |
| 423 | 6.83E-121 | 0.610203304 | 0.281 | 0.115 | 2.02E-116 | CD4_IL7R | PRR5 |
| 424 | 2.44E-117 | 0.68478426 | 0.457 | 0.248 | 7.20E-113 | CD4_IL7R | ERN1 |
| 425 | 1.59E-116 | 0.646981734 | 0.332 | 0.149 | 4.69E-112 | CD4_IL7R | TMIGD2 |
| 426 | 1.11E-111 | 0.593807279 | 0.903 | 0.782 | 3.28E-107 | CD4_IL7R | CXCR4 |
| 427 | 2.38E-110 | 0.531849561 | 0.3 | 0.131 | 7.03E-106 | CD4_IL7R | PERP |
| 428 | 9.69E-108 | 0.602349171 | 0.516 | 0.31 | 2.86E-103 | CD4_IL7R | MGAT4A |
| 429 | 1.99E-102 | 0.501589882 | 0.799 | 0.579 | 5.88E-98 | CD4_IL7R | ANXA1 |

| | | | | | | | |
|---|---|---|---|---|---|---|---|
| 430 | 1.99E-101 | 0.535634322 | 0.233 | 0.093 | 5.86E-97 | CD4_IL7R | ADAM19 |
| 431 | 3.01E-100 | 0.555840943 | 0.35 | 0.174 | 8.87E-96 | CD4_IL7R | ANKRD28 |
| 432 | 8.52E-97 | 0.542825173 | 0.893 | 0.756 | 2.51E-92 | CD4_IL7R | CD52 |
| 433 | 5.07E-84 | 0.500059931 | 0.235 | 0.102 | 1.49E-79 | CD4_IL7R | MYBL1 |
| 434 | 3.17E-83 | 0.502838974 | 0.624 | 0.447 | 9.34E-79 | CD4_IL7R | YWHAQ |
| 435 | 5.59E-79 | 0.566648755 | 0.94 | 0.853 | 1.65E-74 | CD4_IL7R | VIM |
| 436 | 6.56E-79 | 0.527313485 | 0.429 | 0.255 | 1.93E-74 | CD4_IL7R | ODF2L |
| 437 | 1.88E-78 | 0.571639111 | 0.174 | 0.067 | 5.55E-74 | CD4_IL7R | AMICA1 |
| 438 | 6.22E-77 | 0.56041462 | 0.286 | 0.144 | 1.83E-72 | CD4_IL7R | SESN1 |
| 439 | 3.69E-70 | 0.622248559 | 0.851 | 0.732 | 1.09E-65 | CD4_IL7R | CD69 |
| 440 | 1.18E-69 | 0.54314146 | 0.487 | 0.316 | 3.49E-65 | CD4_IL7R | CXCR6 |
| 441 | 2.04E-68 | 0.65982584 | 0.721 | 0.584 | 6.02E-64 | CD4_IL7R | FOS |
| 442 | 6.60E-67 | 0.640306708 | 0.233 | 0.111 | 1.95E-62 | CD4_IL7R | AC092580.4 |
| 443 | 2.71E-66 | 0.722130208 | 0.364 | 0.217 | 8.00E-62 | CD4_IL7R | CEBPD |
| 444 | 4.19E-62 | 0.667711979 | 0.793 | 0.685 | 1.23E-57 | CD4_IL7R | NFKBIA |
| 445 | 4.72E-57 | 0.695020167 | 0.725 | 0.605 | 1.39E-52 | CD4_IL7R | RGS1 |
| 446 | 1.93E-54 | 0.490661088 | 0.585 | 0.433 | 5.69E-50 | CD4_IL7R | ID2 |
| 447 | 3.26E-37 | 0.511694631 | 0.145 | 0.072 | 9.61E-33 | CD4_IL7R | RP11-138A9.2 |
| 448 | 7.32E-33 | 0.521650816 | 0.539 | 0.442 | 2.16E-28 | CD4_IL7R | AHNAK |
| 449 | 1.82E-23 | 0.794788369 | 0.102 | 0.053 | 5.37E-19 | CD4_IL7R | TRBV20-1 |
| 450 | 9.53E-13 | 0.505638624 | 0.263 | 0.208 | 2.81E-08 | CD4_IL7R | LMNA |
| 451 | 0 | 1.836204661 | 0.543 | 0.069 | 0 | CD4_CCR7 | CCR7 |
| 452 | 0 | 1.525231071 | 0.911 | 0.76 | 0 | CD4_CCR7 | RPL7 |
| 453 | 0 | 1.38664999 | 0.4 | 0.08 | 0 | CD4_CCR7 | SELL |
| 454 | 0 | 1.379420068 | 0.963 | 0.921 | 0 | CD4_CCR7 | RPL21 |
| 455 | 0 | 1.34104015 | 0.78 | 0.367 | 0 | CD4_CCR7 | LTB |
| 456 | 0 | 1.326992832 | 0.894 | 0.77 | 0 | CD4_CCR7 | RPL13A |
| 457 | 0 | 1.207498621 | 0.368 | 0.044 | 0 | CD4_CCR7 | LEF1 |
| 458 | 0 | 1.193929008 | 0.466 | 0.118 | 0 | CD4_CCR7 | TCF7 |
| 459 | 0 | 1.151071502 | 0.999 | 0.994 | 0 | CD4_CCR7 | RPL34 |
| 460 | 0 | 1.095937109 | 0.995 | 0.977 | 0 | CD4_CCR7 | RPS6 |
| 461 | 0 | 1.084653592 | 0.919 | 0.827 | 0 | CD4_CCR7 | RPL31 |
| 462 | 0 | 1.066012648 | 0.287 | 0.034 | 0 | CD4_CCR7 | TSHZ2 |
| 463 | 0 | 1.007037205 | 0.849 | 0.615 | 0 | CD4_CCR7 | LDHB |
| 464 | 0 | 1.006773828 | 0.707 | 0.45 | 0 | CD4_CCR7 | EIF3E |
| 465 | 0 | 1.000665189 | 0.993 | 0.97 | 0 | CD4_CCR7 | RPL3 |
| 466 | 0 | 0.985229635 | 0.975 | 0.926 | 0 | CD4_CCR7 | RPL9 |
| 467 | 0 | 0.95868876 | 0.995 | 0.979 | 0 | CD4_CCR7 | RPS2 |
| 468 | 0 | 0.950061454 | 0.998 | 0.983 | 0 | CD4_CCR7 | RPS3A |
| 469 | 0 | 0.923725875 | 0.999 | 0.992 | 0 | CD4_CCR7 | RPS18 |
| 470 | 0 | 0.91511854 | 0.998 | 0.993 | 0 | CD4_CCR7 | RPS8 |
| 471 | 0 | 0.908648174 | 0.991 | 0.985 | 0 | CD4_CCR7 | RPL41 |
| 472 | 0 | 0.869147291 | 1 | 0.997 | 0 | CD4_CCR7 | RPL32 |
| 473 | 0 | 0.836925893 | 0.928 | 0.798 | 0 | CD4_CCR7 | PABPC1 |

| | | | | | | | |
|---|---|---|---|---|---|---|---|
| 474 | 0 | 0.827228574 | 0.975 | 0.922 | 0 | CD4_CCR7 | RPL22 |
| 475 | 0 | 0.815798365 | 0.998 | 0.997 | 0 | CD4_CCR7 | RPS27 |
| 476 | 0 | 0.806252774 | 0.975 | 0.935 | 0 | CD4_CCR7 | RPL10A |
| 477 | 0 | 0.791975789 | 0.939 | 0.853 | 0 | CD4_CCR7 | RPL4 |
| 478 | 0 | 0.784678905 | 0.995 | 0.982 | 0 | CD4_CCR7 | RPL36 |
| 479 | 0 | 0.784062861 | 0.999 | 0.997 | 0 | CD4_CCR7 | RPL39 |
| 480 | 0 | 0.780251948 | 1 | 0.997 | 0 | CD4_CCR7 | EEF1A1 |
| 481 | 0 | 0.772415194 | 0.992 | 0.98 | 0 | CD4_CCR7 | RPLP2 |
| 482 | 0 | 0.771400165 | 1 | 0.998 | 0 | CD4_CCR7 | RPL13 |
| 483 | 1.61E-303 | 1.32166833 | 0.516 | 0.23 | 4.76E-299 | CD4_CCR7 | KLF2 |
| 484 | 2.45E-290 | 1.341844661 | 0.473 | 0.208 | 7.22E-286 | CD4_CCR7 | GNB2L1 |
| 485 | 1.48E-262 | 1.053363449 | 0.918 | 0.845 | 4.36E-258 | CD4_CCR7 | RPS29 |
| 486 | 4.69E-260 | 0.933165901 | 0.85 | 0.709 | 1.38E-255 | CD4_CCR7 | RPL36A |
| 487 | 1.24E-256 | 1.047519886 | 0.436 | 0.18 | 3.65E-252 | CD4_CCR7 | TMEM66 |
| 488 | 8.83E-249 | 1.022551257 | 0.763 | 0.594 | 2.60E-244 | CD4_CCR7 | RPL23 |
| 489 | 3.16E-245 | 1.001109845 | 0.385 | 0.15 | 9.33E-241 | CD4_CCR7 | GLTSCR2 |
| 490 | 7.36E-237 | 0.876341741 | 0.614 | 0.343 | 2.17E-232 | CD4_CCR7 | GPR183 |
| 491 | 1.22E-236 | 0.80046029 | 0.884 | 0.779 | 3.59E-232 | CD4_CCR7 | RPS10 |
| 492 | 1.06E-235 | 0.845547876 | 0.847 | 0.719 | 3.12E-231 | CD4_CCR7 | RPL38 |
| 493 | 1.15E-232 | 0.806322148 | 0.884 | 0.78 | 3.40E-228 | CD4_CCR7 | RPS11 |
| 494 | 4.49E-223 | 1.159714829 | 0.788 | 0.649 | 1.33E-218 | CD4_CCR7 | RPL27A |
| 495 | 1.17E-222 | 0.793346622 | 0.48 | 0.24 | 3.44E-218 | CD4_CCR7 | TMEM123 |
| 496 | 4.81E-204 | 0.906309049 | 0.337 | 0.132 | 1.42E-199 | CD4_CCR7 | FYB |
| 497 | 2.83E-193 | 0.776133905 | 0.381 | 0.163 | 8.34E-189 | CD4_CCR7 | ATP5G2 |
| 498 | 1.22E-190 | 1.128102976 | 0.693 | 0.534 | 3.60E-186 | CD4_CCR7 | RPS20 |
| 499 | 1.61E-179 | 0.877782623 | 0.896 | 0.817 | 4.74E-175 | CD4_CCR7 | JUNB |
| 500 | 7.14E-168 | 0.782020889 | 0.426 | 0.23 | 2.10E-163 | CD4_CCR7 | NOSIP |
| 501 | 0 | 2.980103202 | 0.65 | 0.096 | 0 | Treg | TNFRSF4 |
| 502 | 0 | 2.440463012 | 0.872 | 0.285 | 0 | Treg | BATF |
| 503 | 0 | 2.044985839 | 0.701 | 0.168 | 0 | Treg | TNFRSF18 |
| 504 | 0 | 2.043852872 | 0.512 | 0.02 | 0 | Treg | FOXP3 |
| 505 | 0 | 1.986470671 | 0.904 | 0.606 | 0 | Treg | SAT1 |
| 506 | 0 | 1.936327175 | 0.806 | 0.259 | 0 | Treg | CTLA4 |
| 507 | 0 | 1.610937994 | 0.779 | 0.399 | 0 | Treg | CACYBP |
| 508 | 0 | 1.587840745 | 0.478 | 0.068 | 0 | Treg | GK |
| 509 | 0 | 1.54927262 | 0.358 | 0.049 | 0 | Treg | IL2RA |
| 510 | 0 | 1.543193947 | 0.74 | 0.293 | 0 | Treg | DUSP4 |
| 511 | 0 | 1.535204625 | 0.747 | 0.358 | 0 | Treg | CARD16 |
| 512 | 0 | 1.499388837 | 0.61 | 0.221 | 0 | Treg | GPX1P1 |
| 513 | 0 | 1.465692494 | 0.652 | 0.222 | 0 | Treg | DNPH1 |
| 514 | 0 | 1.363190406 | 0.446 | 0.074 | 0 | Treg | TBC1D4 |
| 515 | 0 | 1.315522264 | 0.863 | 0.504 | 0 | Treg | SPOCK2 |
| 516 | 0 | 1.297138402 | 0.485 | 0.162 | 0 | Treg | MIR4435-2HG |
| 517 | 0 | 1.296344723 | 0.244 | 0.033 | 0 | Treg | SOX4 |

| | | | | | | | |
|---|---|---|---|---|---|---|---|
| 518 | 0 | 1.267804092 | 0.289 | 0.031 | 0 | Treg | LAIR2 |
| 519 | 0 | 1.227921135 | 0.265 | 0.008 | 0 | Treg | IL1R2 |
| 520 | 0 | 1.062822663 | 0.225 | 0.003 | 0 | Treg | CCR8 |
| 521 | 0 | 1.061801414 | 0.328 | 0.064 | 0 | Treg | GNA15 |
| 522 | 0 | 1.060151463 | 0.103 | 0.001 | 0 | Treg | CD177 |
| 523 | 3.91E-288 | 1.129637679 | 0.615 | 0.263 | 1.15E-283 | Treg | ICOS |
| 524 | 3.07E-278 | 1.061696345 | 0.457 | 0.155 | 9.05E-274 | Treg | ZNRF1 |
| 525 | 2.62E-276 | 1.056336068 | 0.352 | 0.091 | 7.73E-272 | Treg | TNFRSF9 |
| 526 | 2.75E-275 | 1.173026968 | 0.634 | 0.284 | 8.12E-271 | Treg | TIGIT |
| 527 | 9.53E-265 | 1.092142718 | 0.665 | 0.33 | 2.81E-260 | Treg | PBXIP1 |
| 528 | 5.57E-260 | 1.131273718 | 0.491 | 0.182 | 1.64E-255 | Treg | PELI1 |
| 529 | 3.50E-255 | 1.775501318 | 0.697 | 0.372 | 1.03E-250 | Treg | HSPA1A |
| 530 | 1.16E-239 | 1.11855554 | 0.626 | 0.29 | 3.43E-235 | Treg | PHLDA1 |
| 531 | 4.99E-239 | 1.205712865 | 0.293 | 0.075 | 1.47E-234 | Treg | MAGEH1 |
| 532 | 3.71E-222 | 1.058760977 | 0.579 | 0.277 | 1.09E-217 | Treg | CORO1B |
| 533 | 7.10E-217 | 1.082454113 | 0.609 | 0.309 | 2.09E-212 | Treg | BTG3 |
| 534 | 9.18E-216 | 1.042059282 | 0.758 | 0.494 | 2.71E-211 | Treg | UCP2 |
| 535 | 2.44E-214 | 1.030163042 | 0.842 | 0.651 | 7.19E-210 | Treg | SOD1 |
| 536 | 9.73E-182 | 1.223539956 | 0.283 | 0.084 | 2.87E-177 | Treg | CCL20 |
| 537 | 9.84E-178 | 1.370560448 | 0.588 | 0.319 | 2.90E-173 | Treg | HSPH1 |
| 538 | 1.03E-160 | 1.557465672 | 0.542 | 0.286 | 3.04E-156 | Treg | HSPA1B |
| 539 | 6.23E-159 | 1.376928376 | 0.67 | 0.436 | 1.84E-154 | Treg | HSPD1 |
| 540 | 1.56E-153 | 1.231791663 | 0.637 | 0.4 | 4.60E-149 | Treg | HSPB1 |
| 541 | 3.51E-147 | 1.368384739 | 0.943 | 0.876 | 1.04E-142 | Treg | HSP90AA1 |
| 542 | 2.29E-143 | 1.120285506 | 0.22 | 0.063 | 6.77E-139 | Treg | SERPINH1 |
| 543 | 5.10E-143 | 1.41333807 | 0.747 | 0.559 | 1.50E-138 | Treg | HSPE1 |
| 544 | 9.35E-134 | 1.035603672 | 0.381 | 0.171 | 2.76E-129 | Treg | FKBP4 |
| 545 | 9.69E-132 | 1.119273767 | 0.891 | 0.81 | 2.86E-127 | Treg | HSP90AB1 |
| 546 | 3.77E-119 | 1.26922724 | 0.428 | 0.224 | 1.11E-114 | Treg | PLIN2 |
| 547 | 9.76E-110 | 1.333920472 | 0.322 | 0.145 | 2.88E-105 | Treg | BAG3 |
| 548 | 7.11E-76 | 1.331052662 | 0.748 | 0.645 | 2.10E-71 | Treg | DNAJB1 |
| 549 | 3.78E-18 | 1.112536617 | 0.1 | 0.055 | 1.12E-13 | Treg | TRBV20-1 |
| 550 | 1.45E-09 | 1.286874132 | 0.121 | 0.084 | 4.27E-05 | Treg | CXCL13 |

Supplementary Table S5 DEGs of myeloid cells subclusters

| | p_val | avg_log2FC | pct.1 | pct.2 | p_val_adj | cluster | gene |
|---|---|---|---|---|---|---|---|
| 1 | 0 | 7.617678746 | 0.999 | 0.014 | 0 | CPA3_Mast | TPSAB1 |
| 2 | 0 | 6.346406347 | 0.762 | 0.006 | 0 | CPA3_Mast | TPSB2 |
| 3 | 0 | 5.978952223 | 0.965 | 0.004 | 0 | CPA3_Mast | CPA3 |
| 4 | 0 | 5.074217987 | 0.645 | 0.001 | 0 | CPA3_Mast | TPSD1 |
| 5 | 0 | 4.262540984 | 0.9 | 0.003 | 0 | CPA3_Mast | GATA2 |
| 6 | 0 | 4.094019083 | 0.855 | 0.007 | 0 | CPA3_Mast | KIT |
| 7 | 0 | 3.933724095 | 0.837 | 0.131 | 0 | CPA3_Mast | CD69 |
| 8 | 0 | 3.877764496 | 0.874 | 0.024 | 0 | CPA3_Mast | CLU |

| | | | | | | | |
|---|---|---|---|---|---|---|---|
| 9 | 0 | 3.219982877 | 0.847 | 0.022 | 0 | CPA3_Mast | HPGDS |
| 10 | 0 | 3.20657519 | 0.829 | 0.029 | 0 | CPA3_Mast | VWA5A |
| 11 | 0 | 3.146662429 | 0.866 | 0.1 | 0 | CPA3_Mast | LTC4S |
| 12 | 0 | 2.939252088 | 0.719 | 0.003 | 0 | CPA3_Mast | HDC |
| 13 | 0 | 2.897004253 | 0.707 | 0.169 | 0 | CPA3_Mast | SOCS1 |
| 14 | 0 | 2.858994837 | 0.752 | 0.002 | 0 | CPA3_Mast | SLC18A2 |
| 15 | 0 | 2.811883855 | 0.672 | 0.001 | 0 | CPA3_Mast | MS4A2 |
| 16 | 0 | 2.630506098 | 0.497 | 0.036 | 0 | CPA3_Mast | CSF1 |
| 17 | 0 | 2.598886495 | 0.712 | 0.002 | 0 | CPA3_Mast | IL1RL1 |
| 18 | 0 | 2.446202557 | 0.556 | 0.007 | 0 | CPA3_Mast | SLC24A3 |
| 19 | 0 | 2.443545204 | 0.581 | 0.046 | 0 | CPA3_Mast | HPGD |
| 20 | 0 | 2.44327556 | 0.567 | 0.001 | 0 | CPA3_Mast | RHEX |
| 21 | 0 | 2.286331914 | 0.878 | 0.291 | 0 | CPA3_Mast | CD9 |
| 22 | 0 | 2.22589512 | 0.858 | 0.456 | 0 | CPA3_Mast | LMNA |
| 23 | 0 | 2.221518723 | 0.631 | 0.017 | 0 | CPA3_Mast | BACE2 |
| 24 | 0 | 2.150311629 | 0.608 | 0.002 | 0 | CPA3_Mast | MAOB |
| 25 | 0 | 2.089524729 | 0.873 | 0.445 | 0 | CPA3_Mast | LAPTM4A |
| 26 | 0 | 2.078140715 | 0.517 | 0.004 | 0 | CPA3_Mast | SIGLEC17P |
| 27 | 0 | 2.038814681 | 0.595 | 0.166 | 0 | CPA3_Mast | TSC22D1 |
| 28 | 0 | 2.033720945 | 0.82 | 0.428 | 0 | CPA3_Mast | NFKBIZ |
| 29 | 0 | 2.02913178 | 0.582 | 0.174 | 0 | CPA3_Mast | NSMCE1 |
| 30 | 9.7094E-271 | 2.528949964 | 0.715 | 0.398 | 3.1935E-266 | CPA3_Mast | TUBA1A |
| 31 | 0 | 3.23900778 | 0.441 | 0.006 | 0 | CXCR2_Neu | CMTM2 |
| 32 | 0 | 2.981525016 | 0.429 | 0.015 | 0 | CXCR2_Neu | FCGR3B |
| 33 | 0 | 2.84968948 | 0.272 | 0.002 | 0 | CXCR2_Neu | AL049651.1 |
| 34 | 2.0253E-229 | 2.766112637 | 0.658 | 0.091 | 6.6613E-225 | CXCR2_Neu | FP236383.1 |
| 35 | 1.2463E-211 | 3.719328325 | 0.748 | 0.178 | 4.0993E-207 | CXCR2_Neu | CSF3R |
| 36 | 3.24E-209 | 3.497805059 | 0.939 | 0.32 | 1.07E-204 | CXCR2_Neu | G0S2 |
| 37 | 5.47E-191 | 4.663403601 | 0.968 | 0.658 | 1.80E-186 | CXCR2_Neu | IFITM2 |
| 38 | 1.96E-185 | 3.532711619 | 0.875 | 0.299 | 6.44E-181 | CXCR2_Neu | CXCL8 |
| 39 | 3.63E-177 | 3.397029728 | 0.959 | 0.571 | 1.20E-172 | CXCR2_Neu | NAMPT |
| 40 | 4.54E-146 | 3.544889177 | 0.614 | 0.158 | 1.49E-141 | CXCR2_Neu | CPD |
| 41 | 1.46E-137 | 3.175145337 | 0.661 | 0.195 | 4.80E-133 | CXCR2_Neu | IFITM1 |
| 42 | 3.31E-136 | 3.417092634 | 0.562 | 0.131 | 1.09E-131 | CXCR2_Neu | IL1R2 |
| 43 | 3.72E-131 | 2.870598295 | 0.464 | 0.088 | 1.22E-126 | CXCR2_Neu | RIPOR2 |
| 44 | 1.50E-126 | 3.077438406 | 0.693 | 0.253 | 4.95E-122 | CXCR2_Neu | TREM1 |
| 45 | 2.71E-123 | 2.883939674 | 0.548 | 0.138 | 8.91E-119 | CXCR2_Neu | AQP9 |
| 46 | 1.90E-115 | 2.952297701 | 0.435 | 0.086 | 6.26E-111 | CXCR2_Neu | LUCAT1 |
| 47 | 1.07E-108 | 2.980277461 | 0.754 | 0.412 | 3.53E-104 | CXCR2_Neu | RNF149 |
| 48 | 2.68E-108 | 3.222371302 | 0.559 | 0.17 | 8.82E-104 | CXCR2_Neu | ARHGAP26 |
| 49 | 2.70E-105 | 2.607872373 | 0.838 | 0.621 | 8.89E-101 | CXCR2_Neu | LITAF |
| 50 | 3.84E-105 | 2.761889756 | 0.788 | 0.332 | 1.26E-100 | CXCR2_Neu | S100A8 |
| 51 | 7.75E-94 | 2.678426524 | 0.664 | 0.297 | 2.55E-89 | CXCR2_Neu | FPR1 |
| 52 | 3.65E-83 | 2.953768444 | 0.58 | 0.24 | 1.20E-78 | CXCR2_Neu | SMCHD1 |

| | | | | | | | |
|---|---|---|---|---|---|---|---|
| 53 | 1.36E-82 | 2.7701375 | 0.62 | 0.283 | 4.48E-78 | CXCR2_Neu | MXD1 |
| 54 | 2.10E-76 | 3.266512071 | 0.62 | 0.308 | 6.91E-72 | CXCR2_Neu | ACSL1 |
| 55 | 2.33E-75 | 3.119797098 | 0.42 | 0.121 | 7.65E-71 | CXCR2_Neu | DOCK4 |
| 56 | 2.18E-70 | 2.615567422 | 0.368 | 0.096 | 7.18E-66 | CXCR2_Neu | IL1RAP |
| 57 | 7.43E-49 | 2.598584401 | 0.481 | 0.23 | 2.44E-44 | CXCR2_Neu | PELI1 |
| 58 | 6.77E-47 | 2.552761136 | 0.409 | 0.167 | 2.23E-42 | CXCR2_Neu | NEDD9 |
| 59 | 2.59E-42 | 3.001738178 | 0.359 | 0.138 | 8.53E-38 | CXCR2_Neu | CAMK1D |
| 60 | 2.16E-41 | 2.549915174 | 0.504 | 0.289 | 7.10E-37 | CXCR2_Neu | PDE4B |
| 61 | 0.00E+00 | 2.669484744 | 0.744 | 0.074 | 0.00E+00 | VCAN_Mono | FCN1 |
| 62 | 0.00E+00 | 2.665762848 | 0.947 | 0.381 | 0.00E+00 | VCAN_Mono | S100A9 |
| 63 | 0.00E+00 | 2.432257298 | 0.891 | 0.262 | 0.00E+00 | VCAN_Mono | S100A8 |
| 64 | 0.00E+00 | 2.161863518 | 0.539 | 0.066 | 0.00E+00 | VCAN_Mono | S100A12 |
| 65 | 0.00E+00 | 1.9184019 | 0.672 | 0.126 | 0.00E+00 | VCAN_Mono | VCAN |
| 66 | 3.83E-298 | 1.710415215 | 0.368 | 0.033 | 1.26E-293 | VCAN_Mono | APOBEC3A |
| 67 | 7.90E-292 | 2.149185221 | 0.718 | 0.217 | 2.60E-287 | VCAN_Mono | THBS1 |
| 68 | 1.39E-270 | 1.494099511 | 0.685 | 0.218 | 4.58E-266 | VCAN_Mono | CSTA |
| 69 | 1.87E-267 | 1.96941241 | 0.624 | 0.162 | 6.15E-263 | VCAN_Mono | EREG |
| 70 | 3.88E-267 | 1.271773862 | 0.737 | 0.243 | 1.28E-262 | VCAN_Mono | FPR1 |
| 71 | 2.11E-242 | 1.301131683 | 0.85 | 0.366 | 6.95E-238 | VCAN_Mono | SERPINA1 |
| 72 | 5.01E-234 | 1.155917508 | 0.992 | 0.788 | 1.65E-229 | VCAN_Mono | S100A4 |
| 73 | 3.15E-224 | 1.199346986 | 0.749 | 0.258 | 1.03E-219 | VCAN_Mono | SLC11A1 |
| 74 | 3.13E-221 | 1.590244813 | 0.954 | 0.535 | 1.03E-216 | VCAN_Mono | LYZ |
| 75 | 2.01E-203 | 1.632228895 | 0.938 | 0.723 | 6.60E-199 | VCAN_Mono | TIMP1 |
| 76 | 1.31E-201 | 1.278951719 | 0.453 | 0.097 | 4.31E-197 | VCAN_Mono | CD300E |
| 77 | 1.77E-192 | 1.134421148 | 0.921 | 0.614 | 5.82E-188 | VCAN_Mono | COTL1 |
| 78 | 2.94E-188 | 1.14780848 | 0.672 | 0.239 | 9.66E-184 | VCAN_Mono | SLC25A37 |
| 79 | 3.68E-188 | 1.375782064 | 0.737 | 0.333 | 1.21E-183 | VCAN_Mono | UPP1 |
| 80 | 7.35E-171 | 1.380687266 | 0.619 | 0.233 | 2.42E-166 | VCAN_Mono | NLRP3 |
| 81 | 3.74E-157 | 1.154088392 | 0.896 | 0.551 | 1.23E-152 | VCAN_Mono | CTSS |
| 82 | 7.12E-157 | 1.162888611 | 0.838 | 0.464 | 2.34E-152 | VCAN_Mono | LST1 |
| 83 | 1.39E-149 | 1.302415888 | 0.63 | 0.245 | 4.56E-145 | VCAN_Mono | IL1B |
| 84 | 3.66E-149 | 1.176510142 | 0.332 | 0.068 | 1.20E-144 | VCAN_Mono | MCEMP1 |
| 85 | 1.04E-143 | 1.24083296 | 0.549 | 0.204 | 3.42E-139 | VCAN_Mono | HBEGF |
| 86 | 1.42E-132 | 1.362899 | 0.564 | 0.242 | 4.68E-128 | VCAN_Mono | PPIF |
| 87 | 6.04E-132 | 1.226722033 | 0.716 | 0.427 | 1.99E-127 | VCAN_Mono | STXBP2 |
| 88 | 2.10E-125 | 1.261503703 | 0.4 | 0.122 | 6.90E-121 | VCAN_Mono | OLR1 |
| 89 | 5.50E-116 | 1.21406046 | 0.284 | 0.064 | 1.81E-111 | VCAN_Mono | RETN |
| 90 | 9.37E-40 | 1.493979227 | 0.173 | 0.059 | 3.08E-35 | VCAN_Mono | IL8 |
| 91 | 0.00E+00 | 4.756231515 | 0.526 | 0.033 | 0.00E+00 | Angio-TAM | CXCL5 |
| 92 | 0.00E+00 | 3.35692631 | 0.587 | 0.088 | 0.00E+00 | Angio-TAM | CCL20 |
| 93 | 0.00E+00 | 2.992784756 | 0.952 | 0.304 | 0.00E+00 | Angio-TAM | C15orf48 |
| 94 | 0.00E+00 | 2.638567319 | 0.759 | 0.18 | 0.00E+00 | Angio-TAM | IL1RN |
| 95 | 0.00E+00 | 2.588459667 | 0.977 | 0.572 | 0.00E+00 | Angio-TAM | SOD2 |
| 96 | 0.00E+00 | 2.091672268 | 1 | 0.998 | 0.00E+00 | Angio-TAM | FTH1 |

| | | | | | | | |
|---|---|---|---|---|---|---|---|
| 97 | 7.12E-292 | 3.443251237 | 0.719 | 0.191 | 2.34E-287 | Angio-TAM | CXCL3 |
| 98 | 1.24E-261 | 2.825683685 | 0.77 | 0.243 | 4.07E-257 | Angio-TAM | IL1B |
| 99 | 5.14E-259 | 3.290385835 | 0.487 | 0.078 | 1.6917E-254 | Angio-TAM | SPP1 |
| 100 | 1.2723E-245 | 2.45723144 | 0.794 | 0.269 | 4.1849E-241 | Angio-TAM | CXCL8 |
| 101 | 3.7732E-240 | 2.107906403 | 0.937 | 0.727 | 1.241E-235 | Angio-TAM | MIF |
| 102 | 6.1145E-229 | 1.749807806 | 0.99 | 0.912 | 2.0111E-224 | Angio-TAM | GAPDH |
| 103 | 1.0622E-224 | 2.597549066 | 0.76 | 0.269 | 3.4937E-220 | Angio-TAM | CXCL2 |
| 104 | 1.30E-221 | 2.605390822 | 0.951 | 0.731 | 4.28E-217 | Angio-TAM | TIMP1 |
| 105 | 1.65E-215 | 1.724114571 | 0.291 | 0.026 | 5.43E-211 | Angio-TAM | IL1A |
| 106 | 3.19E-213 | 2.074308295 | 0.369 | 0.049 | 1.05E-208 | Angio-TAM | TNFAIP6 |
| 107 | 5.20E-199 | 1.968102039 | 0.923 | 0.659 | 1.71E-194 | Angio-TAM | CSTB |
| 108 | 4.82E-191 | 2.624952133 | 0.736 | 0.274 | 1.58E-186 | Angio-TAM | CCL3 |
| 109 | 2.56E-190 | 3.475715888 | 0.2 | 0.01 | 8.43E-186 | Angio-TAM | PPBP |
| 110 | 6.43E-177 | 1.795971986 | 0.619 | 0.234 | 2.11E-172 | Angio-TAM | C1orf122 |
| 111 | 1.42E-145 | 3.372345153 | 0.832 | 0.587 | 4.67E-141 | Angio-TAM | MT2A |
| 112 | 3.91E-141 | 2.715408905 | 0.551 | 0.189 | 1.29E-136 | Angio-TAM | CCL3L1 |
| 113 | 2.87E-138 | 2.654050702 | 0.397 | 0.094 | 9.45E-134 | Angio-TAM | CXCL1 |
| 114 | 1.48E-128 | 2.197100909 | 0.484 | 0.166 | 4.88E-124 | Angio-TAM | MT1F |
| 115 | 2.05E-116 | 2.342782557 | 0.434 | 0.139 | 6.76E-112 | Angio-TAM | MT1E |
| 116 | 1.80E-100 | 2.9215094 | 0.574 | 0.286 | 5.92E-96 | Angio-TAM | MT1X |
| 117 | 6.66E-96 | 1.894171304 | 0.651 | 0.316 | 2.19E-91 | Angio-TAM | CCL4 |
| 118 | 1.73E-90 | 2.795550569 | 0.36 | 0.115 | 5.68E-86 | Angio-TAM | MT1G |
| 119 | 2.52E-68 | 2.552280242 | 0.178 | 0.036 | 8.29E-64 | Angio-TAM | MT1H |
| 120 | 6.69E-46 | 1.743326881 | 0.436 | 0.242 | 2.20E-41 | Angio-TAM | PTGS2 |
| 121 | 0.00E+00 | 3.101309441 | 0.879 | 0.263 | 0.00E+00 | TREM2_TAM | APOE |
| 122 | 0.00E+00 | 3.005478233 | 0.836 | 0.226 | 0.00E+00 | TREM2_TAM | APOC1 |
| 123 | 0.00E+00 | 2.153691189 | 0.966 | 0.47 | 0.00E+00 | TREM2_TAM | CTSD |
| 124 | 0.00E+00 | 2.146437624 | 0.861 | 0.295 | 0.00E+00 | TREM2_TAM | ACP5 |
| 125 | 0.00E+00 | 2.118286511 | 0.996 | 0.648 | 0.00E+00 | TREM2_TAM | CTSB |
| 126 | 0.00E+00 | 2.109834421 | 0.919 | 0.237 | 0.00E+00 | TREM2_TAM | C1QB |
| 127 | 0.00E+00 | 1.988908673 | 0.913 | 0.235 | 0.00E+00 | TREM2_TAM | C1QC |
| 128 | 0.00E+00 | 1.939739537 | 0.928 | 0.259 | 0.00E+00 | TREM2_TAM | C1QA |
| 129 | 0.00E+00 | 1.903403664 | 0.964 | 0.479 | 0.00E+00 | TREM2_TAM | CTSZ |
| 130 | 0.00E+00 | 1.870172512 | 0.693 | 0.089 | 0.00E+00 | TREM2_TAM | GPNMB |
| 131 | 0.00E+00 | 1.836210609 | 0.783 | 0.252 | 0.00E+00 | TREM2_TAM | FABP5 |
| 132 | 0.00E+00 | 1.83372099 | 0.848 | 0.269 | 0.00E+00 | TREM2_TAM | CTSL |
| 133 | 0.00E+00 | 1.741959707 | 1 | 0.993 | 0.00E+00 | TREM2_TAM | FTL |
| 134 | 0.00E+00 | 1.674267907 | 0.842 | 0.299 | 0.00E+00 | TREM2_TAM | PLD3 |
| 135 | 0.00E+00 | 1.612926661 | 0.886 | 0.433 | 0.00E+00 | TREM2_TAM | CD68 |
| 136 | 0.00E+00 | 1.546102325 | 0.556 | 0.054 | 0.00E+00 | TREM2_TAM | TREM2 |
| 137 | 0.00E+00 | 1.539643239 | 0.763 | 0.195 | 0.00E+00 | TREM2_TAM | LGMN |
| 138 | 0.00E+00 | 1.513398141 | 0.655 | 0.127 | 0.00E+00 | TREM2_TAM | PLA2G7 |
| 139 | 0.00E+00 | 1.470175619 | 0.877 | 0.393 | 0.00E+00 | TREM2_TAM | CTSC |
| 140 | 0.00E+00 | 1.463099598 | 0.932 | 0.511 | 0.00E+00 | TREM2_TAM | GRN |

| | | | | | | | |
|---|---|---|---|---|---|---|---|
| 141 | 0.00E+00 | 1.432629669 | 0.99 | 0.808 | 0.00E+00 | TREM2_TAM | PSAP |
| 142 | 2.56E-297 | 1.353037469 | 0.646 | 0.174 | 8.43E-293 | TREM2_TAM | LIPA |
| 143 | 2.52E-293 | 2.589442063 | 0.417 | 0.056 | 8.29E-289 | TREM2_TAM | CCL18 |
| 144 | 5.50E-272 | 1.331075316 | 0.93 | 0.483 | 1.81E-267 | TREM2_TAM | ANXA2 |
| 145 | 6.53E-272 | 1.456030051 | 0.951 | 0.625 | 2.15E-267 | TREM2_TAM | LGALS3 |
| 146 | 1.28E-230 | 2.432031035 | 0.338 | 0.046 | 4.21E-226 | TREM2_TAM | MMP12 |
| 147 | 8.77E-230 | 1.483470693 | 0.846 | 0.433 | 2.88E-225 | TREM2_TAM | HLA-DRB5 |
| 148 | 1.22E-200 | 1.561695262 | 0.415 | 0.089 | 4.03E-196 | TREM2_TAM | MMP9 |
| 149 | 9.61E-128 | 1.956347675 | 0.236 | 0.044 | 3.16E-123 | TREM2_TAM | FN1 |
| 150 | 3.63E-50 | 1.44376196 | 0.279 | 0.125 | 1.19E-45 | TREM2_TAM | IFI27 |
| 151 | 6.3679E-145 | 2.00534569 | 0.922 | 0.335 | 2.0945E-140 | C1QC_TAM | C1QC |
| 152 | 2.4712E-123 | 2.093954156 | 0.629 | 0.164 | 8.1279E-119 | C1QC_TAM | SELENOP |
| 153 | 9.3372E-116 | 1.653047935 | 0.909 | 0.359 | 3.0711E-111 | C1QC_TAM | C1QA |
| 154 | 3.5011E-111 | 1.696472244 | 0.875 | 0.341 | 1.1516E-106 | C1QC_TAM | C1QB |
| 155 | 2.63761E-98 | 1.375023742 | 1 | 0.994 | 8.67537E-94 | C1QC_TAM | FTL |
| 156 | 1.89633E-51 | 1.256956405 | 0.89 | 0.657 | 6.23721E-47 | C1QC_TAM | HLA-DPA1 |
| 157 | 3.91E-51 | 1.827761024 | 0.303 | 0.084 | 1.29E-46 | C1QC_TAM | DNASE1L3 |
| 158 | 1.48E-48 | 1.373453393 | 0.747 | 0.64 | 4.88E-44 | C1QC_TAM | PRDX1 |
| 159 | 3.47E-48 | 1.804912014 | 0.46 | 0.197 | 1.14E-43 | C1QC_TAM | SLC40A1 |
| 160 | 2.42E-43 | 1.539907278 | 0.603 | 0.329 | 7.97E-39 | C1QC_TAM | APOC1 |
| 161 | 2.32E-42 | 1.533750111 | 0.765 | 0.675 | 7.64E-38 | C1QC_TAM | TUBA1B |
| 162 | 8.59E-37 | 1.28597267 | 0.645 | 0.367 | 2.83E-32 | C1QC_TAM | APOE |
| 163 | 1.57E-36 | 1.929178993 | 0.413 | 0.189 | 5.18E-32 | C1QC_TAM | IGKV1-5 |
| 164 | 2.83E-31 | 1.304847532 | 0.386 | 0.188 | 9.30E-27 | C1QC_TAM | GGTA1P |
| 165 | 2.70E-29 | 1.200595092 | 0.629 | 0.478 | 8.89E-25 | C1QC_TAM | TMEM176B |
| 166 | 2.69E-28 | 1.307355529 | 0.624 | 0.479 | 8.84E-24 | C1QC_TAM | CTSC |
| 167 | 9.02E-26 | 1.161032686 | 0.65 | 0.513 | 2.97E-21 | C1QC_TAM | CD68 |
| 168 | 1.24E-24 | 2.365454993 | 0.198 | 0.066 | 4.09E-20 | C1QC_TAM | IGHV4-34 |
| 169 | 2.46E-24 | 1.214870585 | 0.232 | 0.089 | 8.10E-20 | C1QC_TAM | ADAMDEC1 |
| 170 | 5.79E-21 | 1.187325748 | 0.535 | 0.408 | 1.90E-16 | C1QC_TAM | YWHAH |
| 171 | 2.47E-18 | 1.445868737 | 0.337 | 0.189 | 8.11E-14 | C1QC_TAM | JCHAIN |
| 172 | 6.41E-18 | 2.297015535 | 0.292 | 0.149 | 2.11E-13 | C1QC_TAM | IGLV2-14 |
| 173 | 2.61E-15 | 1.644135177 | 0.136 | 0.046 | 8.58E-11 | C1QC_TAM | MT1H |
| 174 | 1.09E-13 | 1.305436589 | 0.366 | 0.26 | 3.57E-09 | C1QC_TAM | LIPA |
| 175 | 5.24E-11 | 2.048077818 | 0.24 | 0.136 | 1.72E-06 | C1QC_TAM | MT1G |
| 176 | 9.75E-10 | 1.195783935 | 0.436 | 0.35 | 3.21E-05 | C1QC_TAM | FABP5 |
| 177 | 1.37E-08 | 1.160593596 | 0.185 | 0.098 | 4.49E-04 | C1QC_TAM | MMP12 |
| 178 | 1.71E-08 | 1.263871541 | 0.191 | 0.109 | 5.64E-04 | C1QC_TAM | IGLV1-40 |
| 179 | 2.28E-08 | 2.893029328 | 0.172 | 0.094 | 7.49E-04 | C1QC_TAM | IGKV3-15 |
| 180 | 3.86E-08 | 1.993118654 | 0.164 | 0.089 | 1.27E-03 | C1QC_TAM | IGLV2-23 |
| 181 | 0.00E+00 | 3.365485489 | 0.625 | 0.088 | 0.00E+00 | FOLR2_TAM | F13A1 |
| 182 | 0.00E+00 | 3.117595936 | 0.807 | 0.139 | 0.00E+00 | FOLR2_TAM | SELENOP |
| 183 | 0.00E+00 | 2.467951901 | 0.355 | 0.016 | 0.00E+00 | FOLR2_TAM | LYVE1 |
| 184 | 6.72E-252 | 3.561385427 | 0.758 | 0.19 | 2.21E-247 | FOLR2_TAM | RNASE1 |

| | | | | | | | |
|---|---|---|---|---|---|---|---|
| 185 | 1.12E-247 | 2.441499 | 0.606 | 0.108 | 3.70E-243 | FOLR2_TAM | FOLR2 |
| 186 | 1.75E-235 | 2.795458405 | 0.797 | 0.255 | 5.77E-231 | FOLR2_TAM | STAB1 |
| 187 | 3.63E-203 | 1.98003691 | 0.917 | 0.346 | 1.19E-198 | FOLR2_TAM | C1QA |
| 188 | 4.50E-192 | 2.536913138 | 0.634 | 0.159 | 1.48E-187 | FOLR2_TAM | PLTP |
| 189 | 3.47E-189 | 2.703164573 | 0.644 | 0.175 | 1.14E-184 | FOLR2_TAM | PDK4 |
| 190 | 1.45E-185 | 1.8342437 | 0.893 | 0.327 | 4.79E-181 | FOLR2_TAM | C1QB |
| 191 | 1.18E-183 | 1.745655298 | 0.895 | 0.324 | 3.87E-179 | FOLR2_TAM | C1QC |
| 192 | 4.59E-153 | 1.641026385 | 0.865 | 0.423 | 1.51E-148 | FOLR2_TAM | MS4A6A |
| 193 | 2.66E-143 | 2.067290728 | 0.6 | 0.189 | 8.75E-139 | FOLR2_TAM | MRC1 |
| 194 | 1.20E-137 | 1.684721629 | 0.602 | 0.179 | 3.93E-133 | FOLR2_TAM | SLC40A1 |
| 195 | 4.47E-136 | 1.809226892 | 0.687 | 0.27 | 1.47E-131 | FOLR2_TAM | DAB2 |
| 196 | 3.45E-130 | 1.691278581 | 0.85 | 0.512 | 1.13E-125 | FOLR2_TAM | FCGRT |
| 197 | 3.46E-130 | 1.658788211 | 0.557 | 0.17 | 1.14E-125 | FOLR2_TAM | GGTA1P |
| 198 | 1.35E-129 | 1.915300499 | 0.722 | 0.316 | 4.46E-125 | FOLR2_TAM | CD163 |
| 199 | 3.08E-128 | 1.547543578 | 0.39 | 0.08 | 1.01E-123 | FOLR2_TAM | LILRB5 |
| 200 | 3.26E-128 | 1.716380503 | 0.795 | 0.425 | 1.07E-123 | FOLR2_TAM | MS4A7 |
| 201 | 1.9869E-126 | 1.908095098 | 0.642 | 0.246 | 6.5351E-122 | FOLR2_TAM | MS4A4A |
| 202 | 5.2297E-116 | 1.717234506 | 0.675 | 0.275 | 1.7201E-111 | FOLR2_TAM | LGMN |
| 203 | 8.4274E-115 | 1.527610775 | 0.548 | 0.182 | 2.7718E-110 | FOLR2_TAM | SLCO2B1 |
| 204 | 1.6287E-113 | 1.51189723 | 0.553 | 0.179 | 5.3571E-109 | FOLR2_TAM | A2M |
| 205 | 3.9079E-103 | 1.609259187 | 0.698 | 0.331 | 1.28534E-98 | FOLR2_TAM | MAFB |
| 206 | 3.84589E-99 | 1.741090153 | 0.713 | 0.361 | 1.26495E-94 | FOLR2_TAM | FGL2 |
| 207 | 5.80678E-90 | 1.770668678 | 0.689 | 0.36 | 1.90991E-85 | FOLR2_TAM | RHOB |
| 208 | 1.45394E-84 | 1.72058755 | 0.644 | 0.319 | 4.78215E-80 | FOLR2_TAM | CFD |
| 209 | 1.14916E-71 | 1.592451465 | 0.235 | 0.047 | 3.77969E-67 | FOLR2_TAM | PGA3 |
| 210 | 1.67436E-56 | 1.503652871 | 0.636 | 0.401 | 5.50713E-52 | FOLR2_TAM | ATF3 |
| 211 | 0 | 1.925334334 | 0.474 | 0.021 | 0 | CD1C_DC | CD1C |
| 212 | 0 | 1.421872795 | 0.364 | 0.018 | 0 | CD1C_DC | CD1E |
| 213 | 2.35E-286 | 2.05439973 | 0.993 | 0.606 | 7.74E-282 | CD1C_DC | HLA-DPB1 |
| 214 | 5.38E-282 | 1.731365358 | 0.594 | 0.103 | 1.77E-277 | CD1C_DC | CLEC10A |
| 215 | 9.09E-259 | 1.838719975 | 0.997 | 0.634 | 2.99E-254 | CD1C_DC | HLA-DPA1 |
| 216 | 2.36E-252 | 1.994981115 | 0.977 | 0.525 | 7.77E-248 | CD1C_DC | HLA-DQB1 |
| 217 | 1.61E-247 | 1.790241534 | 0.999 | 0.672 | 5.28E-243 | CD1C_DC | HLA-DRA |
| 218 | 7.41E-221 | 1.723015224 | 0.994 | 0.695 | 2.44E-216 | CD1C_DC | HLA-DRB1 |
| 219 | 2.99E-220 | 1.71934772 | 0.97 | 0.483 | 9.83E-216 | CD1C_DC | HLA-DQA1 |
| 220 | 1.10E-213 | 1.847584095 | 0.473 | 0.084 | 3.61E-209 | CD1C_DC | NAPSB |
| 221 | 1.92E-200 | 1.608818513 | 0.741 | 0.244 | 6.30E-196 | CD1C_DC | PPA1 |
| 222 | 1.96E-192 | 1.433193456 | 1 | 0.834 | 6.45E-188 | CD1C_DC | CD74 |
| 223 | 6.48E-174 | 1.41459782 | 0.868 | 0.378 | 2.13E-169 | CD1C_DC | LSP1 |
| 224 | 1.54E-162 | 1.447417521 | 0.751 | 0.282 | 5.06E-158 | CD1C_DC | CPVL |
| 225 | 1.89E-157 | 1.041452408 | 0.787 | 0.285 | 6.23E-153 | CD1C_DC | HLA-DMB |
| 226 | 1.87E-154 | 1.111053975 | 0.513 | 0.129 | 6.17E-150 | CD1C_DC | JAML |
| 227 | 1.16E-140 | 1.036309801 | 0.916 | 0.466 | 3.81E-136 | CD1C_DC | HLA-DMA |
| 228 | 3.06E-140 | 1.104520575 | 0.834 | 0.398 | 1.01E-135 | CD1C_DC | CORO1A |

| | | | | | | | |
|---|---|---|---|---|---|---|---|
| 229 | 1.48E-137 | 1.489541513 | 0.847 | 0.449 | 4.88E-133 | CD1C_DC | GPR183 |
| 230 | 4.73E-136 | 1.501093755 | 0.98 | 0.859 | 1.55E-131 | CD1C_DC | CST3 |
| 231 | 2.74E-135 | 1.182600936 | 0.486 | 0.124 | 9.01E-131 | CD1C_DC | CST7 |
| 232 | 6.83E-135 | 1.221651717 | 0.616 | 0.202 | 2.24E-130 | CD1C_DC | LGALS2 |
| 233 | 1.10E-134 | 1.569357879 | 0.402 | 0.091 | 3.62E-130 | CD1C_DC | FCER1A |
| 234 | 3.10E-105 | 1.061248045 | 0.22 | 0.033 | 1.02E-100 | CD1C_DC | AMICA1 |
| 235 | 1.10E-103 | 1.074104267 | 0.942 | 0.628 | 3.63E-99 | CD1C_DC | COTL1 |
| 236 | 1.21E-97 | 1.126773012 | 0.327 | 0.076 | 3.97E-93 | CD1C_DC | LTB |
| 237 | 6.34E-97 | 1.225829808 | 0.436 | 0.138 | 2.08E-92 | CD1C_DC | HLA-DQA2 |
| 238 | 1.68E-87 | 1.248872711 | 0.619 | 0.273 | 5.54E-83 | CD1C_DC | INSIG1 |
| 239 | 1.52E-76 | 1.140289924 | 0.807 | 0.47 | 5.01E-72 | CD1C_DC | HERPUD1 |
| 240 | 4.14E-60 | 1.701517003 | 0.256 | 0.072 | 1.36E-55 | CD1C_DC | S100B |
| 241 | 0.00E+00 | 4.622910227 | 0.778 | 0.016 | 0.00E+00 | LAMP3_DC | CCL22 |
| 242 | 0.00E+00 | 4.362890853 | 0.706 | 0.011 | 0.00E+00 | LAMP3_DC | CCL19 |
| 243 | 0.00E+00 | 3.989928978 | 0.937 | 0.035 | 0.00E+00 | LAMP3_DC | CCR7 |
| 244 | 0.00E+00 | 2.769251421 | 0.825 | 0.035 | 0.00E+00 | LAMP3_DC | LAMP3 |
| 245 | 0.00E+00 | 2.2782058 | 0.698 | 0.019 | 0.00E+00 | LAMP3_DC | LAD1 |
| 246 | 1.54E-301 | 4.393316234 | 0.929 | 0.067 | 5.07E-297 | LAMP3_DC | FSCN1 |
| 247 | 7.08E-237 | 3.043031887 | 0.754 | 0.051 | 2.33E-232 | LAMP3_DC | IDO1 |
| 248 | 2.67E-233 | 2.25022211 | 0.579 | 0.028 | 8.80E-229 | LAMP3_DC | RAMP1 |
| 249 | 3.34E-207 | 2.894844489 | 0.381 | 0.012 | 1.10E-202 | LAMP3_DC | CCL17 |
| 250 | 1.1011E-181 | 2.251711363 | 0.54 | 0.032 | 3.6217E-177 | LAMP3_DC | UBD |
| 251 | 1.8798E-178 | 1.96110567 | 0.802 | 0.083 | 6.1829E-174 | LAMP3_DC | DAPP1 |
| 252 | 2.5957E-163 | 2.834090108 | 0.516 | 0.034 | 8.5376E-159 | LAMP3_DC | WFDC21P |
| 253 | 3.83E-105 | 2.788924909 | 0.746 | 0.129 | 1.26E-100 | LAMP3_DC | EBI3 |
| 254 | 8.44E-97 | 2.835947064 | 0.746 | 0.139 | 2.78E-92 | LAMP3_DC | DUSP5 |
| 255 | 5.90E-91 | 1.949595965 | 0.833 | 0.198 | 1.94E-86 | LAMP3_DC | RAB9A |
| 256 | 1.66E-90 | 1.816532102 | 0.722 | 0.133 | 5.46E-86 | LAMP3_DC | MGLL |
| 257 | 9.05E-86 | 2.96776754 | 0.921 | 0.299 | 2.98E-81 | LAMP3_DC | BIRC3 |
| 258 | 8.26E-85 | 1.898466677 | 0.706 | 0.127 | 2.72E-80 | LAMP3_DC | IL32 |
| 259 | 2.14E-81 | 2.397939445 | 0.73 | 0.149 | 7.05E-77 | LAMP3_DC | CST7 |
| 260 | 2.36E-77 | 2.955880362 | 0.841 | 0.23 | 7.76E-73 | LAMP3_DC | MARCKSL1 |
| 261 | 2.27E-76 | 1.82016023 | 0.762 | 0.181 | 7.48E-72 | LAMP3_DC | GRSF1 |
| 262 | 7.68E-73 | 1.987715738 | 0.786 | 0.176 | 2.53E-68 | LAMP3_DC | IL7R |
| 263 | 1.61E-72 | 2.562936193 | 0.952 | 0.416 | 5.28E-68 | LAMP3_DC | LSP1 |
| 264 | 5.85E-66 | 2.238547383 | 0.849 | 0.282 | 1.93E-61 | LAMP3_DC | PPA1 |
| 265 | 7.25E-66 | 3.496237881 | 0.984 | 0.701 | 2.38E-61 | LAMP3_DC | TXN |
| 266 | 1.48E-56 | 1.851948334 | 0.778 | 0.257 | 4.87E-52 | LAMP3_DC | ANXA6 |
| 267 | 1.97E-55 | 2.581550442 | 0.976 | 0.541 | 6.47E-51 | LAMP3_DC | CRIP1 |
| 268 | 9.18E-45 | 1.837683463 | 0.857 | 0.475 | 3.02E-40 | LAMP3_DC | SYNGR2 |
| 269 | 2.14E-41 | 1.829523017 | 0.31 | 0.047 | 7.02E-37 | LAMP3_DC | PPP1R14A |
| 270 | 1.08E-38 | 1.84627313 | 0.992 | 0.987 | 3.56E-34 | LAMP3_DC | ACTB |
| 271 | 0.00E+00 | 5.662472408 | 0.982 | 0.021 | 0.00E+00 | GZMB_pDC | GZMB |
| 272 | 0.00E+00 | 3.632427308 | 0.842 | 0.014 | 0.00E+00 | GZMB_pDC | CLIC3 |

| | | | | | | | |
|---|---|---|---|---|---|---|---|
| 273 | 0.00E+00 | 2.869398919 | 0.456 | 0.004 | 0.00E+00 | GZMB_pDC | TCL1A |
| 274 | 0.00E+00 | 2.830608888 | 0.702 | 0.004 | 0.00E+00 | GZMB_pDC | LILRA4 |
| 275 | 0.00E+00 | 2.770228125 | 0.789 | 0.014 | 0.00E+00 | GZMB_pDC | CXCR3 |
| 276 | 0.00E+00 | 2.275074351 | 0.544 | 0.001 | 0.00E+00 | GZMB_pDC | SCT |
| 277 | 0.00E+00 | 2.24170672 | 0.807 | 0.005 | 0.00E+00 | GZMB_pDC | SMPD3 |
| 278 | 3.43E-256 | 2.883226751 | 0.895 | 0.033 | 1.13E-251 | GZMB_pDC | TSPAN13 |
| 279 | 5.19E-232 | 2.593806738 | 0.772 | 0.026 | 1.71E-227 | GZMB_pDC | SPIB |
| 280 | 7.17E-213 | 2.346621065 | 0.737 | 0.025 | 2.36E-208 | GZMB_pDC | MZB1 |
| 281 | 2.24E-184 | 3.06809615 | 0.842 | 0.042 | 7.38E-180 | GZMB_pDC | C12orf75 |
| 282 | 1.12E-131 | 2.41322109 | 0.912 | 0.071 | 3.69E-127 | GZMB_pDC | PTPRCAP |
| 283 | 5.79E-122 | 3.618707662 | 0.316 | 0.008 | 1.90E-117 | GZMB_pDC | PTGDS |
| 284 | 2.94E-116 | 2.958311387 | 0.789 | 0.061 | 9.69E-112 | GZMB_pDC | IRF4 |
| 285 | 1.57E-108 | 3.152871366 | 0.895 | 0.09 | 5.15E-104 | GZMB_pDC | PLD4 |
| 286 | 3.96E-105 | 2.690948022 | 0.93 | 0.101 | 1.30E-100 | GZMB_pDC | TCF4 |
| 287 | 3.89E-90 | 3.384347345 | 0.93 | 0.117 | 1.28E-85 | GZMB_pDC | PLAC8 |
| 288 | 2.07E-73 | 2.25954995 | 0.86 | 0.124 | 6.82E-69 | GZMB_pDC | CCDC50 |
| 289 | 6.30E-71 | 3.125323572 | 0.772 | 0.095 | 2.07E-66 | GZMB_pDC | LTB |
| 290 | 6.60E-69 | 2.323710347 | 0.825 | 0.119 | 2.17E-64 | GZMB_pDC | IL3RA |
| 291 | 7.41E-60 | 3.466456967 | 0.965 | 0.225 | 2.44E-55 | GZMB_pDC | IRF7 |
| 292 | 4.00E-59 | 3.565049334 | 0.579 | 0.065 | 1.31E-54 | GZMB_pDC | IGJ |
| 293 | 6.46E-57 | 2.265687318 | 0.807 | 0.134 | 2.12E-52 | GZMB_pDC | SERPINF1 |
| 294 | 5.69E-49 | 2.563442117 | 0.912 | 0.224 | 1.87E-44 | GZMB_pDC | ITM2C |
| 295 | 9.12E-46 | 2.63722283 | 0.825 | 0.18 | 3.00E-41 | GZMB_pDC | LDLRAD4 |
| 296 | 1.28E-42 | 2.547707983 | 0.877 | 0.237 | 4.20E-38 | GZMB_pDC | IRF8 |
| 297 | 3.43E-40 | 2.444742055 | 0.737 | 0.145 | 1.13E-35 | GZMB_pDC | DUSP5 |
| 298 | 3.20E-37 | 2.357776705 | 0.965 | 0.384 | 1.05E-32 | GZMB_pDC | CXCR4 |
| 299 | 1.27E-35 | 2.34377251 | 0.93 | 0.331 | 4.17E-31 | GZMB_pDC | PPP1R14B |
| 300 | 1.25E-32 | 2.859655587 | 0.947 | 0.484 | 4.10E-28 | GZMB_pDC | GPR183 |

Supplementary Table S6 DEGs of cancer associated fibroblasts subclusters

| | p_val | avg_log2FC | pct.1 | pct.2 | p_val_adj | cluster | gene |
|---|---|---|---|---|---|---|---|
| 1 | 0.00E+00 | 3.635211574 | 0.557 | 0.01 | 0.00E+00 | iCAF | CXCL5 |
| 2 | 0.00E+00 | 2.408238443 | 0.416 | 0.007 | 0.00E+00 | iCAF | IL24 |
| 3 | 3.14E-271 | 1.904239634 | 0.489 | 0.023 | 1.03E-266 | iCAF | C15orf48 |
| 4 | 7.24E-253 | 2.094899989 | 0.235 | 0.002 | 2.38E-248 | iCAF | CSF3 |
| 5 | 3.78E-240 | 3.105947872 | 0.706 | 0.069 | 1.24E-235 | iCAF | CXCL8 |
| 6 | 2.51E-225 | 2.910460622 | 0.837 | 0.124 | 8.24E-221 | iCAF | COL7A1 |
| 7 | 3.07E-197 | 3.907237733 | 0.633 | 0.071 | 1.01E-192 | iCAF | MMP1 |
| 8 | 1.51E-189 | 1.861069804 | 0.697 | 0.093 | 4.96E-185 | iCAF | SLC16A3 |
| 9 | 8.73E-184 | 4.406802711 | 0.529 | 0.05 | 2.87E-179 | iCAF | MMP3 |
| 10 | 6.05E-171 | 3.010181625 | 0.692 | 0.098 | 1.99E-166 | iCAF | CHI3L1 |
| 11 | 1.19E-164 | 2.371482699 | 0.326 | 0.018 | 3.90E-160 | iCAF | SAA1 |
| 12 | 1.44E-148 | 2.095238831 | 0.43 | 0.039 | 4.75E-144 | iCAF | CXCL6 |
| 13 | 2.10E-129 | 1.860585782 | 0.529 | 0.072 | 6.92E-125 | iCAF | MT1F |

| | | | | | | | |
|---|---|---|---|---|---|---|---|
| 14 | 7.31E-117 | 1.943559409 | 0.783 | 0.193 | 2.41E-112 | iCAF | TMEM158 |
| 15 | 8.22E-114 | 3.480670552 | 0.674 | 0.144 | 2.71E-109 | iCAF | CXCL1 |
| 16 | 1.35E-111 | 2.70865655 | 1 | 0.99 | 4.43E-107 | iCAF | FTH1 |
| 17 | 2.78E-103 | 1.939323873 | 0.991 | 0.87 | 9.15E-99 | iCAF | MIF |
| 18 | 8.16E-100 | 2.059167301 | 0.995 | 0.937 | 2.69E-95 | iCAF | GAPDH |
| 19 | 1.13E-99 | 2.526430012 | 0.968 | 0.551 | 3.72E-95 | iCAF | SOD2 |
| 20 | 3.63E-95 | 2.077607019 | 0.457 | 0.071 | 1.19E-90 | iCAF | CXCL3 |
| 21 | 1.14E-90 | 2.895687702 | 0.995 | 0.828 | 3.75E-86 | iCAF | MT2A |
| 22 | 2.23E-85 | 1.952876469 | 0.751 | 0.241 | 7.34E-81 | iCAF | ADM |
| 23 | 7.73E-78 | 1.902041279 | 0.57 | 0.133 | 2.54E-73 | iCAF | G0S2 |
| 24 | 1.15E-75 | 1.816882585 | 0.946 | 0.748 | 3.79E-71 | iCAF | TPI1 |
| 25 | 6.17E-74 | 1.818978133 | 0.864 | 0.431 | 2.03E-69 | iCAF | COL12A1 |
| 26 | 7.82E-67 | 2.294251387 | 0.606 | 0.174 | 2.57E-62 | iCAF | ANGPTL4 |
| 27 | 7.23E-58 | 2.324432094 | 0.914 | 0.605 | 2.38E-53 | iCAF | MT1E |
| 28 | 7.06E-51 | 2.119188783 | 0.701 | 0.304 | 2.32E-46 | iCAF | PLAU |
| 29 | 1.50E-48 | 2.144691588 | 0.878 | 0.514 | 4.94E-44 | iCAF | MT1X |
| 30 | 1.75E-48 | 2.063142967 | 0.805 | 0.454 | 5.75E-44 | iCAF | DDIT4 |
| 31 | 0.00E+00 | 3.798114756 | 0.717 | 0.183 | 0.00E+00 | mCAF1 | POSTN |
| 32 | 0.00E+00 | 2.815122648 | 0.801 | 0.216 | 0.00E+00 | mCAF1 | CXCL14 |
| 33 | 0.00E+00 | 2.225770084 | 0.665 | 0.281 | 0.00E+00 | mCAF1 | PLAT |
| 34 | 0.00E+00 | 1.997004868 | 0.76 | 0.363 | 0.00E+00 | mCAF1 | PDGFRA |
| 35 | 0.00E+00 | 1.424280957 | 0.884 | 0.571 | 0.00E+00 | mCAF1 | TMEM176B |
| 36 | 6.98E-296 | 1.700946064 | 0.408 | 0.015 | 2.30E-291 | mCAF1 | NSG1 |
| 37 | 1.21E-280 | 1.432675952 | 0.734 | 0.359 | 4.00E-276 | mCAF1 | VCAN |
| 38 | 9.85E-276 | 1.925654959 | 0.392 | 0.018 | 3.24E-271 | mCAF1 | ENHO |
| 39 | 2.87E-274 | 1.521324315 | 0.438 | 0.053 | 9.45E-270 | mCAF1 | PDGFD |
| 40 | 2.33E-260 | 2.711094139 | 0.458 | 0.07 | 7.66E-256 | mCAF1 | CCL11 |
| 41 | 1.72E-259 | 1.367381755 | 0.388 | 0.023 | 5.64E-255 | mCAF1 | HSD17B2 |
| 42 | 2.56E-257 | 1.989175362 | 0.624 | 0.272 | 8.41E-253 | mCAF1 | F3 |
| 43 | 4.88E-249 | 1.199876762 | 0.728 | 0.401 | 1.60E-244 | mCAF1 | EMILIN1 |
| 44 | 2.38E-247 | 1.163196522 | 0.821 | 0.643 | 7.84E-243 | mCAF1 | NBL1 |
| 45 | 1.19E-226 | 1.324738128 | 0.455 | 0.103 | 3.93E-222 | mCAF1 | EMID1 |
| 46 | 3.22E-213 | 1.193714037 | 0.343 | 0.027 | 1.06E-208 | mCAF1 | WNT4 |
| 47 | 1.18E-204 | 1.687585639 | 0.418 | 0.084 | 3.90E-200 | mCAF1 | AGT |
| 48 | 4.37E-200 | 1.480722926 | 0.551 | 0.23 | 1.44E-195 | mCAF1 | BMP4 |
| 49 | 4.94E-194 | 1.152872139 | 0.64 | 0.322 | 1.63E-189 | mCAF1 | PDPN |
| 50 | 1.57E-181 | 1.559965192 | 0.648 | 0.301 | 5.15E-177 | mCAF1 | APOD |
| 51 | 1.397E-157 | 1.36200839 | 0.226 | 0.001 | 4.5948E-153 | mCAF1 | VSTM2A |
| 52 | 2.9919E-140 | 1.657156613 | 0.763 | 0.567 | 9.8407E-136 | mCAF1 | CTGF |
| 53 | 5.52E-120 | 1.307322392 | 0.784 | 0.627 | 1.82E-115 | mCAF1 | IGFBP5 |
| 54 | 7.72E-119 | 1.566610384 | 0.396 | 0.139 | 2.54E-114 | mCAF1 | IGFBP3 |
| 55 | 2.34E-107 | 1.265858335 | 0.462 | 0.207 | 7.71E-103 | mCAF1 | PTGDS |
| 56 | 4.53E-86 | 1.518082044 | 0.162 | 0.017 | 1.49E-81 | mCAF1 | PPAP2B |
| 57 | 1.10E-65 | 2.778926523 | 0.131 | 0.015 | 3.63E-61 | mCAF1 | CST1 |

| | | | | | | | |
|---|---|---|---|---|---|---|---|
| 58 | 8.80E-63 | 1.276709045 | 0.198 | 0.056 | 2.89E-58 | mCAF1 | GNB2L1 |
| 59 | 2.66E-62 | 1.265479932 | 0.376 | 0.232 | 8.75E-58 | mCAF1 | ABCA8 |
| 60 | 4.99E-62 | 1.205739754 | 0.23 | 0.083 | 1.64E-57 | mCAF1 | DEFB1 |
| 61 | 0.00E+00 | 4.910943583 | 0.602 | 0.042 | 0.00E+00 | mCAF2 | PLA2G2A |
| 62 | 0.00E+00 | 4.211962903 | 0.821 | 0.285 | 0.00E+00 | mCAF2 | CLEC3B |
| 63 | 0.00E+00 | 4.099724233 | 0.963 | 0.128 | 0.00E+00 | mCAF2 | SFRP2 |
| 64 | 0.00E+00 | 4.025015818 | 0.996 | 0.513 | 0.00E+00 | mCAF2 | CFD |
| 65 | 0.00E+00 | 3.884763518 | 0.955 | 0.169 | 0.00E+00 | mCAF2 | CLU |
| 66 | 0.00E+00 | 3.719667277 | 0.534 | 0.008 | 0.00E+00 | mCAF2 | MYOC |
| 67 | 0.00E+00 | 3.680147392 | 0.72 | 0.022 | 0.00E+00 | mCAF2 | PI16 |
| 68 | 0.00E+00 | 3.580375457 | 0.896 | 0.076 | 0.00E+00 | mCAF2 | SFRP1 |
| 69 | 0.00E+00 | 3.464818879 | 0.853 | 0.075 | 0.00E+00 | mCAF2 | MFAP5 |
| 70 | 0.00E+00 | 3.37820759 | 0.998 | 0.527 | 0.00E+00 | mCAF2 | MGP |
| 71 | 0.00E+00 | 3.301948756 | 0.993 | 0.815 | 0.00E+00 | mCAF2 | GSN |
| 72 | 0.00E+00 | 3.257483925 | 0.972 | 0.536 | 0.00E+00 | mCAF2 | IGFBP6 |
| 73 | 0.00E+00 | 2.986453403 | 0.693 | 0.072 | 0.00E+00 | mCAF2 | SLPI |
| 74 | 0.00E+00 | 2.533838399 | 0.979 | 0.436 | 0.00E+00 | mCAF2 | CCDC80 |
| 75 | 0.00E+00 | 2.477989504 | 0.877 | 0.119 | 0.00E+00 | mCAF2 | FBLN2 |
| 76 | 0.00E+00 | 2.472702664 | 0.657 | 0.021 | 0.00E+00 | mCAF2 | WISP2 |
| 77 | 0.00E+00 | 2.441643049 | 0.994 | 0.589 | 0.00E+00 | mCAF2 | FBLN1 |
| 78 | 0.00E+00 | 2.321624828 | 0.952 | 0.344 | 0.00E+00 | mCAF2 | EFEMP1 |
| 79 | 0.00E+00 | 2.304364246 | 0.985 | 0.598 | 0.00E+00 | mCAF2 | PLAC9 |
| 80 | 0.00E+00 | 2.275517604 | 0.895 | 0.2 | 0.00E+00 | mCAF2 | ADH1B |
| 81 | 0.00E+00 | 2.219557335 | 0.879 | 0.072 | 0.00E+00 | mCAF2 | OGN |
| 82 | 0.00E+00 | 2.19052851 | 0.948 | 0.3 | 0.00E+00 | mCAF2 | C3 |
| 83 | 0.00E+00 | 2.164730984 | 0.86 | 0.163 | 0.00E+00 | mCAF2 | GPNMB |
| 84 | 0.00E+00 | 2.098996657 | 0.999 | 0.846 | 0.00E+00 | mCAF2 | DCN |
| 85 | 0.00E+00 | 2.094336385 | 0.88 | 0.164 | 0.00E+00 | mCAF2 | MGST1 |
| 86 | 0.00E+00 | 2.013554221 | 0.879 | 0.17 | 0.00E+00 | mCAF2 | TNXB |
| 87 | 0.00E+00 | 1.875930058 | 0.81 | 0.105 | 0.00E+00 | mCAF2 | PRELP |
| 88 | 0.00E+00 | 1.873462376 | 1 | 0.932 | 0.00E+00 | mCAF2 | CST3 |
| 89 | 3.23E-212 | 2.204047303 | 0.559 | 0.156 | 1.06E-207 | mCAF2 | RARRES1 |
| 90 | 4.35E-12 | 1.869310043 | 0.331 | 0.219 | 1.43E-07 | mCAF2 | IGKV3-20 |
| 91 | 2.17E-232 | 4.101836024 | 0.696 | 0.059 | 7.15E-228 | vCAF1 | HHIP |
| 92 | 4.31E-227 | 2.399023142 | 0.596 | 0.041 | 1.42E-222 | vCAF1 | NPNT |
| 93 | 2.61E-219 | 3.434536518 | 0.918 | 0.12 | 8.58E-215 | vCAF1 | MYH11 |
| 94 | 5.84E-212 | 2.261538613 | 0.684 | 0.062 | 1.92E-207 | vCAF1 | KCNMB1 |
| 95 | 3.79E-205 | 4.537400912 | 0.924 | 0.145 | 1.25E-200 | vCAF1 | ACTG2 |
| 96 | 1.77E-152 | 3.499798572 | 0.368 | 0.022 | 5.82E-148 | vCAF1 | DES |
| 97 | 3.11E-145 | 1.86204847 | 0.468 | 0.04 | 1.02E-140 | vCAF1 | WFDC2 |
| 98 | 8.47E-138 | 2.720155599 | 0.778 | 0.136 | 2.79E-133 | vCAF1 | CNN1 |
| 99 | 7.80E-105 | 1.880661306 | 0.509 | 0.07 | 2.57E-100 | vCAF1 | COL23A1 |
| 100 | 1.08E-104 | 2.87878206 | 0.947 | 0.395 | 3.55E-100 | vCAF1 | MYLK |
| 101 | 7.26437E-96 | 1.905174582 | 0.632 | 0.122 | 2.38932E-91 | vCAF1 | SORBS1 |

| | | | | | | | |
|---|---|---|---|---|---|---|---|
| 102 | 1.07209E-95 | 2.799176675 | 0.988 | 0.539 | 3.5262E-91 | vCAF1 | ACTA2 |
| 103 | 6.12524E-94 | 2.156447179 | 0.795 | 0.221 | 2.01465E-89 | vCAF1 | LMOD1 |
| 104 | 5.34819E-91 | 2.84938805 | 0.743 | 0.205 | 1.75907E-86 | vCAF1 | CKB |
| 105 | 2.00318E-87 | 2.538116267 | 1 | 0.594 | 6.58866E-83 | vCAF1 | TAGLN |
| 106 | 7.19577E-86 | 2.587516751 | 0.971 | 0.594 | 2.36676E-81 | vCAF1 | FLNA |
| 107 | 5.12336E-82 | 2.304828613 | 0.994 | 0.752 | 1.68513E-77 | vCAF1 | TPM1 |
| 108 | 1.49629E-74 | 1.73136944 | 0.988 | 0.953 | 4.92145E-70 | vCAF1 | MYL6 |
| 109 | 1.99468E-74 | 2.129322445 | 0.982 | 0.806 | 6.5607E-70 | vCAF1 | TPM2 |
| 110 | 2.87918E-73 | 2.279484387 | 0.883 | 0.445 | 9.4699E-69 | vCAF1 | PDLIM3 |
| 111 | 1.46625E-68 | 1.940439569 | 0.895 | 0.446 | 4.82264E-64 | vCAF1 | CSRP1 |
| 112 | 1.01073E-66 | 1.883018885 | 0.988 | 0.808 | 3.32439E-62 | vCAF1 | MYL9 |
| 113 | 1.13751E-65 | 2.204371948 | 0.83 | 0.388 | 3.74138E-61 | vCAF1 | LTBP1 |
| 114 | 2.86486E-60 | 1.734525217 | 0.895 | 0.475 | 9.42282E-56 | vCAF1 | ACTN1 |
| 115 | 2.67242E-57 | 1.686823546 | 0.76 | 0.284 | 8.78986E-53 | vCAF1 | WFDC1 |
| 116 | 1.71001E-56 | 1.763156425 | 0.895 | 0.53 | 5.6244E-52 | vCAF1 | LPP |
| 117 | 2.02811E-54 | 1.717620484 | 0.947 | 0.8 | 6.67066E-50 | vCAF1 | NDUFA4 |
| 118 | 5.63716E-52 | 1.674123874 | 0.784 | 0.346 | 1.85412E-47 | vCAF1 | SYNPO2 |
| 119 | 1.87503E-45 | 1.679938258 | 0.889 | 0.638 | 6.16717E-41 | vCAF1 | FHL1 |
| 120 | 9.0115E-44 | 1.680793988 | 0.637 | 0.256 | 2.96397E-39 | vCAF1 | SMTN |
| 121 | 0 | 4.388143739 | 0.923 | 0.166 | 0 | vCAF2 | RGS5 |
| 122 | 0 | 3.289196498 | 0.86 | 0.133 | 0 | vCAF2 | NDUFA4L2 |
| 123 | 0 | 2.940854234 | 0.595 | 0.018 | 0 | vCAF2 | CD36 |
| 124 | 0 | 2.749179202 | 0.681 | 0.026 | 0 | vCAF2 | COX4I2 |
| 125 | 0 | 2.587606205 | 0.764 | 0.118 | 0 | vCAF2 | ARHGDIB |
| 126 | 0 | 2.470195123 | 0.609 | 0.034 | 0 | vCAF2 | GJA4 |
| 127 | 0 | 2.404632456 | 0.568 | 0.012 | 0 | vCAF2 | HIGD1B |
| 128 | 0 | 2.223228369 | 0.691 | 0.121 | 0 | vCAF2 | NOTCH3 |
| 129 | 0 | 1.871689462 | 0.472 | 0.045 | 0 | vCAF2 | STEAP4 |
| 130 | 0 | 1.864017484 | 0.995 | 0.931 | 0 | vCAF2 | IGFBP7 |
| 131 | 0 | 1.790822789 | 0.578 | 0.067 | 0 | vCAF2 | TINAGL1 |
| 132 | 0 | 1.711879409 | 0.508 | 0.051 | 0 | vCAF2 | FAM162B |
| 133 | 3.7151E-299 | 2.081770305 | 0.689 | 0.169 | 1.2219E-294 | vCAF2 | CSRP2 |
| 134 | 3.8489E-297 | 1.811918271 | 0.601 | 0.13 | 1.266E-292 | vCAF2 | 45173 |
| 135 | 1.7278E-271 | 1.735185306 | 0.622 | 0.141 | 5.6829E-267 | vCAF2 | MCAM |
| 136 | 1.4891E-267 | 2.150909097 | 0.693 | 0.212 | 4.8977E-263 | vCAF2 | IGFBP2 |
| 137 | 8.7571E-243 | 1.679240274 | 0.599 | 0.17 | 2.8803E-238 | vCAF2 | MYO1B |
| 138 | 1.55E-219 | 2.36502917 | 0.78 | 0.435 | 5.10E-215 | vCAF2 | COL4A1 |
| 139 | 1.94E-192 | 1.953945326 | 0.807 | 0.55 | 6.36E-188 | vCAF2 | COL4A2 |
| 140 | 4.37E-184 | 1.876176104 | 0.536 | 0.149 | 1.44E-179 | vCAF2 | FABP5 |
| 141 | 2.87E-178 | 1.90657895 | 0.666 | 0.284 | 9.44E-174 | vCAF2 | FRZB |
| 142 | 6.81E-178 | 1.571504496 | 0.646 | 0.276 | 2.24E-173 | vCAF2 | EPAS1 |
| 143 | 1.27E-174 | 1.737934853 | 0.701 | 0.343 | 4.17E-170 | vCAF2 | MAP1B |
| 144 | 1.19E-169 | 1.829926633 | 0.79 | 0.566 | 3.91E-165 | vCAF2 | COL18A1 |
| 145 | 7.16E-164 | 1.673107705 | 0.702 | 0.396 | 2.35E-159 | vCAF2 | PDGFRB |

| | | | | | | | |
|---|---|---|---|---|---|---|---|
| 146 | 1.42E-107 | 2.540857718 | 0.316 | 0.079 | 4.67E-103 | vCAF2 | FABP4 |
| 147 | 1.66E-74 | 1.762118758 | 0.367 | 0.136 | 5.45E-70 | vCAF2 | FP236383.1 |
| 148 | 3.66E-64 | 1.833913849 | 0.29 | 0.104 | 1.20E-59 | vCAF2 | RPL41P1 |
| 149 | 4.06E-10 | 6.162458136 | 0.117 | 0.064 | 1.34E-05 | vCAF2 | IGHV7-4-1 |
| 150 | 1.64E-03 | 5.399168542 | 0.132 | 0.104 | 1.00E+00 | vCAF2 | IGLV3-25 |
| 151 | 0.00E+00 | 3.37444987 | 0.928 | 0.09 | 0.00E+00 | vCAF3 | MYH11 |
| 152 | 0.00E+00 | 2.866378007 | 0.726 | 0.057 | 0.00E+00 | vCAF3 | SORBS2 |
| 153 | 0.00E+00 | 2.627532889 | 0.632 | 0.06 | 0.00E+00 | vCAF3 | SNCG |
| 154 | 0.00E+00 | 2.304181655 | 0.501 | 0.006 | 0.00E+00 | vCAF3 | RERGL |
| 155 | 0.00E+00 | 2.181145698 | 0.642 | 0.023 | 0.00E+00 | vCAF3 | PLN |
| 156 | 3.68E-305 | 2.285928271 | 0.675 | 0.082 | 1.21E-300 | vCAF3 | BCAM |
| 157 | 2.68E-300 | 1.803625999 | 0.775 | 0.1 | 8.81E-296 | vCAF3 | TINAGL1 |
| 158 | 1.04E-280 | 2.374184375 | 0.928 | 0.2 | 3.41E-276 | vCAF3 | CSRP2 |
| 159 | 1.06E-266 | 2.270337894 | 0.647 | 0.086 | 3.48E-262 | vCAF3 | C2orf40 |
| 160 | 4.04E-229 | 3.197217897 | 0.987 | 0.444 | 1.33E-224 | vCAF3 | ADIRF |
| 161 | 3.27E-223 | 3.360043679 | 0.997 | 0.579 | 1.07E-218 | vCAF3 | TAGLN |
| 162 | 2.27E-215 | 2.119978354 | 0.673 | 0.12 | 7.45E-211 | vCAF3 | CNN1 |
| 163 | 1.01E-200 | 2.766418896 | 1 | 0.801 | 3.31E-196 | vCAF3 | MYL9 |
| 164 | 3.69E-196 | 2.969202867 | 0.982 | 0.523 | 1.22E-191 | vCAF3 | ACTA2 |
| 165 | 1.51E-182 | 2.349685746 | 0.985 | 0.804 | 4.97E-178 | vCAF3 | DSTN |
| 166 | 1.60E-149 | 2.301209178 | 0.987 | 0.8 | 5.28E-145 | vCAF3 | TPM2 |
| 167 | 2.25E-136 | 2.30404034 | 0.916 | 0.51 | 7.41E-132 | vCAF3 | CRIP1 |
| 168 | 5.80E-134 | 2.032476935 | 0.867 | 0.432 | 1.91E-129 | vCAF3 | CSRP1 |
| 169 | 7.10E-132 | 2.076129206 | 0.742 | 0.275 | 2.34E-127 | vCAF3 | PCSK7 |
| 170 | 3.15E-131 | 1.815206093 | 0.887 | 0.456 | 1.04E-126 | vCAF3 | CRIP2 |
| 171 | 1.30E-122 | 2.643865098 | 0.816 | 0.37 | 4.28E-118 | vCAF3 | MT1M |
| 172 | 2.99E-116 | 1.802849275 | 0.88 | 0.431 | 9.82E-112 | vCAF3 | PPP1R14A |
| 173 | 7.90E-113 | 1.819691821 | 0.903 | 0.585 | 2.60E-108 | vCAF3 | FLNA |
| 174 | 7.35E-92 | 2.244452287 | 0.852 | 0.54 | 2.42E-87 | vCAF3 | GADD45B |
| 175 | 1.24E-53 | 1.935498246 | 0.772 | 0.511 | 4.08E-49 | vCAF3 | MT1X |
| 176 | 1.70E-49 | 1.861025408 | 0.478 | 0.193 | 5.59E-45 | vCAF3 | MT1A |
| 177 | 9.19E-48 | 1.850318746 | 0.777 | 0.53 | 3.02E-43 | vCAF3 | CDKN1A |
| 178 | 5.32E-30 | 2.513657276 | 0.12 | 0.023 | 1.75E-25 | vCAF3 | CCL21 |
| 179 | 1.06E-21 | 2.689022188 | 0.123 | 0.031 | 3.48E-17 | vCAF3 | CCL19 |
| 180 | 6.32E-12 | 2.035760135 | 0.194 | 0.095 | 2.08E-07 | vCAF3 | CH25H |

**Supplementary Table S7 Multilabel immunofluorescence staining antibody**

| Panel | Primary Antibodies | dilution | Secondary Antibodies | dilution |
|---|---|---|---|---|
| Figure 4K | TIGIT (ab243903, Abcam, UK) | 1:800 | Polymer HRP-anti-Mouse/Rabbit IgG (10013001010, Panovue, China) | 1: 3 |
| | CD8A-CST70306 | 1:100 | | |
| | Nectin 2 (ab233384, Abcam, UK) | 1:150 | | |
| | CD4 (ZM0418, Zsgb-Bio, China) | 1:200 | | |
| | Pan-CK (CST4545, CST, USA) | 1:500 | | |

| | | | | |
|---|---|---|---|---|
| Figure 5G | SPP1 (ab214050, Abcam, UK) | 1:1000 | | |
| | VEGFA (66828-1-Ig, proteintech, USA) | 1:300 | | |
| | CD68 (CST76437, CST, USA) | 1:500 | | |
| | TREM2 (ab223684, Abcam, UK) | 1:300 | Donkey anti-Goat IgG H&L (HRP) (ab6885, Abcam, UK) | 1: 800 |
